# Multiple *alr* genes exhibit allorecognition-associated variation in the colonial cnidarian *Hydractinia*

**DOI:** 10.1101/2022.03.25.485889

**Authors:** Henry Rodriguez-Valbuena, Andrea Gonzalez-Muñoz, Luis F. Cadavid

## Abstract

The genetics of allorecognition has been determined from inbred lines of *Hydractinia symbiolongicarpus*, in which genetic control is attributed mainly to the highly polymorphic loci *allorecognition 1 (alr1)* and *allorecognition 2 (alr2)* located within the Allorecognition Complex (ARC). While allelic variation at *alr1* and *alr2* can predict the phenotypes in inbred lines, these two loci do not entirely predict the allorecognition phenotypes in wild-type colonies and their progeny, suggesting the presence of additional uncharacterized genes that improve the prediction of these phenotypes. Comparative genomics analyses were used to identify coding sequence differences in assembled chromosomal intervals of the ARC and genomic scaffold sequences between two incompatible *H. symbiolongicarpus* siblings from a backcross population. New IgSF-like genes are reported for the ARC, five of these genes are closely related to the *alr1* and *alr2* genes, suggesting the presence of multiple *alr-type* genes within this complex. Cloning evidence revealed that the allelic polymorphism of eight Ig-SF-like genes is associated with allorecognition phenotypes in a backcross population of *H. symbiolongicarpus*. Remarkably, alternative splicing was found as a mechanism that contributes to the functional variability of these genes by changing putative activating receptors to inhibitory receptors, or generating secreted isoforms of allorecognition proteins. Our findings demonstrate that allorecognition in *H. symbiolongicarpus* is a multigenic phenomenon controlled by genetic variation in at least eight genes in the ARC complex, as well as putative uncharacterized variation outside of this region.

## INTRODUCTION

Benthic colonial invertebrates often encounter conspecifics while growing on their hard substrata, with the encounters typically resulting in either fusion or rejection (Rosengarten and Nicotra 2011). This allorecognition phenomenon determines the outcomes of colony competition for habitable space, and it is widely distributed among diverse animal groups, including sponges, bryozoans, corals, anemones, hydrozoans, and ascidians (Taketa and De Tomaso 2015; Grice et al. 2017; Buckley and Dooley 2022; Nicotra 2022). Allorecognition phenomena has been studied extensively in the colonial hydroid *Hydractinia* (Cnidaria; Hydrozoa) (Nicotra 2022) and, together with the ascidian *Botryllus schlosseri*, they are established model systems for invertebrate allorecognition (Taketa and De Tomaso 2015). *Hydractinia* colonies grow on gastropod shells occupied by pagurid hermit crabs and are composed of polyps communicated through endodermal canals, both of which are sandwiched into two ectodermal epithelial layers (Buss and Yund 1989). Physical contact between *Hydractinia* colonies results in one of three allorecognition phenotypes: *i*) fusion, where colonies adhere and merge to form a permanent chimera sharing a functional gastrovascular system, *ii*) rejection, where colonies fail to adhere and experience a massive migration of stinging cells (cnidocytes) towards the contact zone, which upon discharge cause extensive necrosis to the allogenic tissue; and *iii*) transitory fusion, where colonies first fuse, only to develop a rejection response after a few days (Hauenschild 1954; Shenk and Buss 1991; Powell et al. 2007).

Analyses using *Hydractinia* inbred lines have shown that allorecognition is controlled in a single chromosomal interval, known as the Allorecognition Complex (ARC) (Mokady and Buss 1996; Cadavid et al. 2004). The ARC contains two linked, polymorphic, and codominantly expressed loci, *allorecognition (alr) 1* and *2*, which together determine the allorecognition outcomes, such that colonies fuse if the share one or two alleles at both loci, reject if they share no alleles at either locus, and display transitory fusion if they share at least one allele at a single locus (Cadavid et al. 2004). The ARC region was mapped at a 2.0 cM resolution (Cadavid et al. 2004; Powell et al. 2007) and the allorecognition genes *alr1* and *alr2* were identified by positional cloning (Nicotra et al. 2009; Rosa et al. 2010) *Alr1* gene encodes a transmembrane protein with two variable extracellular immunoglobulin (Ig)- like domains and a cytoplasmic region having an immunoreceptor tyrosine-based activation motif (ITAM)-like motif (Rosa et al. 2010). In addition, nine other genes encoding Ig domain-containing transmembrane proteins were found in the vicinity of the *alr1* gene (Rosa et al. 2010). *Alr2* encodes a transmembrane protein with three variable Ig-like domains and an immunoreceptor tyrosine-based inhibitory motif (ITIM)-like in the cytoplasmic tail (Nicotra et al. 2009). Additionally, two *alr2-like* pseudogenes were found in close proximity to *alr2* (Nicotra et al. 2009), displaying high sequence similarity to the *alr2* exons encoding the extracellular Ig-like domains (Rosengarten et al. 2011).

The *alr1* and *alr2* genes have proven to be highly polymorphic (Nicotra et al. 2009; Rosa et al. 2010; Gloria-Soria et al. 2012). For instance, 180 unique *alr2* amino acid sequences have been reported in 239 *Hydractinia* colonies (Gloria-Soria et al. 2012). These *alr2* alleles showed higher variability in the first Ig-domain (exon 2) than the second and third Ig-domains, with several positions under positive selection (Rosengarten et al. 2011). The *alr2* alleles were classified into two allele types (type I and II) based on sequence differences between exons 5, 6, and 7, encoding the stem region, the transmembrane domain, and the cytoplasmic region, respectively. Furthermore, chimeric *alr2* alleles between type I and II alleles have been detected (Rosengarten et al. 2011). Remarkably, haplotypes with two or three *alr2* pseudogenes were identified, indicating haplotypic variation in this genomic region (Rosengarten et al. 2011). Recently, it has been shown that functional diversity of the *alr2* gene is generated through a mechanism of point mutations, by which successive single nucleotide changes create new *alr*-binding specificities, with the feature that intermediate allele stages have broader *alr*-binding specificities (Huene et al. 2021).

The variation in the *alr1* gene has been studied to a lesser than that of *alr2*. The amino acid sequence identity among Alr1 alleles in the extracellular region was found to be nearly 30%, whereas in the cytoplasmic region it was 94% (Rosa et al. 2010). In addition, splicing variants were detected involving exon 4 (coding the stem region) of *alr1*, resulting in shorter versions of the protein (Rosa et al. 2010). Moreover, Karadge et al. (2015) established that the Alr1 and Alr2 proteins interact through a homophilic *trans* interaction based on *in vitro* expression studies. Furthermore, this interaction occurs between the same *alr*-allele or close variants, indicating an allele-specific interaction (Karadge et al. 2015).

The known extension of the ARC includes five reported regions sequenced from 27 Bacterial Artificial Chromosome (BAC) clones, which include the *alr1* and *alr2* chromosomal intervals and three uncharacterized intervals (Nicotra et al. 2009; Rosa et al. 2010). However, the full extension of the ARC has not been fully sequenced and the plausible existence of additional Ig-like genes and/or other variable allodeterminants within this interval represents a topic of active investigation. Unexplored variation within the ARC could explain deviations from the two-locus model often observed in *Hydractinia* wild-type colonies and, occasionally, in inbred lines (Rosa et al. 2010; Powell et al. 2011; Nicotra 2022). Indeed, Grosberg et al. (1996) proposed an alternative genetic model based on the frequency of allorecognition phenotypes of full-siblings and half-siblings, where allorecognition in *H. symbiolongicarpus* is controlled by three to seven loci with a low number of alleles per locus (approximately five to seven alleles) (Grosberg et al. 1996). This model has not predicted the polymorphism of *alr1* and *alr2* that were subsequently characterized (Nicotra et al. 2009; Rosa et al. 2010; Rosengarten et al. 2011; Gloria-Soria et al. 2012); however, it proposed the existence of additional allorecognition genes in *Hydractinia*. Moreover, it has been reported that interacting colonies sharing alleles at both *alr1* and *alr2* genes can display transitory fusion or rejection phenotypes, instead of the fusions predicted by the two-locus model (Rosa et al. 2010).

Here we analyzed the genomic variation at the ARC between two incompatible *H. symbiolongicarpus* siblings derived from a backcross population. We fully annotated the five ARC intervals previously sequenced and identified three new *alr1-like* genes, two new *alr2- like* genes, and an *Ig-like* gene. Complementary DNAs from these genes was sequenced, and their variation largely correlated with the allorecognition phenotypes in the backcross population. Furthermore, alternative splicing was detected as a common mechanism generating additional variability in *alr* genes. Our results indicate that allorecognition in *Hydractinia* is controlled by multiple *alr* genes located in the ARC, and that the genes vary substantially through point mutations, recombination, and alternative splicing.

## MATERIALS AND METHODS

### H. symbiolongicarpus culture, crosses, and fusibility assays

*H. symbiolongicarpus* colonies were collected in Woods Hole, MA (USA), and were grown on microscope glass slides in 30-liter aquaria with artificial seawater (Instant Ocean, US) at a relative density of 1.022 and an average temperature of 21°C, as detailed in (Blackstone and Buss 1991). Animals were fed with two-day-old *Artemia salina* nauplii, with water changes of 20% three times per week. Two wild-type individuals (HWB29 and HWB53 - ***H****ydractinia*, **W**oods Hole, **B**ogotá-) were crossed to obtain an F1 population, following the methodology described in (Cadavid et al. 2004). A female F1 individual (F1-8) was crossed with its male parent (HWB29) to generate a backcross (BC) population of 33 individuals. These animals were tested for fusibility among themselves and with their parental colonies using the colony assay (Müller 1964; Ivker 1972). To test the correlation between *alr* gene variability and fusibility phenotypes, two fusibility groups (A and B) were assembled from the BC population, where all individuals from the same group fused and rejected all individuals from the other group.

### Whole genome sequencing and de novo genome assembly

High-molecular weight DNA was extracted from two histo-incompatible sibling colonies, namely BC-15 (Group A) and BC-3 (Group B), using a modified phenol-chloroform protocol (Sanders et al. 2018). For each individual, 200 bp-insert paired end libraries and 3 Kb-insert mate pair ‘long jumping distance’ (LJD) libraries were sequenced on an Illumina HiSeq 2000 platform at EuroFins MWG Operon (USA). Sequencing depth was calculated by Lander and Waterman’s general equation (Lander and Waterman 1988) based on a read length of 100 bp and an estimated genome size of 514 Mb (Frank et al. 2020). Read quality control was assessed using FastQC (Andrews 2010) and reads with a quality score above 20 and minimum read length of 70 were retained using Cutadapt (Martin 2011). Scaffolded assemblies were generated *de novo* with ABySS v1.5.1 (Simpson et al. 2009) using the ‘rescaffolding’ option with *H. symbiolongicarpus* contigs assembled from RNA-Seq short read data (Zárate-Potes and Cadavid 2014; Zárate-Potes et al. 2019). Assembly statistics were calculated using QUAST (Gurevich et al. 2013). Raw sequencing data are available under NCBI BioProject ID PRJNA801058. The genome assemblies for BC-3 and BC-15 are available in Zenodo under DOI 10.5281/zenodo.5784056.

### Assembly of allorecognition complex chromosomal intervals

Five ARC intervals were assembled for individuals BC-3 and BC-15 by mapping the sequenced reads and scaffolds to five reference ARC intervals previously sequenced from a *H. symbiolongicarpus* individual (Rosa et al. 2010). To accomplish this, the reference ARC intervals were first assembled from 27 BAC clone sequences downloaded from GenBank (Supplementary Material, Table S1). An overlapping assembly approach employing BLASTN was used to extend the clones to the longest possible contigs. The five reference intervals corresponded to two genomic regions containing the reported *alr1* and *alr2* genes, respectively, and three uncharacterized regions in the ARC, which are designated here as Intervals 1, 2, and 3. Next, reads and assembled scaffolds for individuals BC-3 and BC-15 were separately mapped to the reference intervals using the Burrows-Wheeler Algorithm (*bwa*) software (Li and Durbin 2010) and the sequences were generated from the alignments using Samtools (Li et al. 2009). The final sequences for each interval per individual were generated from a consensus of both mapping results (i.e., reads and scaffold sequences).

### Gene prediction and annotation

Gene prediction was performed using the MAKER2 genome annotation pipeline (Holt and Yandell 2011) by integrating *ab-initio* predictions obtained with SNAP (Korf 2004) and AUGUSTUS (Stanke and Morgenstern 2005). Both prediction programs were trained using *H. symbiolongicarpus* mRNA evidence (Zárate-Potes and Cadavid 2014; Zárate-Potes et al. 2019) to generate *Hydractinia-*specific gene models for accurate gene prediction. SNAP was trained according to the MAKER2 pipeline tutorial (http://gmod.org/wiki/MAKER_Tutorial_2013) and AUGUSTUS was trained through the Web Augustus Training Server (Hoff and Stanke 2013) (http://bioinf.uni-greifswald.de/webaugustus/training/create). Gene prediction was performed on the assembled ARC sequences and genome scaffolds greater or equal to 10 Kb. Homology-based annotation of the predicted proteins was obtained by a protein BLAST search against RefSeq and UniprotKB/SwissProt databases. Functional annotation was assigned based on predicted protein architectures and functions retrieved from Interproscan searches (Hunter et al. 2012) (http://www.ebi.ac.uk/interpro/).

### Comparative analyses of coding sequences

Comparative analyses were done to identify the genetic variation between homologous coding sequences of BC-3 and BC-15 individuals. Additionally, the transcriptomes of two wild-type *H. symbiolongicarpus* individuals, namely HWB29 and HWB103, were included in these analyses. HWB29 is the male parent of the backcross population, while HWB103 is an unrelated wild-type individual. Transcriptome assemblies for these individuals are derived from previous studies in our laboratory (Zárate-Potes and Cadavid 2014; Zárate-Potes et al. 2019) and are available at Zenodo under DOI 10.5281/zenodo.5784056. Homologous proteins in the ARC intervals between individuals BC-3 and BC-15 were established based on synteny, while homologous proteins across BC-3, BC-15, HWB29 and HWB103 were retrieved by reciprocal best hits (RBH) (Moreno-Hagelsieb and Latimer 2008; Ward and Moreno-Hagelsieb 2014). Point mutations were identified first through pairwise BLASTP alignments between BC-3 and BC-15 predicted protein sequences. The selected variable proteins were later aligned to transcript sequences from HWB29 and HWB103 using MUSCLE (Edgar 2004). Genetic variation was quantified by estimating the pairwise p-distance (Nei and Kumar 2000) between homologous protein sequences using MEGA5.2 (Tamura et al. 2011).

The allorecognition genes were analyzed in two additional transcriptomes of *H. symbiolongicarpus*. The first one was published by (Sanders et al. 2014) (Accession Number GAWH00000000), and the second one is publicly available (https://research.nhgri.nih.gov/hydractinia/) and corresponds to a male wild-type individual (strain 291-10) that was sequenced by the *Hydractinia* Genome Project Team (Frank et al. 2020). Furthermore, allorecognition genes were analyzed in the publicly available preliminary draft genome of *H. symbiolongicarpus* that was Illumina-sequenced by the *Hydractinia* Genome Project Team (Frank et al. 2020) (https://research.nhgri.nih.gov/hydractinia/).

Phylogenetic relationships of the predicted Alr1-type and Alr2-type proteins were analyzed based on the sequences from colonies BC-3, BC-15, HWB29, HWB103, as well as reported alleles retrieved from the NCBI Nucleotide database. Phylogenetic analyses were based on protein alignments generated with ClustalW using default parameters. Maximum Likelihood (ML) trees were obtained with MEGA-X software (Kumar et al. 2018). For the Alr1-type proteins, the extracellular region spanning the second Ig-domain to the transmembrane was aligned and the analyses were inferred using the Jones-Taylor-Thornton (JTT) amino acid substitution model (Jones et al. 1992) with gamma substitution rates, partial deletion of missing data with 95% cutoff, and 100 Bootstrap replicates. The first Ig-domain region did not align well across the Alr1-type proteins due to its extremely high variability, and therefore, it was excluded from the alignment. For the Alr2-type proteins, the complete extracellular region spanning the Ig-domains to the transmembrane was aligned and the analyses were run using the WAG model (Whelan and Goldman 2001) with gamma substitution rates, partial deletion of missing data (95% cutoff), and 100 Bootstrap replicates. The best model for each dataset was selected using the best-fit model tool in MEGA-X (Kumar et al. 2018).

### Cloning and sequencing of allorecognition genes

Allorecognition genes showing variability between the incompatible siblings BC-3 and BC-15 were amplified in the parental individuals HWB29, HWB53, and F1-8, and in ten backcross individuals from fusibility groups A and B. Briefly, total RNA was extracted using the Trizol^TM^ reagent (Thermo Fisher, US) following the manufacturer’s protocol. cDNA synthesis was performed using the High-Capacity cDNA Reverse Transcription Kit (Thermo Fisher, US). Genes were amplified with the primers specified in Table S2 and Taq DNA polymerase (Thermo Fisher, US). Amplification was performed following an initial step at 95°C for 4 minutes, 30 cycles at 95°C for 40 seconds, 57°C for 35 seconds and 72°C for 2 minutes, and a final step at 72°C for 10 minutes. PCR products were purified with the GeneJET PCR Purification Kit (Thermo Fisher, US) and cloned into the pGEM®-T Easy Vector system (Promega, US). Recombinant colonies were evaluated by colony PCR using SP6 and T7 primers, and an amplification protocol that consisted of an initial step at 95°C for 3 minutes, 30 cycles at 95°C for 40 seconds, 50°C for 30 seconds, and 72°C for 2 minutes, and a final step at 72°C for 5 minutes. Ethanol-purified products were Sanger-sequenced by the sequencing service of the Instituto de Genética-Universidad Nacional de Colombia. The sequences were edited and analyzed in BioEdit (Hall 1999) and Geneious Prime® 2021.1.1 (https://www.geneious.com). A sequence was reported if at least three identical clones were found. If exceptions to this rule were included, they are explicitly described in the text with the number of clones. The cloned sequences were deposited in GenBank under accession numbers OM745700-OM745748.

## RESULTS

### Gene composition within the Allorecognition Complex

Five ARC chromosomal regions were assembled for siblings BC-3 and BC-15 based on reads and genomic scaffold alignments to the five reference intervals derived from BAC clones (Fig. 1). The assembled intervals spanned an average of 2.77 Mb in both individuals, and were designated as Alr1, Alr2, Interval 1, Interval 2, and Interval 3. The previously reported Alr2 region of 375,101 bp (Nicotra et al. 2009) (GenBank Accession EU219736) was extended to 529,880 bp (Fig. 1). This extension allowed predicting new genes flanking this ARC region. The Alr1 interval was approximately 1.2 Mb in length and contained 43 genes, the Alr2 interval covered approximately 0.52 Mb and had 24 genes, Interval 1 was 0.59 Mb and contained 33 genes, Interval 2 was 0.21 Mb and had 15 genes, and Interval 3 covered 0.21 Mb and contained 4 genes (Fig. 1 and Table S3). In addition, the Alr1 and Alr2 intervals were fully annotated for individual BC-15 (i.e., used as reference sibling due to fewer assembly gaps than BC-3) and the BAC-derived reference sequences (Fig. S1). Comparisons between these annotations showed that individual BC-15 had two putative functional genes similar to *alr2* gene (named here as *alr2B* and *alr2C* genes), which were in the same locations of the two reported *alr2* pseudogenes in the Alr2 reference interval (Fig. 1 and Fig. S1). Other differences between the BC-15 assembly and the reference intervals included the presence of an uncharacterized gene encoding a transmembrane protein in the Alr1 reference interval that was absent in individual BC-15, the presence of a malate dehydrogenase gene and a hypothetical gene in the Alr2 reference interval that were absent in the assembly for individual BC-15, and the presence of a zinc finger-encoding gene and an uncharacterized gene that were present in the BC-15 assembly but absent in the reference Alr2 interval (Fig. S1).

**Fig. 1.**
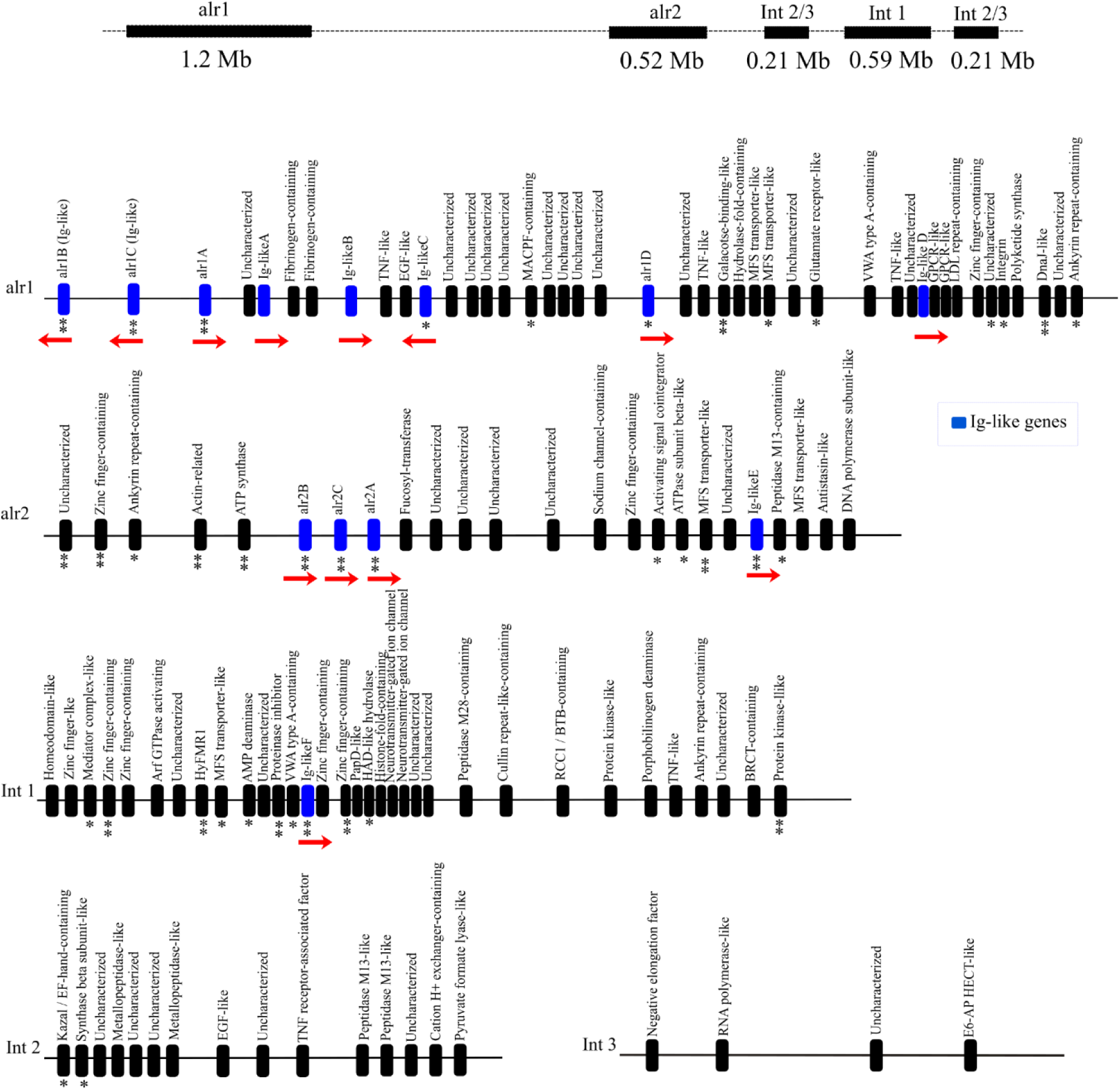
Gene content of the ARC intervals alr1, alr2, Interval 1, Interval 2, and Interval 3 assembled in *Hydractinia symbiolongicarpus* individuals BC-3 and BC-15. Top: Hypothetical relative location of the intervals within the ARC (based on Rosa et al. (2010)). Bottom: Relative location of predicted genes in each interval, identified by their predicted function. The letters A-F were arbitrarily assigned to the IgSF-like genes found in this study and do not correspond to the identification given to these genes by Rosa et al. (2010). The red arrows indicate the direction of transcription (right: sense 5’-3’, left, anti-sense 3’-5’). The number of asterisks represents the sequence variation between the encoded proteins in individuals BC-3 and BC-15, based on Nei’s p-distance: *** (p- distance ≥ 0.20), ** (0.02 ≤ p-distance < 0.20), * (p-distance < 0.02)

From the 119 genes predicted in the ARC region 52 were newly reported genes in Intervals 1, 2, and 3, and 88 of these genes (74%) were functionally annotated (Fig. 1, Table S4). Eleven newly reported IgSF-like genes (excluding the *alr1* and *alr2* genes) were found in the intervals Alr1, Alr2, and Interval 1 (Fig. 1, blue boxes). Among these, five genes (*alr1B*, *alr1C*, *alr1D*, *alr2B*, and *alr2C*) were similar to the *alr1* and *alr2* reference genes (which hereafter we refer to as *alr1A* and *alr2A*, respectively). Eight IgSF-like genes, namely *alr1A, alr1B*, *alr1C*, *alr1D*, *Ig-like A*, *Ig-like B*, *Ig-like C*, and *Ig-like D*, were annotated in the Alr1 interval. Although previous reports indicated the presence of ten IgSF-like genes in this interval (Rosa et al. 2010), it was not possible to identify them because the sequences were not made available for comparison. Therefore, we annotated the reported Alr1 interval (Table S5) and compared it with the annotations derived from individual BC-15 (Fig. S1). In the Alr2 interval, six out of nine genes from the reference interval were confirmed in the BC-15 assembly (Nicotra et al. 2009), namely *alr2A*, *ankyrin repeat-containing* gene, *actin-related* gene, *ATP synthase* gene, f*ucosyltransferase*, and an *uncharacterized gene* (Fig. S1). Overall, seven predicted genes or coding sequences were different between the reference intervals Alr1 and Alr2 and the assemblies for individuals BC-3 and BC-15 (Fig. S1).

### Sequence variation within the ARC

We analyzed the variability of the predicted ARC genes between individuals BC-3 and BC-15. Forty coding sequences showed nucleotide variation between these two individuals (Fig. 1) and their corresponding predicted proteins were classified into seven functional groups: 1) Immunity, recognition, and cellular adhesion; 2) Protein-protein interaction; 3) Proteolysis; 4) Enzymes; 5) Protease inhibitors; 6) Ion transport; and 7) Other unrelated functions (Fig. S2A). In addition, two variable genes that encoded proteins with unknown domains could not be functionally annotated. Remarkably, the most variable proteins were encoded in the Alr1 and Alr2 intervals, and in Interval 1, and comprised members of the Ig-like super family, including Alr receptors. Other variable proteins within the ARC included Major Facilitator Superfamily (MFS)-like proteins and ATP synthases, kinases and proteinase inhibitors, Zinc finger-containing proteins, and a DnaJ domain-containing protein. Although in the ascidian *B. schlosseri* the allorecognition HSP40-like protein is polymorphic and contains a DnaJ- and two Zinc-finger domains (Nydam et al. 2013), it is not homologous to the ones encoded in the *Hydractinia* ARC.

### Alr1 and Alr2 gene complexes

Thirty-four newly reported genes were found in the Alr1 interval (43 total genes), including 21 that were functionally annotated (Fig. 1). Within this region, three new IgSF-like genes were similar to the reported *alr1* gene (renamed here as *alr1A*) and were designed as *alr1B, alr1C*, and *alr1D* (Fig. 1). As previously reported, *alr1A* is composed of eight exons, where exon 1 encodes the signal peptide, exons 2 and 3 encode the first and second Ig-like domains, respectively, exon 4 encodes the stem region, exon 5 encodes the transmembrane (TM) domain, and exons 6, 7 and 8 encode the cytoplasmic region containing an ITAM-like motif (Rosa et al. 2010)(Fig. 2A). Interestingly, four alternative exons coding a new region for the *alr1A* gene were found downstream from exon 8 of the *alr1A* gene (Fig. 2A). The first alternative exon (exon 5’) was predicted to encode new TM domain whereas the other three alternative exons (exons 6’ to exon 8’) encoded a new cytoplasmic region (Fig. 2A). In addition, *alr1B*, *alr1C* and *alr1D* were composed of 6 exons, where exons 2 and 3 encoded the putative first and second Ig-like domains, exon 5 encodes the TM-domain, and exon 6 encoded the cytoplasmic region containing ITAM- or ITIM-like sequences (Fig. 2A). Interestingly, the cytoplasmic region of *alr1A* is encoded by three exons (the small exons 6 and 7, and the larger exon 8 with the ITAM-motif), while the cytoplasmic regions of *alr1B*, *alr1C* and *alr1D* genes are encoded by single exons that contains tyrosine-based motifs (Fig. 2A). Additionally, although the exon encoding the signal peptides of *alr1B* and *alr1C* genes were not identified in the available genomes, it was identified in the *alr1D* gene. Also, the first two exons of *alr1C* were identified only in the genome or transcriptome from strain 291-10 of the *Hydractinia* Genome Project (Frank et al. 2020) (Fig. 2A).

**Fig. 2.**
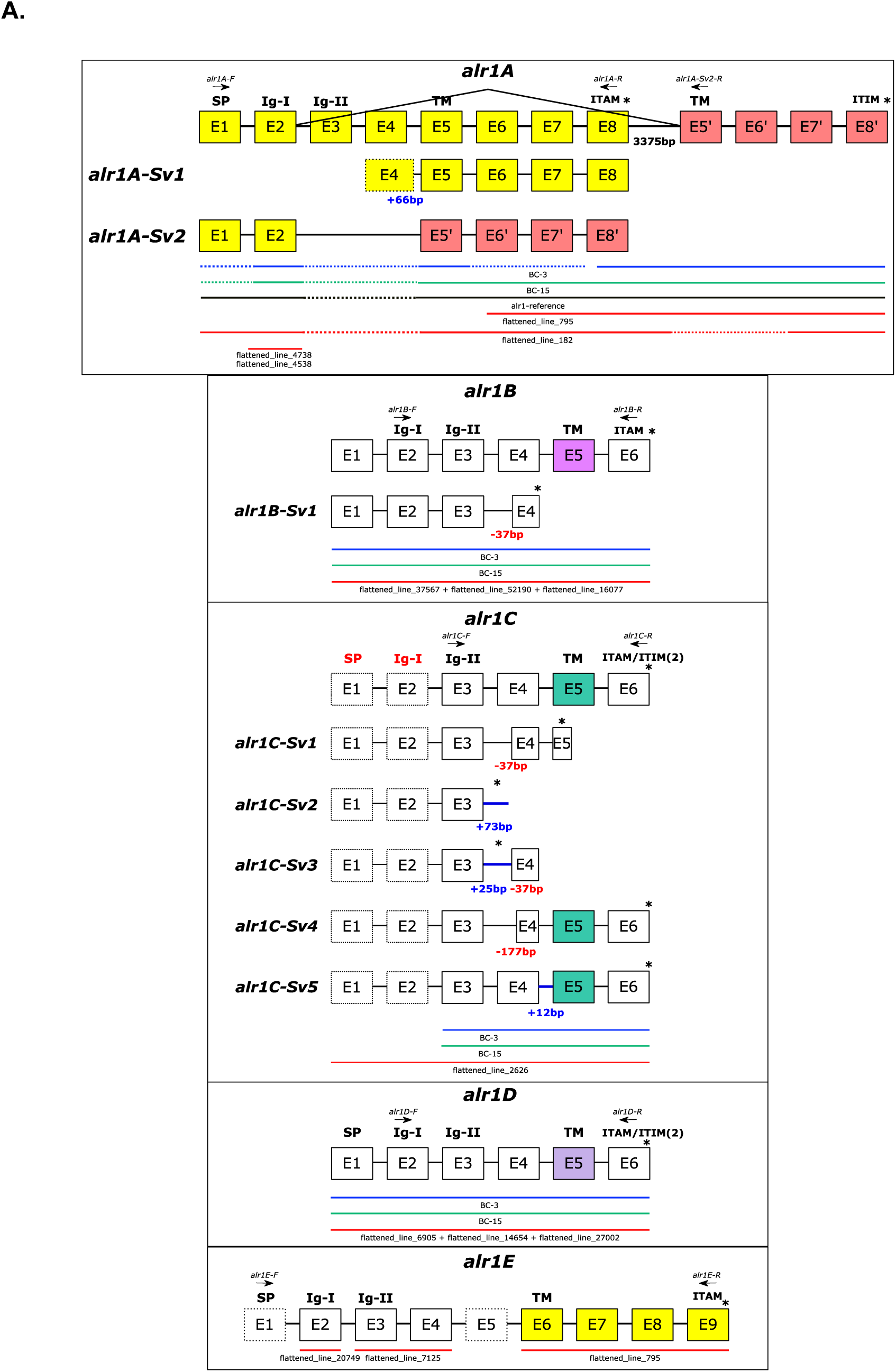
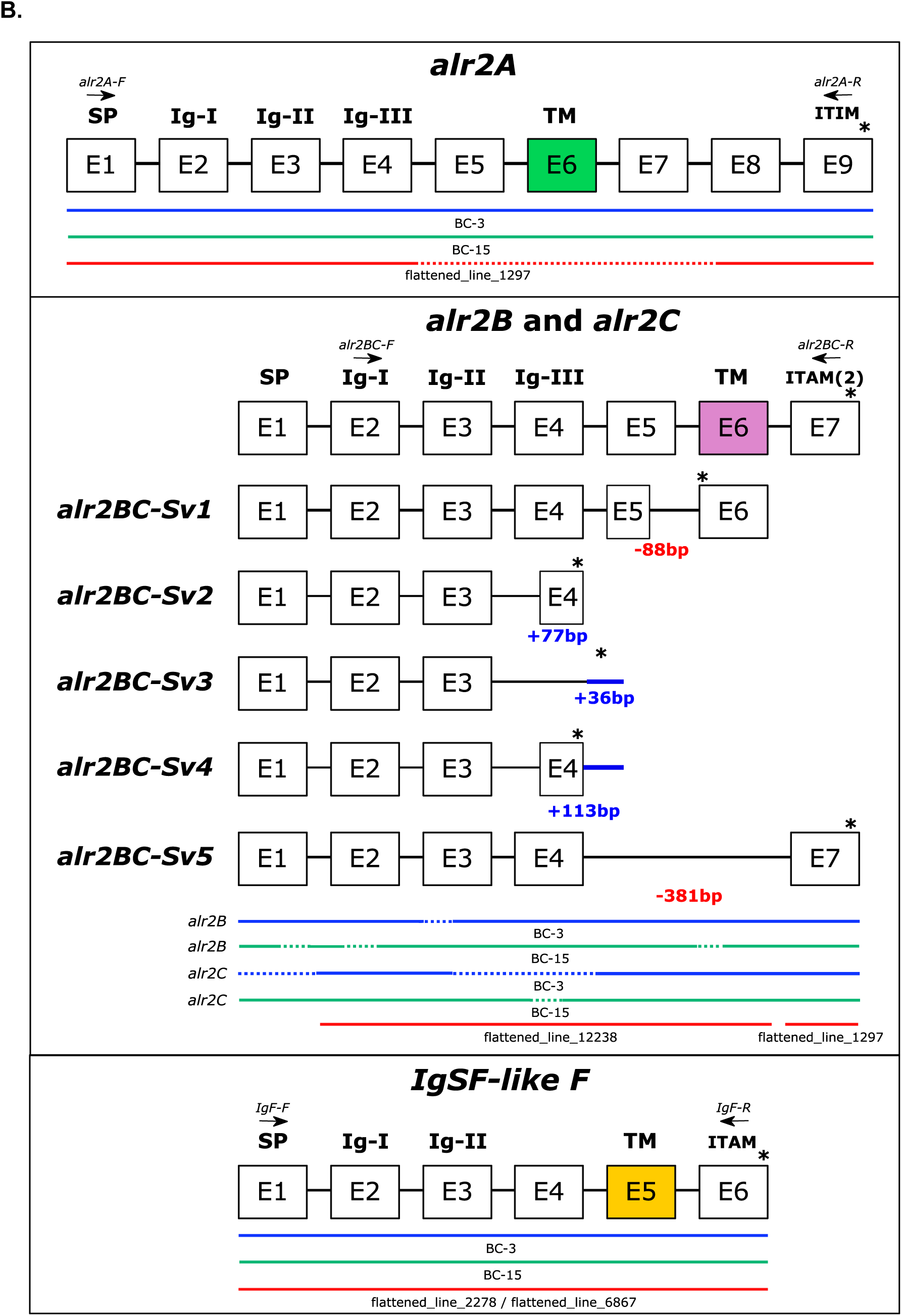
(A) Genomic structure of *alr1-type* genes. *alr1A* gene has eight exons (highlighted in yellow) and two splicing variants (Sv1 and Sv2). *alr1A-Sv1* is a splicing variant of the allele 2 found in our backcross population with a small insertion (66bp) before the TM domain. *alr1A-Sv2* uses the first two exons of the *alr1A* gene and four alternative exons (highlighted in red) downstream of exon 8. *alr1B* and *alr1C* genes showed splicing variants (Sv) that are represented below the genomic structure of the gene. The size of the insertions and deletions of splicing variants are indicated in blue and red colors, respectively. Exons 1 and 2 of the *alr1C* gene (dotted rectangles) were detected only in the genome of strain 291-10. Signal peptide and first Ig-domain of the *alr1C* gene (highlighted in red) that were identified at the transcriptomic level, could not be mapped to the genome. *alr1A* and *alr1E* genes shared the last four exons (highlighted in yellow). First five exons of the *alr1E* gene are unique for this molecule. Exons 1 and 5 of the *alr1E* gene were not detected in the genomic data (dotted squares). **(B) Genomic structure of *alr2-type* and *IgSF-F* genes.** *alr2BC* genes showed five splicing variants (Sv1-Sv5) that are described as splicing variants for the *alr1A*, *alr1B* and *alr1C* genes. The primers used in the cloning process are shown on the top of each gene. On the bottom of each gene, the genomic evidence from the individuals BC-3, BC-15, and wild-type strain 291-10 are represented in blue, green, and red lines, respectively, with the scaffolds’ names. Dotted lines indicate that those regions were not detected in the genomes. Exons that encode the signal peptide (SP), Immunoglobulin domains I, II and III (Ig-I, Ig-II, and Ig-III), transmembrane domain (TM), ITAM and ITIM motifs are shown above the exons. Stop codons are indicated by asterisks

On the other hand, 18 newly reported genes out of 24 genes were predicted in the Alr2 interval (Fig. 1). Thirteen of these genes were annotated by function, including three IgSF- like genes named here as *alr2B*, *alr2C*, and *IgSF-like E*. The first two displayed high sequence similarity to the exons encoding the extracellular region of *alr2A*, but differed in the exons encoding the stem, transmembrane, and cytoplasmic regions, as discussed later (Fig. S3). Furthermore, *alr2B* and *alr2C* shared sequence similarity and mapped to the genomic locations of the two previously reported *alr2* pseudogenes (Nicotra et al. 2009). Our gene predictions indicate that these are protein-coding genes in the BC-3 and BC-15 individuals because they show no evidence of premature stop codons (Fig. S1 and Fig. S3B). Indeed, these two genes were found in the transcriptomes from individuals HWB29 and HWB103 and were cloned and sequenced in backcross population (Fig. S3B and Fig. S14). Thus, *alr2A* has at least two related genes, namely *alr2B* and *alr2C*, that could have been pseudogenized in the inbred lines and might configure an instance of haplotype diversity within the ARC. Whether this haplotype diversity affects the allorecognition phenotypes remains to be investigated.

The *alr2B* and *alr2C* genes shared similar genomic structure with the *alr2A* gene (Fig. 2B). As reported before, the structure of *alr2A* is composed of nine exons, with exon 1 encoding the signal peptide, exons 2 - 4 the three Ig-like domains, exon 6 the TM domain, and exons 7, 8, 9 the cytoplasmic domain containing an ITIM-like motif (Nicotra et al. 2009). Meanwhile, the genomic structure of *alr2B* and *alr2C* genes comprises seven exons, where exons 1 to 5 encode the extracellular region similar to the *alr2A* gene. However, the cytoplasmic regions of *alr2B* and *alr2C* genes were encoded by a single exon (exon 7) containing two putative ITAM motifs (Fig. 2B and Fig. S14). Thus, *alr2B* and *alr2C* differ from *alr2A* gene in the number of exons encoding the cytoplasmic region, which was also observed when *alr1B*, *alr1C* and *alr1D* were compared with the *alr1A* gene (Fig. 2A). This dynamic in the cytoplasmic regions of *alr-type* genes could indicate that this region is constantly changing to generate activating receptors from inhibitory receptors (e.g., *alr2B* and *alr2C* genes from the *alr2A* gene), or vice versa.

### Variation of allorecognition genes in a segregating population

We evaluated the segregation of *alr* genes and their correlation to the allorecognition phenotypes in a backcross population (Fig. 3A). Two wild-type individuals HWB29 and HWB53 were mated to generate an F1 population of eight colonies. A female colony F1-8 was then mated back to its male parent HWB29 to generate a backcross population of 33 individuals. Fusibility assays were set among the colonies of this backcross population, including the parental individuals (Fig. 3B and Table S7). Thus, two fusibility groups were established where individuals belonging to the same group fused and individuals from different groups rejected each other (Fig. 3B). The fusibility group A included individuals BC- 15, BC-23, BC-36, BC-40, and BC-70, while fusibility group B included individuals BC-3, BC- 9, BC-22, BC-50, and BC-53 (Fig. 3A).

**Fig. 3.**
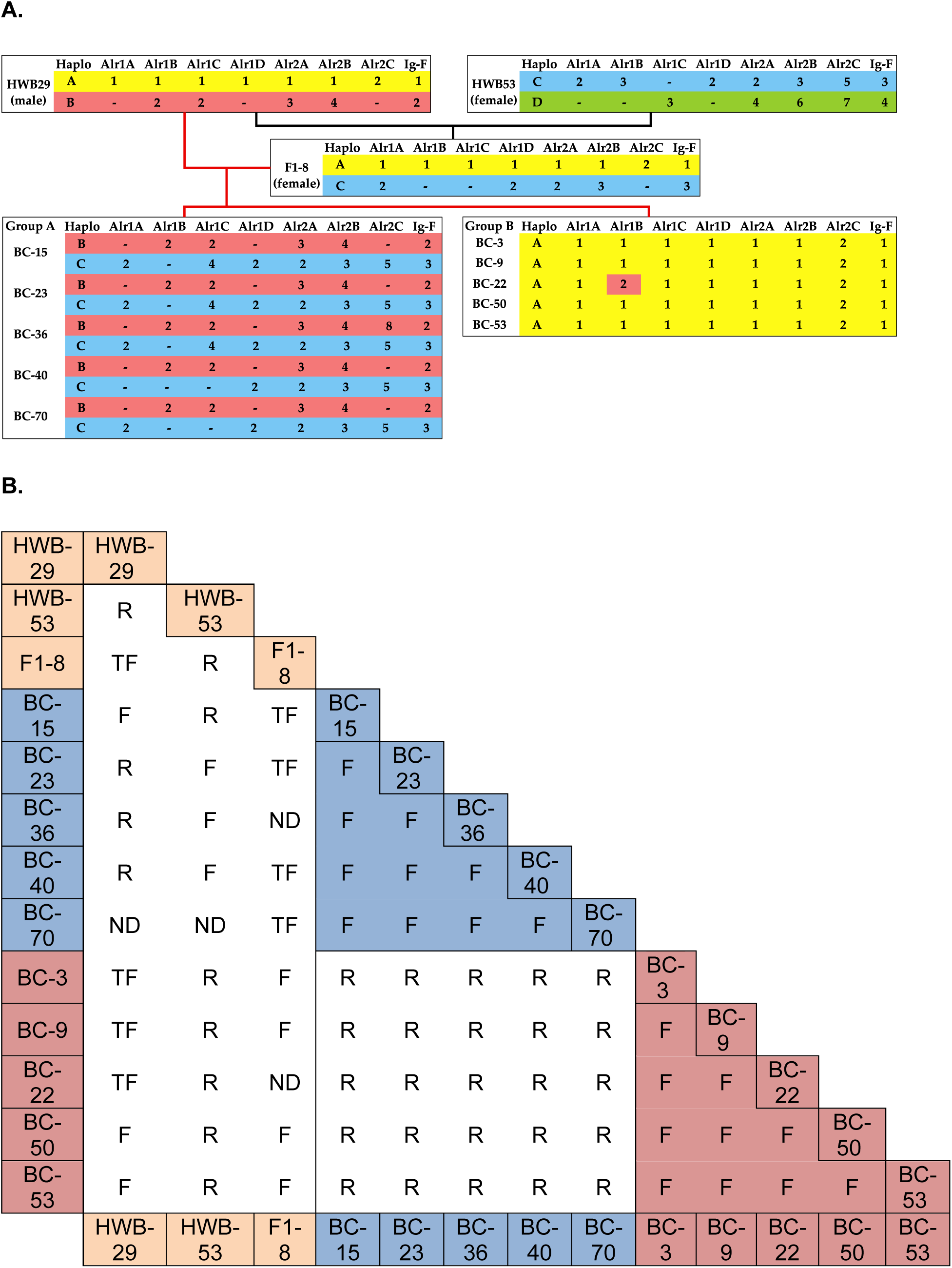
(A) Segregation of *alr1-type*, *alr2-type,* and *IgSF-like* F alleles in the backcross population. Four haplotypes for these genes were found in the wild-type individuals HWB29 (haplotypes A and B) and HWB53 (haplotypes C and D). Group A inherited two different haplotypes (haplotypes B and C) and group B inherited the two identical haplotypes (haplotypes A) from its parentals. Groups A and B have different haplotypes with the only exception of individual BC-22 that has the allele 2 of the *alr1B* gene, which is shared with the group A. Haplotypes A, B, C and D are highlighted in yellow, red, blue and green colors, respectively. BC: Backcross individual. Haplo: Haplotype. Dashes represent alleles that were not found. **(B) Allorecognition phenotypes of our backcross population.** Parentals, fusibility groups A and B individuals are highlighted in brown, blue and red, respectively. F: Fusion, R: Rejection and TF: Transitory fusion. ND: Non-determined

To determine the association of *alr* genes with allorecognition phenotypes, we established two filters based on the expected variation for i) the fusibility groups and ii) the progeny (F1 and backcross) and their parentals. For the first criterion, the individuals from fusibility groups A and B were found to not share alleles at the *alr1-type* (*alr1A*, *alr1B*, *alr1C* and *alr1D*) and *alr2-type* (*alr2A*, *alr2B*, *alr2C*) genes (Fig. 3A), which correlated with rejection observed between individuals from different fusibility groups (Fig. 3B). An exception to this result was allele 2 for the *alr1B* gene in individual BC-22 (group B), which shared this allele with the individuals of group A. Additionally, we found evidence that the individuals from the same fusibility group shared alleles for the *alr1-type* and *alr2-type* genes (Fig. 3A), which correlated with fusion observed between individuals from the same fusibility group (Fig. 3B). Although some alleles for certain individuals were not detected, two haplotypes were observed in fusibility group A (haplotypes B -red and C -blue), while a single and different haplotype was observed in fusibility group B (haplotype A -yellow) (Fig. 3A).

The second criterion was that the variability of the *alr1-type* and *alr2-type* genes correlated with the allorecognition phenotypes of the parental individuals HWB29, HWB53 and F1-8 (Fig. 3B). Wild-type individuals HWB29 and HWB53 had more or less two different haplotypes each. A single allele was detected for the *alr1A*, *alr1D* and *alr2C* genes in individual HWB29, while a single allele was detected for the *alr1A*, *alr1B*, *alr1C* and *alr1D* genes in individual HWB53. For a highly polymorphic gene such as the *alr1A* gene (Rosa et al. 2010), this could be explained by a primer-bias that amplified preferentially a group of alleles. Meanwhile, for a gene such as *alr1D* (Fig. 4D, nine variable residues between alleles 1 and 2), this could be explained by a low polymorphism of this gene in the population. Despite this, individuals HWB29 and HWB53 did not share any alleles for the *alr1-type* and *alr2-type* genes (Fig. 3A), which correlated with the rejection observed between these individuals (Fig. 3B). Meanwhile, F1-8 individual inherited a haplotype from each parental wild-type individual (haplotype A from individual HWB29 and haplotype C from individual HWB53) (Fig. 3A); therefore, fusion is expected between parents and offspring based on the current two-locus model. However, the allorecognition phenotype of F1-8 with individual HWB29 was a transitory fusion, and with individual HWB53 it was a rejection (Fig. 3B). This deviation from the predicted fusion between parents and their offspring indicates that sharing a haplotype at these *alr1-type* and *alr2-type* genes is not enough to establish fusion between these individuals. Explaining the allorecognition phenotypes between the parentals and their offspring, which are not always resolved in fusion, is one of the biggest challenges in *Hydractinia* allorecognition research. However, this conundrum could be partially addressed by considering that recombination could occur among *alr1-type* genes or *alr2-type* genes. For instance, *alr2-type* genes (*alr2A*, *alr2B* and *alr2C* genes) have a high similarity in their extracellular regions (Rosengarten et al. 2011; Gloria-Soria et al. 2012) (Fig. S3B), which could indicate that these genes recombine. Additionally, alternative splicing could also explain the discrepancy in the prediction of allorecognition phenotypes since we found evidence of alternative splicing for the *alr1A* gene, which is discussed later.

**Fig. 4.**
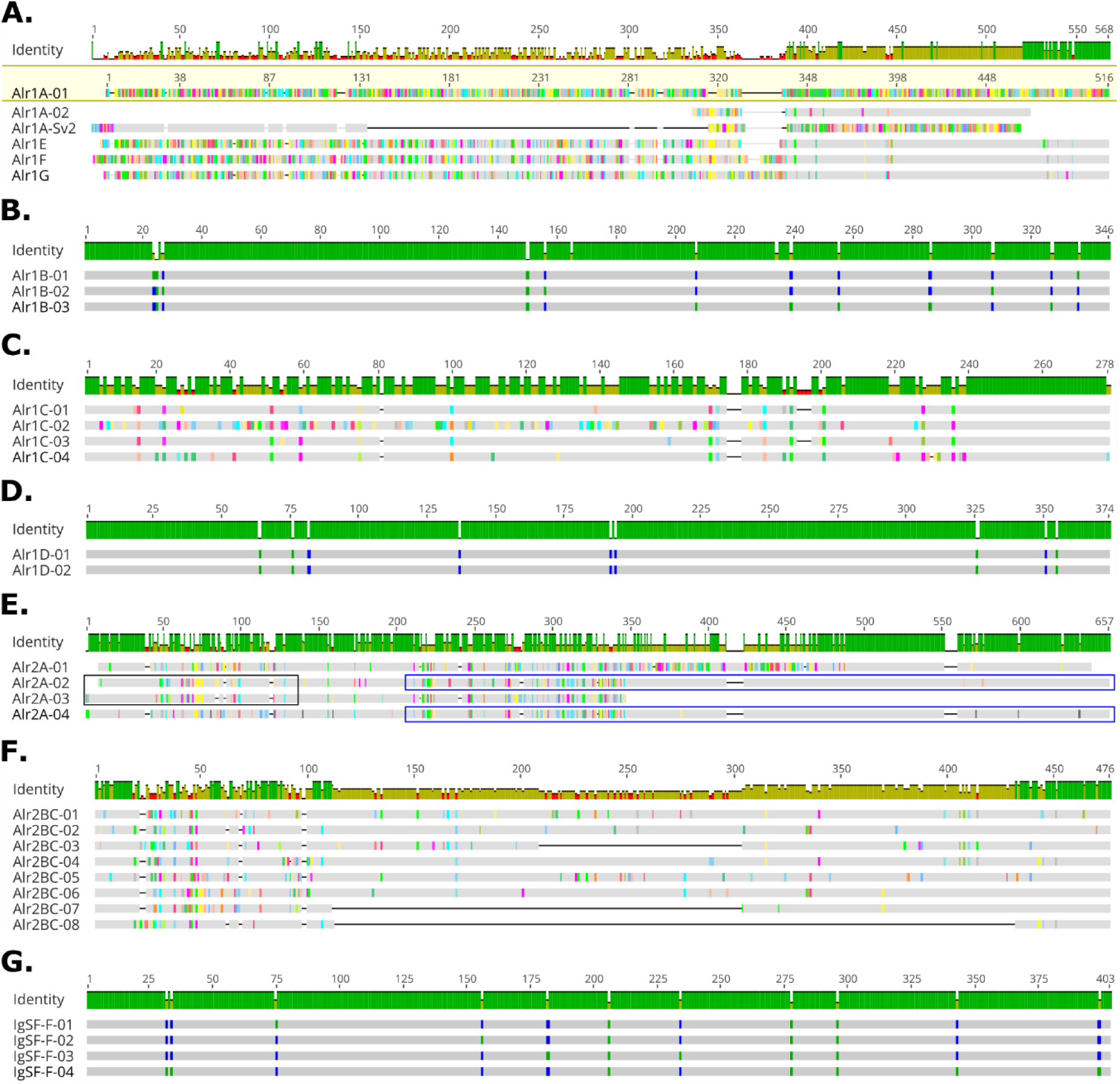
(A) Alignment of alleles of Alr1A, Alr1A-Sv2, Alr1E, Alr1F and Alr1G proteins. Notice that Alr1A and Alr1A-Sv2 share the N-terminal region of the protein, but they have different C-terminal regions. By contrast, Alr1A, Alr1E, Alr1F and Alr1G have different N-terminal regions, but they share the C-terminal region of the protein. In this alignment, the first sequence (Alr1A-01) is used as a reference to compare the other sequences. **(B) Alignment of Alr1B protein. (C) Alignment of Alr1C protein. (D) Alignment of Alr1D protein. (E) Alignment of Alr2A protein.** Evidence of inter-allelic recombination can be observed in the chimeric alleles 2 (recombination between alleles 3 and 4) and 3 (recombination between alleles 1 and 2). **(F) Alignment of Alr2BC proteins.** Notice that the N-terminal region (first Ig-domain) is the most variable region in these Alr2BC proteins. Alleles 3, 7 and 8 have deletions in the middle of the protein, however, they do not have premature stop codons. All eight Alr2BC alleles encode putative functional proteins. **(G) Alignment of IgSF-F protein.** In all alignments, the identical residues are shown in gray, and the variable residues are shown in other colors. With the exception of Alr1A alignment, where these colors indicate the conservation between the reference sequence (Alr1A-01) and the other sequences

A comparable situation was observed regarding the allorecognition phenotypes of fusibility groups A and B with their parentals HWB29, HWB53 and F1-8. For instance, each individual of fusibility group A shared at least one single haplotype with each parent, i.e., haplotype B was shared between the group A and individual HWB29 (Fig. 3A), and haplotype C was shared among group A and individuals F1-8 and HWB53 (Fig. 3A). Based on the two-locus model, these individuals should establish a fusion, which was observed in interactions between HWB29 and BC-15, HWB53 and BC-23, BC-36, and BC-40 (Fig. 3B). However, rejections and transitory fusions were also observed at a similar frequency. For example, HWB29 rejected individuals BC-23, BC-36, and BC-40; HWB53 rejected individual BC-15; and F1-8 showed transitory fusion with individuals BC-15, BC-23, BC-40, and BC-70. At the same time, fusibility group B shared haplotype A with individuals HWB29 and F1-8 (Fig. 3A), which predicts a fusion among these individuals. Although fusions were observed, the interactions between HWB29 and individuals BC-3, BC-9 and BC-22 showed transitory fusions (Fig. 3B). Finally, individual HWB53 and group B did not share any alleles (Fig. 3A), which predicts rejection phenotypes among these individuals. Indeed, HWB53 rejected all individuals of group B (Fig. 3B), as predicted by the two-locus model. Thus, the predictions of the two-locus model were more accurate for the allorecognition phenotypes among parentals and group B than with group A. This can be partially explained because group B was homozygous for the *alr1-type* and *alr2-type* genes, while group A was heterozygous for most genes. Thus, genetic homogenization of group B could have made the two-locus model more accurate for this group.

The pedigree shows missing alleles in individuals HWB29, HWB53, F1-8 and fusibility group A (Fig. 3A), despite sequencing a high number of clones in these individuals. For instance, for fusibility group A approximately 58 clones were sequenced for the *alr1A* gene, 55 clones for the *alr1B* gene, 153 clones for the *alr1C* gene, 59 clones for the *alr1D* gene, and 163 clones for the *alr2B* and *alr2C* genes. This suggests that primer-bias was likely present during the cloning process. For example, in fusibility group A, allele 2 of the *alr1C* gene was found in 149 clones, while allele 4 for the same gene was only found in four clones (Fig. 3A). This amplification bias was persistent across the *alr1-type* and *alr2-type* genes, and partially explains why some alleles were not detected. To prevent this primer-bias, a good practice in future approaches should include full-length transcriptome sequencing of *Hydractinia* individuals to rescue the complete set of alleles for these genes.

During the establishment of the fusibility groups A and B, a total of 215 fusibility assays were performed among the backcross population, where 89 resulted in fusion (41.4%), 95 in rejection (44.2%), and 31 in transitory fusion (14.4%) (Table S7). The last phenotype was actually a heterogeneous collection of phenotypes that varied in the duration of the initial fusion and the aggressiveness of the rejection response (Fig. S17). Moreover, we analyzed the proportion of fusions, rejections, and transitory fusions for each individual of our backcross population (Table S8). A slight increase in the frequency of rejection (mean: 48.5%) was observed compared with fusion (mean: 40.4%), while transitory fusion was found as the phenotype with lowest frequency (mean: 11.1%). However, a high variation was observed in some individuals where fusion had a higher or equal frequency than rejection; for instance, BC-59 individual showed 62.5% fusions and 12.5% rejections, and BC-3 individual displayed 40.6% fusions and 40.6% rejections (Table S8). These results limited being able to determine the inheritance pattern for these phenotypes.

### alr1A, alr1A splicing variants, and alr1E

Two alleles were found for the *alr1A* gene in our backcross population (Fig. 4A and Fig. S4). Allele 1 was detected in individuals HWB29, F1-8 and fusibility group B, while allele 2 was detected in HWB53, F1-8 and most individuals of fusibility group A (Fig. 3A). Although the same primers were used to clone these two alleles (Table S2), allele 2 showed a shorter extracellular region than allele 1 (Fig. 4A). Allele 2 (Alr1A-02 approximately 78 clones) does not have the two Ig-domains of the Alr1A protein, although it has a short extracellular region differing from allele 1 (Fig. 4A and Fig. S4). This small size of allele 2 could be explained by PCR artifacts in our cloning process; however, a biological explanation cannot be ruled out. Additionally, a splicing variant was observed for *alr1A* allele 2 that consists of an in-frame insertion (22 residues) before the TM domain (Fig. S4, Alr1A-02-Sv1, highlighted in a black rectangle), which resembles the splicing variant previously reported in exon 4 of this gene (Rosa et al. 2010). This splicing variant was named here *alr1A-Sv1*. This insertion was observed in HWB53, F1-8, and most group A individuals (BC-15, BC-23, BC-36, BC-70).

An additional *alr1A* splicing variant was identified in the transcriptome of individual HWB29 (i.e., HWB29_comp401220_c1_seq2, HWB29_comp401220_c1_seq8 and HWB29_comp418031_c5_seq20), which was mapped to the genomes of individuals BC-3 and BC-15, as well as the reported Alr1 interval (Rosa et al. 2010). This splicing variant uses exons 1 and 2 from the *alr1A* gene with four alternative exons located ∼3.3 Kb downstream from the last exon (i.e., exon 8) of the *alr1A* gene (Fig. 2A). This splicing variant was called *alr1A-Sv2*. In addition to the transcriptomic and genomic evidence, this alternative splicing variant was cloned in all individuals of fusibility group B. The reverse primer used to amplify this splicing variant was located in the alternative exon 5 (Fig. 2A), thus alternative exons 6, 7 and 8 were only confirmed by genomic and transcriptome evidence. Moreover, HWB29_comp401220_c1_seq8 and HWB29_comp418031_c5_seq20 transcripts retained fragments of the alternative intron 5 (i.e., 48bp and 1303bp, respectively) located downstream of alternative exon 5. These insertions caused premature stop codons after the TM domain, which contrasts with the HWB29_comp401220_c1_seq2 transcript that has a longer cytoplasmic region (data not shown). Whether the retention of alternative intron 5 represents an additional splicing variant or an artifact of the transcriptome assembly will be determined by further characterization of this gene.

The protein architecture of Alr1A-Sv2 is composed of a signal peptide and a single Ig-domain that is similar to the Alr1A protein. In contrast, this splicing variant is missing the second Ig-domain that is present in the Alr1A protein (Fig. 2A and Fig. S4). Additionally, the transmembrane and cytoplasmic regions of the Alr1A and Alr1A-Sv2 proteins are different. For instance, the ITAM-like motif previously reported for the Alr1A protein is missing in the Alr1A-Sv2 protein; however, the cytoplasmic region of Alr1A-Sv2 contains a lymphocyte-specific protein tyrosine kinase (Lck) motif in the alternative exon 8 (identified with Scansite software) (Obenauer et al. 2003) (Fig. 2A and Fig. S4). In T-cells, Lck is a kinase that phosphorylates the ITAM motifs of the TCR/CD3 complex once it is activated (Bommhardt et al. 2019). Remarkably, the motif in the Alr1A-Sv2 protein (EKRDQQ**VYAQV**DRSG) (Fig. S4A, blue box) resembles the ITIM-like motif (N**LYAQV**) present in the Alr2A protein (Nicotra et al. 2009), which could indicate that Alr1A-Sv2 is an inhibitory receptor. Thus, alternative splicing could be a mechanism used to switch a putative activating receptor, such as the Alr1A protein, to a putative inhibitory receptor, such as the Alr1A-Sv2 protein (Fig. S4B).

Furthermore, we found two transcripts (e.g., HWB29_comp418031_c5_seq18 and HWB29_comp401220_c1_seq12) in the transcriptome of individual HWB29 that suggested the presence of an additional *alr1-like* gene. These transcripts show a 3’ region identical to the *alr1A* gene, but a different 5’ region compared to this gene. Therefore, a new forward primer was designed to amplify this sequence (Fig. 2A and Fig. S5, *alr1E* gene). This gene was cloned in individual HWB29 (n=12 clones), most individuals from group A (BC-15, BC- 23, BC-36, and BC-70) (n=39 clones), and individual BC-3 (group B) (n=6 clones). Unlike the *alr1A* gene, this protein sequence was identical among our backcross individuals, including individual BC-3 compared with individuals of group A, indicating a non-polymorphic protein. The domain architecture of this new protein comprises a signal peptide, two Ig-domains and a transmembrane domain, such as the Alr1A protein (Rosa et al. 2010) (Fig. S5, *alr1E* gene). However, the first Ig-domain of this new protein could only be predicted using the Pfam software (Mistry et al. 2021) with a non-significant e-value (e-value 21), indicating that further characterization is necessary to validate the presence of this Ig-domain. Moreover, this protein showed low identity in the extracellular region compared with the Alr1A protein (i.e., approximately 34% with Alr1A-01). Conversely, the Alr1A protein and this new sequence showed a high identity in the transmembrane and cytoplasmic regions (approximately 90% with Alr1A-01), including complete conservation of the ITAM-like motif (Fig. 4A and Fig. S5, *alr1E* gene). Furthermore, these two sequences share the exons encoding the transmembrane and cytoplasmic regions, namely exons 5 to 8 for the *alr1A* gene and the putative exons 6 to 9 for this new sequence (Fig. 2A, *alr1E* gene). Since this new sequence shared the transmembrane and cytoplasmic regions with the Alr1A protein, but it has a different extracellular region, this gene was called *alr1E* (Fig. 2A and Fig. S5). Interestingly, the *alr1E* sequence was not predicted in the genomic assemblies of individuals BC-3 and BC-15, possibly due to assembly gaps. However, the preliminary *H. symbiolongicarpus* genome for the wild-type strain 291-10 has three scaffolds that partially cover this gene (Fig. 2A). The scaffold *flattened_line_795* covers the putative exons 6 to 9 of the *alr1E* gene, which is the region shared with the *alr1A* gene. Most importantly, scaffolds *flattened_line_20749* and *flattened_line_7125* span the putative exon 2, and exons 3 and 4, respectively (Fig. 2A). Additionally, two transcripts were identified in the transcriptome of individual HWB103 (comp57811_c0_seq1 and comp57811_c0_seq2), which partially span the extracellular region and completely cover the transmembrane and cytoplasmic region of the Alr1E protein (Fig. S5). Thus, *alr1E* gene was identified as a new *alr1-type* gene in *H. symbiolongicarpus* that shares the transmembrane and cytoplasmic regions with the *alr1A* gene. This gene requires further genomic characterization to completely determine its exon/intron structure and localization in the ARC complex.

### alr1B, alr1C and alr1D gene variation

The Alr1B protein domain structure is composed of a transmembrane domain based on the Interproscan software prediction (Hunter et al. 2012). A signal peptide was not identified in the sequences isolated from the BC-3 and BC-15 genomes or our backcross population. Additionally, two Ig-domains were predicted in the extracellular region of the Alr1B protein based on the Pfam software prediction (Mistry et al. 2021) (Fig. S6). However, these two domains were supported by non-significant e-values (e-values 0.11 and 27, respectively), indicating that further characterization of the extracellular region of this protein is necessary to confirm the existence of these Ig-domains. Moreover, three alleles for the *alr1B* gene were found in our backcross population, and twelve variable residues were identified in this protein (Fig. 4B and Fig. S6). Furthermore, a splicing variant was detected for allele 3 in individual HWB53, which skips the first region of exon 4 (37bp) (*alr1B-Sv1*) (Fig. 2A and Fig. S7). This deletion causes a frame-shift in the *alr1B* transcript that generates a protein with a premature stop codon before the transmembrane domain, encoding a possible secreted version of this protein (Fig. 2A and Fig. S6). Our gene-prediction strategy was unable to retrieve a full sequence for the cytoplasmic region of the Alr1B protein. However, manual inspection of the BC-3 and BC-15 genomes revealed that the full sequence for the cytoplasmic region of the Alr1B protein contains a putative ITAM motif (**Y**AD**I**NSKAVKEGI**Y**ED**L**) (Fig. S6), suggesting that this protein could be an activating receptor.

The Alr1C protein domain structure is composed of a signal peptide, two Ig-domains and a transmembrane domain (Fig. S8). However, the first Ig-domain could only be predicted using Pfam software (Mistry et al. 2021) with a non-significant e-value (e-value 400). This indicates that the first Ig-domain needs further characterization to validate its presence in this protein. Furthermore, four alleles were detected for the *alr1C* gene in our backcross population (Fig. 4C and Fig. S8). By comparing these alleles, we found approximately 105 variable positions out of 269-278 for the Alr1C protein, suggesting that this protein is more polymorphic than the Alr1B (12 variable positions out of 346; Fig. 4B) and Alr1D proteins (9 variable positions out of 374; Fig. 4D). Interestingly, the alleles of the Alr1C protein between the fusibility groups A and B showed a low identity, i.e., 69.4% between alleles 1 (group B) and 2 (group A), and 86.4% between alleles 1 (group B) and 4 (group A) (for instance, alleles 1 and 3 showed highest identity: 94.7%). This could partly explain the rejection between these two fusibility groups. In contrast, the last residues of the cytoplasmic region of the Alr1C protein were not variable. This region contains putative tyrosine-based motifs (i.e., ITAM or ITIM motifs) (Fig. S8). The differences in polymorphism between the extracellular and cytoplasmic regions of the Alr1C protein could be explained because the extracellular region must be variable to recognize putative polymorphic ligands, while the cytoplasmic region must be conserved to guarantee the correct functioning of the putative signaling pathways.

Additionally, five splicing variants were detected for the Alr1C protein (Fig. 2A and Fig. S8). The first variant (*alr1C-Sv1*, n= 7 clones) has lost the first region of exon 4 (37bp) (Fig. 2A and Fig. S9), causing a frameshift and a premature stop codon before the TM domain (Fig. 2A and Fig. S8), such as the *alr1B* splicing variant (*alr1B-Sv1*) described above (Fig. 2A). Prediction of the transmembrane domain using Phobius software (Käll et al. 2004) suggests that Alr1C-Sv1 protein has one TM domain (Alr1C-01-Sv1) or two TM domains (Alr1C-03- Sv1). However, these predictions were not supported by TMHMM software (Krogh et al. 2001), suggesting that the *alr1C-Sv1* sequence encodes an Alr1C secreted protein. Alr1C- Sv1 was detected for alleles 1 and 3 of the Alr1C protein. This splicing variant for allele 1 is inherited from individual HW29 to offspring F1-8 and BC-53 (Group B), while this splicing variant for allele 3 is only present in the individual HWB53. Furthermore, the second splicing variant detected for the Alr1C protein (*alr1C-Sv2*, n= 6 clones) retains the entire intron 3 (73bp) (Fig. 2A and Fig. S9) that causes a frameshift and a premature stop codon before the TM domain (Fig. 2A and Fig. S8). Similar to the previous splicing variant, the *alr1C-Sv2* sequence encodes an Alr1C secreted protein. *alr1C-Sv2* splicing variant was detected for the alleles 1, 3 and 4 of the Alr1C protein in the individuals F1-8, HWB53, and BC-23 (Group A), respectively. The third splicing variant of Alr1C protein (*alr1C-Sv3*, n= a single clone) retains a fragment of intron 3 (25bp), similar to *alr1C-Sv2*, and loses the first region of exon 4 (37bp), resembling *alr1C-Sv1* (Fig. 2A and Fig. S9). *alr1C-Sv3* splicing variant was detected for allele 3 in individual HWB53 and encodes an Alr1C secreted protein (Fig. 2A and Fig. S8). The fourth splicing variant for the Alr1C protein (*alr1C-Sv4*, n= a single clone) has a deletion in-frame of a fragment of exon 4 (177bp) (Fig. 2A and Fig. S9) and encodes an Alr1C transmembrane protein with a shorter extracellular region (Fig. 2A and Fig. S8). *alr1C-Sv4* splicing variant was detected for allele 4 in individual BC-36 (Group A). Finally, the fifth splicing variant of the Alr1C protein (*alr1C-Sv5*, n= 2 clones) has an insertion of a fragment of intron 4 (12bp) (Fig. 2A and Fig. S9) that encodes a small sequence (KLNC) before the TM domain (Fig. S8). This insertion in the *alr1C-Sv5* splicing variant was similar to allele 2 of the Alr1C protein (Alr1C-02) in that region (KLNN), which could be relevant in the variability of this protein. *alr1C-Sv5* splicing variant was identified for allele 3 in individual HWB53. Given that the *alr1C-Sv3*, *alr1C-Sv4* and *alr1C-Sv5* splice variants were detected in a small number of clones, thus a further characterization is needed to confirm their existence. However, the *alr1C-Sv1* and *alr1C-Sv2* splice variants were detected in a sufficient number of clones to allow affirming that secreted variants of the Alr1C protein can be generated through alternative splicing.

Alr1D protein domain structure is composed of a signal peptide and a transmembrane domain, according to Interproscan software (Hunter et al. 2012). In addition, two hypothetical Ig-domains could be predicted in the extracellular region of this protein with Pfam software (Mistry et al. 2021) (Fig. S10). However, these two Ig-domains were predicted with non-significant e-values (e-values 8100 and 0.089, respectively). Furthermore, two alleles for the *alr1D* gene were found in our backcross population (Fig. 4D and Fig. S10). We did not detect the other two alleles that would be expected in this population if this gene was highly polymorphic, despite the fact that many clones were sequenced (n=132 clones). Additionally, the two alleles identified for the *alr1D* gene showed nine different residues (Fig. 4D and Fig. S10). Together, these results could indicate that *alr1D* is not a highly polymorphic gene. Interestingly, Alr1C and Alr1D proteins have similar cytoplasmic regions that contain tyrosine-based motifs, which are **S**H**Y**AD**I**, **S**I**Y**AD**L** and KQ**Y**AE**I** for the Alr1C protein, and **S**H**Y**AD**L, S**I**Y**AD**L** and KK**Y**AE**I** for the Alr1D protein (Fig. S8 and Fig. S10). These motifs can be assembled into at least three possible combinations of ITAM (YxxI/Lx6- 12YxxI/L, where x is any residue) or ITIM (S/I/V/LxYxxI/V/L) motifs in the cytoplasmic regions: 1) Two ITIM motifs (**S**H**Y**AD**I** and **S**I**Y**AD**L** for the Alr1C protein; **S**H**Y**AD**L** and **S**I**Y**AD**L** for the Alr1D protein), 2) The first two motifs assemble a single ITAM motif (**Y**AD**I**NIKAVKPSI**Y**AD**L** for Alr1C; **Y**AD**L**NVKSVKPSI**Y**AD**L** for Alr1D); or 3) The first motif is an ITIM motif and the last two motifs assemble an ITAM motif (**S**H**Y**AD**I** and **Y**AD**L**TTTADHSKQ**Y**AE**I** for Alr1C; **S**H**Y**AD**L** and **Y**AD**L**TKATDPTKK**Y**AE**I** for Alr1D) (Fig.

S8 and Fig. S10). In contrast, the cytoplasmic region of the Alr1B protein did not have serine residues in the first two tyrosine-based motifs, indicating the lack of ITIM motifs in that region (first two motifs are <u>C</u>H**Y**AD**I** and <u>G</u>I**Y**ED**L**); instead, a single ITAM motif is present in the cytoplasmic region of this protein (**Y**AD**I**NSKAVKEGI**Y**ED**L**) (Fig. S6). Thus, additional experiments are needed to determine if the Alr1C and Alr1D proteins have two ITIM motifs, a single ITAM motif, or ITIM and ITAM motifs in the cytoplasmic region by establishing which motifs are functional and if these proteins are inhibitory or activating receptors.

### Additional alr1-like genes

The transcriptome of individual HWB103 demonstrated twelve transcripts (comp58736_c3_seq1 to comp58736_c3_seq11, comp58736_c3_seq16) that support the presence of a possible new *alr1-type* sequence. These transcripts were previously described for this transcriptome as *alr1A* gene sequences (Zárate-Potes et al. 2019), because they have almost identical transmembrane and cytoplasmic regions with the *alr1A* gene (95.5% protein identity in these regions) (Fig. S5, *alr1F* gene). Additionally, these sequences from individual HWB103 have high identities with some reported *alr1A* alleles, such as LH06-028a or LH07-041a, with 84% and 86% protein identity, respectively. However, the extracellular region of this *alr1-type* sequence shares 34.3% sequence identity with the Alr1A protein (the reference sequence for the Alr1A protein was the reported *alr1-f* allele from the original *Hydractinia* inbred line). Additionally, high variability in the extracellular region was observed when these sequences from individual HWB103 were compared with the *alr1A* sequences derived from our backcross population and the transcriptome of individual HWB29 (Fig. S5, *alr1A* and *alr1F* genes). Thus, the high variability in the extracellular region of this *alr1-type* gene from HWB103 transcriptome compared with Alr1A protein makes it difficult to conclude that these sequences from individual HWB103 are the *alr1A* gene. Furthermore, this pattern of conserved transmembrane and cytoplasmic regions and a variable extracellular region was also observed between *alr1A* and *alr1E* genes (Fig. S5). Therefore, we hypothesized that these transcripts from individual HWB103 correspond to a new *alr1-type* gene, which was called *alr1F* gene (Fig. S5). This protein has a signal peptide, two Ig-domains and a transmembrane domain. However, the first Ig-domain was predicted using Pfam software (Mistry et al. 2021) with a non-significant e-value (e-value 230) (Fig. S5, Alr1F protein). Accordingly, the description of this Ig-domain should be carefully considered until further experiments can be performed. In addition, the cytoplasmic region of this protein has an ITAM-like motif (**Y**P**I**DGAKASEPPT**Y**AP**V**), identical to the motif reported for the Alr1A protein (Rosa et al. 2010).

We analyzed the transcriptome of strain 291-10, sequenced by the *Hydractinia* Genome Project Team (https://research.nhgri.nih.gov/hydractinia/), to search for additional *alr1-type* sequences. Instead of finding the *alr1A* gene, we found a sequence with a variable extracellular region and conserved transmembrane and cytoplasmic regions compared with the Alr1A protein in this transcriptome (Fig. S5, *alr1G* gene). This pattern was the same as that found when Alr1E and Alr1F were compared to Alr1A. Thus, we consider that this sequence from the transcriptome of strain 291-10 could be a new *alr1-type* gene, named here as *alr1G* gene. Similar to the Alr1E and Alr1F proteins, the Alr1G protein comprises a signal peptide, two Ig-domains, and a transmembrane domain (Fig. S5). As described for the first Ig-domain of the Alr1E and Alr1F proteins, Pfam software (Mistry et al. 2021) predicts the first Ig-domain of Alr1G with a non-significant e-value (e-value 28). Therefore, future characterization of Alr1G protein will also aim to validate the presence of this Ig-domain. The cytoplasmic region of the Alr1G protein contains the same ITAM-like motif sequence as the one reported for the Alr1A protein (Fig. S5) (Rosa et al. 2010).

To summarize, four possible *alr1* genes (*alr1A*, *alr1E*, *alr1F*, and *alr1G*) were identified in different transcriptomes (from individuals HWB29, HWB103 and strain 291-10) that share almost identical transmembrane and cytoplasmic regions, including the ITAM-like motif, but have different extracellular regions with the presence of Ig-domains (Fig. S5). It is important to note that the *alr1F* and *alr1G* genes were not analyzed in our backcross population because these sequences were identified in subsequent analysis of the transcriptomes from individual HWB103 and strain 291-10, respectively. Thus, an additional cloning process is required for these genes to establish their polymorphism and correlation with the allorecognition phenotypes of *Hydractinia*.

Moreover, the transcriptome of individual HWB29 was analyzed for possible new *alr1-like* genes. We found a short sequence of the *alr1D* gene (comp413098_c0_seq3) that was almost identical to allele 1 of the *alr1D* gene amplified in our backcross population (Fig. S10, Alr1D-HWB29 sequence). Furthermore, two additional sequences were found that showed similarity to the *alr1D* gene (transcripts for <u>sequence 1</u>: comp413098_c0_seq5, comp413098_c0_seq10, comp415430_c0_seq22, comp415430_c0_seq25 and comp415430_c0_seq37; transcripts for <u>sequence 2</u>: comp415430_c0_seq2, comp415430_c0_seq16, comp415430_c0_seq18, comp415430_c0_seq24, comp415430_c0_seq27, comp415430_c0_seq30, comp415430_c0_seq35, comp415430_c0_seq41 and comp415430_c0_seq45) (Fig. S10, *alr1H* and *alr1I* genes). Each of these two proteins have a transmembrane domain and cytoplasmic region that contains two putative conserved ITIM motifs (**S**H**Y**AD**L** and **S**I**Y**AD**L**), or a single ITAM motif (Alr1H: **Y**AD**L**DVKSVKPSI**Y**AD**L**; Alr1I: **Y**AD**L**NVKSVKPSI**Y**AD**L**), or an ITIM motif and an ITAM-motif (Alr1H: **S**H**Y**AD**L** and **Y**AD**L**TKVSDSSKE**Y**AE**I**; Alr1I: **S**H**Y**AD**L** and **Y**AD**L**TKTSDSSKE**Y**AE**I**) (Fig. S10). This arrangement of tyrosine-based motifs resembles the cytoplasmic regions described above for the Alr1C and Alr1D proteins, which makes it difficult to establish if these are inhibitory or activating receptors (Fig. S8 and S10). In contrast, these two new sequences from the HWB29 transcriptome have different extracellular regions compared with the Alr1D protein (Fig. S10). As mentioned earlier, the Alr1D protein showed only nine variable positions in our backcross population (Fig. 4D). However, these two new sequences from the HWB29 transcriptome have a higher number of variable positions in the extracellular region compared with the Alr1D protein, suggesting the existence of two putatively new genes that were named *alr1H* and *alr1I* (Fig. S10). Importantly, *alr1H* and *alr1I* genes were detected in a re-analysis of HWB29 transcriptome and were not amplified in our backcross population; therefore, their variability and correlation with allorecognition phenotypes of *Hydractinia* will require further characterization.

Furthermore, another three putative new *alr1-like* genes were detected in the transcriptome of individual HWB29. Two of these genes differ in the N-terminal region including the signal peptide (approximately the first 136aa) (transcripts for <u>sequence 1</u>: comp426129_c3_seq4, comp426129_c3_seq9, comp426129_c3_seq10, comp426129_c3_seq12, comp426129_c3_seq14, comp426129_c3_seq19, comp426129_c3_seq22, comp426129_c3_seq24, comp426129_c3_seq25, comp426129_c3_seq26, comp426129_c3_seq28 and comp426129_c3_seq29; transcripts for <u>sequence 2</u>: comp420789_c0_seq6, comp420789_c0_seq10, comp426129_c3_seq1, comp426129_c3_seq3, comp426129_c3_seq6, comp426129_c3_seq8, comp426129_c3_seq11, comp426129_c3_seq17, comp426129_c3_seq20, comp426129_c3_seq21, comp426129_c3_seq23, comp426129_c3_seq27 and comp426129_c3_seq30). After this region, these two proteins share the same sequence for an Ig-domain, a transmembrane domain, and a cytoplasmic region that includes a putative ITIM motif (**V**I**Y**SD**V**) (Fig. S11). The similarity between these two sequences could indicate that they are encoded by the same gene. A BLASTp search showed that these sequences are similar to the reported allele LH06-082b of the *alr1A* gene (protein 1: e-value 2e-50; protein 2: e-value 8e-60). Since these sequences were similar to the *alr1A* gene, they were named *alr1J* and *alr1K* genes, respectively (Fig. S11). However, further experiments are needed to determine if these two similar sequences correspond to different genes or if they are products from the same gene.

Finally, a third sequence was identified in the transcriptome of individual HWB29 (transcripts: comp426236_c0_seq3, comp426236_c0_seq7, comp426236_c0_seq9, comp426236_c0_seq11, comp426236_c0_seq12, comp426236_c0_seq14, comp426236_c0_seq15, comp426236_c0_seq16, comp426236_c0_seq24, comp426236_c0_seq25, comp426236_c0_seq27 and comp426236_c0_seq28), which encodes a protein with a putative signal peptide, a transmembrane domain, and a cytoplasmic region. This last region could have two putative ITIM motifs (**S**Q**Y**TG**L** and **I**V**Y**AD**L**), or ITIM and ITAM motifs (**S**Q**Y**TG**L** and **Y**AD**L**SPPTVNN**Y**SE**I**), or a single ITAM motif (**Y**TG**L**QLEARKEIV**Y**AD**L**) (Fig. S12). This arrangement of tyrosine-based motifs resembles the cytoplasmic regions of the Alr1C, Alr1D, Alr1H and Alr1I proteins, where these proteins could be inhibitory or activating receptors (Fig. S8, Fig. S10, and Fig. S12). A BLASTp search showed that the most significant match of this sequence was with allele LH06-058a of the *alr1A* gene (e-value 5e-21), confirming that this sequence from HWB29 transcriptome is an *alr1-type* gene. Since this sequence was similar to the *alr1-type* genes and encodes a putative inhibitory or activating receptor, it was called the *alr1L* gene. It is important to note that the putative *alr1J*, *alr1K* and *alr1L* genes were not amplified in our backcross population, thus further experiments will be needed to determine the polymorphism and correlation with the phenotypes of allorecognition in *Hydractinia* of these genes.

### alr2A, alr2B and alr2C allelic variation

Four alleles were found for the *alr2A* gene in our backcross population (Fig. 4E and Fig. S13). However, the transmembrane and cytoplasmic regions were not sequenced for allele 3. Comparing these four alleles, the signal peptide and first Ig-domain were more similar between alleles 2 and 3 of the *alr2A* gene compared with alleles 1 and 4 (Fig. 4E and Fig. S13). Meanwhile, the second and third Ig-domains and transmembrane domain were more similar between alleles 2 and 4 of the *alr2A* gene compared with alleles 1 and 3 (Fig. 4E and Fig. S13). An interallelic recombination mechanism has been observed between reported type I and II alleles for the *alr2A* gene, which could explain the chimerism observed for the alleles of the *alr2A* gene in our backcross population (Rosengarten et al. 2011). Remarkably, the last part of the cytoplasmic region is conserved among the alleles of our backcross population, as well as the sequences derived from the genomes of the individuals BC-3 and BC-15, and the sequences isolated from the transcriptomes of individuals HWB29, HWB103 and strain 291-10 (Fig. S13). This conserved region of the Alr2A protein includes the reported ITIM-like motif (NL**Y**AQ**V**) that has been suggested to be functional in the signaling pathways of allorecognition (Nicotra et al. 2009). Additionally, allele 4 of the *alr2A* gene (n= 2 clones) was only found in the wild-type individual HWB53, which is reasonable based on how our crosses were designed, because this allele should not be inherited in our backcross population (Fig. 3A). This could explain why allele 4 was found in a low number of clones compared to alleles 1, 2, and 3. The last three alleles were present in several individuals of the backcross population, increasing their chance to be found in a higher number of clones. Another explanation can be attributed to the possible existence of a PCR-bias that favored the amplification of allele 2 (n= 24 clones) over allele 4 of the *alr2A* gene in individual HWB53 (Fig. 3A).

To analyze the *alr2B* and *alr2C* genes in our backcross population, a set of primers was initially designed to amplify each gene. However, these genes showed high similarity (Fig. S3B, Alr2BC alleles) and a single set of primers was enough to amplify both genes at the same time (Table S2). Although the genomes of individuals BC-3 and BC-15 made it possible to establish that they are two independent genes (Fig. 1), the high similarity among the *alr2B* and *alr2C* genes made it difficult to determine which alleles of our backcross population belonged to each gene (Fig. 4F). Thus, the eight alleles identified in our backcross population for the *alr2B* and *alr2C* genes (four alleles for each gene) were named as *alr2BC* genes (Fig. 4F and Fig. S14). Additionally, the alleles of the *alr2BC* genes showed differences in length; for instance, allele 3 lost exon 4 (third Ig-domain); allele 7 lost exons 3 (second Ig-domain) and 4; and allele 8 lost exons 3, 4, 5 (stem region) and 6 (transmembrane domain) (Fig. S15). Consequently, alleles 3 and 7 encoded Alr2BC transmembrane proteins with two and one Ig-domains, respectively, and allele 8 encoded Alr2BC secreted proteins with a single Ig-domain (Fig. S14). Furthermore, alleles 1, 2, 4, 5, and 6 showed the full structure of the Alr2BC proteins with three Ig-domains and a transmembrane domain (Fig. 4F and Fig. S14). Interestingly, all alleles and isoforms of the *Alr2BC* genes found in our backcross population retain the first Ig-domain (Fig. 4F and Fig. S14), which is the most variable domain compared to the second and third Ig-domains (Table S9, Table S10, and Table S11). Specifically, the first Ig-domain shows identities ranging from 59.8% (alleles 5 and 6) to 84.3% (alleles 6 and 7) (Table S9), while the second Ig-domain ranges from 86% (alleles 3 and 6) to 97.4% (for instance, alleles 2 and 4) (Table S10), and the third Ig-domain ranges from 83.1% (alleles 1 and 5) to 94.8% (for instance, alleles 2 and 4) (Table S11). A similar pattern has been observed for the Alr2A protein, in which the first Ig-domain is more variable (including several positions under positive selection) than the second and third Ig-domains (Rosengarten et al. 2011). Finally, the transmembrane and cytoplasmic regions of the Alr2BC proteins showed higher identities between alleles than the extracellular region, ranging from 85.7% (alleles 1 and 5) to 100% (alleles 6 and 7) (Table S12). This finding indicates that the cytoplasmic region is conserved in the Alr2BC proteins, which could be important in the signaling pathways of allorecognition.

Five splicing variants were detected for the *alr2BC* genes (Fig. 2B). Splicing variant 1 (*alr2BC-Sv1*) lost the last fragment of exon 5 (n= 19 clones) (88bp, stem region) (Fig. 2B and Fig. S15), causing a frameshift and a premature stop codon before the TM domain (Fig. 2B and Fig. S14). Thus, the *alr2BC-Sv1* splicing variant encodes a putative secreted version of the Alr2BC proteins with three Ig-domains, and it was observed for alleles 2, 5, 6, and 7 of the Alr2BC proteins. Furthermore, splicing variants 2, 3, and 4 (*alr2BC-Sv2*, *alr2BC-Sv3,* and *alr2BC-Sv4*) were observed for allele 3 of the Alr2BC proteins (Fig. 2B). The longest sequence amplified for allele 3 of the Alr2BC proteins completely loses exon 4 (third Ig-domain) (Fig. S15). The *alr2BC-Sv2* splicing variant (n= 18 clones) for this allele retains a fragment of exon 4 (77bp) (Fig. 2B and Fig. S15), and the *alr2BC-Sv3* splicing variant (n= 7 clones) retains a fragment of intron 4 (36bp) (Fig. 2B and Fig. S15). Meanwhile, the *alr2BC- Sv4* splicing variant (n= 28 clones) retains a fragment of 113bp derived from exon 4 (77bp) and intron 4 (36bp), which correspond to a combination of *alr2BC-Sv2* and *alr2BC-Sv3* splicing variants (Fig. 2B and Fig. S15). Therefore, *alr2BC-Sv2, alr2BC-Sv3* and *alr2BC-Sv4* splicing variants encode secreted isoforms of the Alr2BC proteins with two Ig-domains (Fig. 2B and Fig. S14). Moreover, splicing variant 5 (*alr2BC-Sv5*) of the Alr2BC proteins (n= 3 clones) loses exons 5 (stem region) and 6 (TM-Domain), corresponding to a fragment of 381bp (Fig. 2B and Fig. S15). Accordingly, the *alr2BC-Sv5* splicing variant, observed for allele 5, encodes secreted Alr2BC proteins with three Ig-domains (Fig. 2B and Fig. S14).

The cytoplasmic regions of the *alr2BC* alleles of our backcross population were partially sequenced (47 of 83 residues that constitute the cytoplasmic region), but this region showed that Alr2B and Alr2C proteins have similar cytoplasmic regions (Fig. 2B and Fig. S14). Moreover, the genomes of individuals BC-3 and BC-15 allowed retrieving the full-length cytoplasmic region of the Alr2B protein. Additionally, a complete cytoplasmic region for the Alr2BC proteins was found in the transcriptome of the individual HWB103 (Fig. S14). This cytoplasmic region has two ITAM motifs (**Y**SE**L**NIINVQHDL**Y**TG**L** and **Y**ED**L**KKKDITENI**Y**TD**L**), which are conserved in the eight alleles amplified in our backcross population (at least up to the region that was amplified) (Fig. S14). This finding suggests that the Alr2BC proteins could be activating receptors in contrast to the Alr2A protein that has been proposed as an inhibitory receptor (Nicotra et al. 2009).

### IgSF-like F gene variation

In addition to the *alr1-type* and *alr2-type* genes, we also determined that the *IgSF-like F* gene in the ARC region showed allelic variation that correlated with the allorecognition phenotypes of *Hydractinia* (Fig. 3A and Fig. 4G). This gene is located in ARC Interval 1 (Fig 1) and is encoded by six exons (Fig. 2B). The signal peptide is encoded by exon 1, the first and second Ig-domains are encoded by exons 2 and 3, the transmembrane domain is encoded by exon 5, and the cytoplasmic region is encoded by exon 6, which contains a putative ITAM motif (**Y**TD**L**SDHNKQT**Y**AE**L)** (Fig. 2B and Fig. S16). Four alleles for the *IgSF-like F* gene were found to segregate in the backcross population (Fig. 3A), which were defined by non-synonymous substitutions distributed predominantly in the two Ig-domains and stem region of this protein (Fig. 4G and Fig. S16). It is important to note that only eleven variable positions were detected in the IgSF-like F protein when the alleles from our backcross population were compared (Fig. 4G). Similar to the *alr1-type* and *alr2-type* genes, *IgSF-like F* gene was amplified in our backcross population to determine if its variation was associated with fusion phenotypes among individuals within the same fusibility group, and rejection among individuals from different fusibility groups. Indeed, the individuals from fusibility group A were heterozygous (alleles 2 and 3) for the *IgSF-like F* gene and shared the same alleles (Fig. 3A), while the individuals from fusibility group B were homozygous (allele 1) and shared the same allele (Fig. 3A), correlating with fusion phenotypes within the fusibility groups (Fig. 3B). In addition, the two alleles of fusibility group A were different from the single allele of fusibility group B (Fig. 3A and Fig. 4G), correlating with rejection among individuals from different fusibility groups (Fig. 3B). Furthermore, similar to the *alr1-type* and *alr2-type* genes, the *IgSF-like F* gene was heterozygous in the wild-type individuals HWB29 (alleles 1 and 2) and HWB53 (alleles 3 and 4), and alleles were not shared between these individuals (Fig. 3A). These differences correlated with rejection phenotype between these wild-type individuals (Fig. 3B and Fig. 4G). These findings allow proposing that *IgSF-like F* gene may participate in the allorecognition system of *Hydractinia*, in addition to *alr1-type* and *alr2-type* genes.

Nevertheless, as observed for the *alr1-type* and *alr2-type* genes, the *IgSF-like F* gene did not completely predict the allorecognition phenotypes among the parentals and their offspring. For instance, individuals of fusibility group A shared allele 3 of the *IgSF-like F* gene with the parental individuals F1-8 and HWB53 (Fig. 3A). According to this, these individuals should fuse; however, transitory fusions (F1-8 vs most individuals of group A) and rejections (HWB53 vs BC-15) were observed (Fig. 3B). Furthermore, HWB29 and individuals of fusibility group A shared allele 2 of the *IgSF-like F* gene (Fig. 3A) that predicts fusion or at least transitory fusion among these individuals, but rejections were observed between HWB29 and individuals BC-23, BC-36, and BC-40 (Fig. 3B). Meanwhile, fusibility group B shared allele 1 of the *IgSF-like F* gene with individuals HWB29 and F1-8 (Fig. 3A); however, transitory fusions were observed between HWB29 and individuals BC-3, BC-9, and BC-22 (Fig. 3B). Finally, individual HWB53 (alleles 3 and 4) and fusibility group B (allele 1) did not share any *IgSF-like F* alleles (Fig. 3A), correlating with the rejection phenotype among these individuals (Fig. 3B).

### Phylogenetic analysis of Alr1-type and Alr2-type proteins

To understand the evolution of Alr proteins, we analyzed the phylogenetic relationships of Alr1-type and Alr2-type proteins. The phylogenetic analysis across Alr1-type proteins was analyzed based on their extracellular regions, and a showed a separation into at least six different clades (Fig. 5). A first clade included the reported alleles *f* and *r* for the Alr1A protein from the inbred lines of *H. symbiolongicarpus* (HM070445.1_Alr1A_F_allele and HM070446.1_Alr1A_R_allele), while the second and third clades included sequences of the Alr1B and Alr1C proteins, respectively (Fig. 5). In addition, a fourth clade grouped the Alr1E, Alr1F proteins, an Alr1-type sequence from the HWB103 transcriptome, and a reported Alr1A allele (Fig. 5). A fifth clade included the Alr1D, Alr1H and Alr1I proteins, and two Alr1-type sequences from the HWB29 transcriptome; and a sixth clade grouped the Alr1J and Alr1K proteins (Fig. 5). Lastly, there were multiple sequences (Alr1G protein, Alr1L protein, an Alr1-type sequence from HWB29 transcriptome and two reported Alr1A alleles) that did not group into any specific clade (Fig. 5). These approximately six clades, together with the other sequences, established a basal polytomy that did not allow to resolve the phylogenetic relationships between these Alr1-type proteins based on their extracellular regions. This polytomy was still unresolved when using the complete protein sequences of the Alr1-type proteins (data not shown). Therefore, the phylogenetic relationships between Alr1-type proteins could not be clearly established, indicating that these genes may have different evolutionary origins, or have other molecular mechanisms acting on them, such as recombination between genes, making it difficult to establish how these genes have evolved. Rosa *et al*. (2010) found a close relationship between the reported *alr1* gene and certain *IgSF-like* genes that they referred to in detail (Rosa et al. 2010). It is possible that those sequences could correspond to the *alr1B, alr1C,* and *alr1D* genes characterized here.

**Fig. 5.**
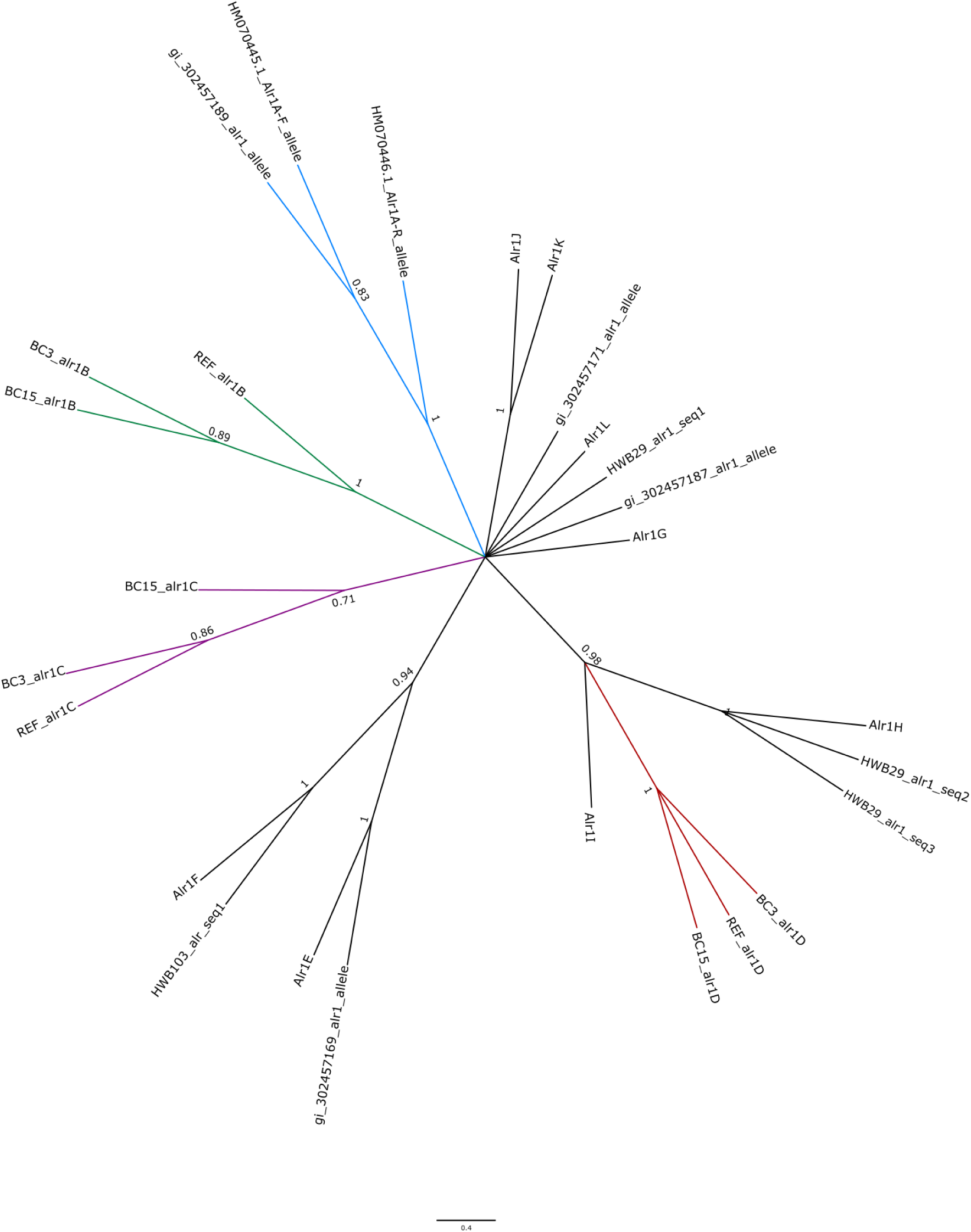
Genetic relationships among the Alr1-type proteins. Maximum likelihood-based phylogenetic tree was based on a multiple alignment of the partial extracellular region (second Ig-domain to the transmembrane region) of the Alr1-type proteins and selected GenBank reported Alr1A alleles. Bootstrap supports from 100 replicates are shown. The clades with the four Alr1-type proteins predicted by genomic evidence (Alr1A, Alr1B, Alr1C, and Alr1D) are highlighted in blue, green, purple, and red, respectively. The genomic *alr1A* sequences for BC-3 and BC-15 were incomplete in the extracellular region so the alignment was done with the reference Alr1A-F and Alr1A-R allele sequences. Alr1A reported alleles are identified according to their GenBank accession number

The phylogenetic analysis across proteins of the *alr2* gene complex showed that the Alr2B and Alr2C proteins form a sister clade that is different to the Alr2A protein clades (Fig. 6). In detail, Alr2B and Alr2C proteins from the BC-3 and BC-15 genomes formed a clade with transcripts from the HWB103 and HWB29 transcriptomes, which can only be classified as Alr2BC coding sequences (i.e., a clear distinction between these two genes is not possible based solely on the transcripts) (Fig. 6, blue clade). Additionally, we observed a clade composed of Alr2A sequences from the BC-3 and BC-15 genomes, HWB29 individual (Alr2A-allele 1), and reported Alr2A alleles (Fig 6, purple clade). Furthermore, three other clades were observed, which included two groups of reported Alr2A sequences and one group of Alr2A transcripts from HWB103 individual, reflecting an expected diversity of Alr2- type sequences (Fig 6, black clades). Thus, this phylogenetic analysis showed that Alr2B and Alr2C proteins are more closely related compared to Alr2A protein (Fig. 6). This close relationship between Alr2B and Alr2C genes, or Alr2A alleles could support a sequence donation scenario, similar to that proposed by (Rosengarten et al. 2011) and (Gloria-Soria et al. 2012) between certain *alr2A* alleles and the *alr2-type* pseudogene sequences.

**Fig. 6.**
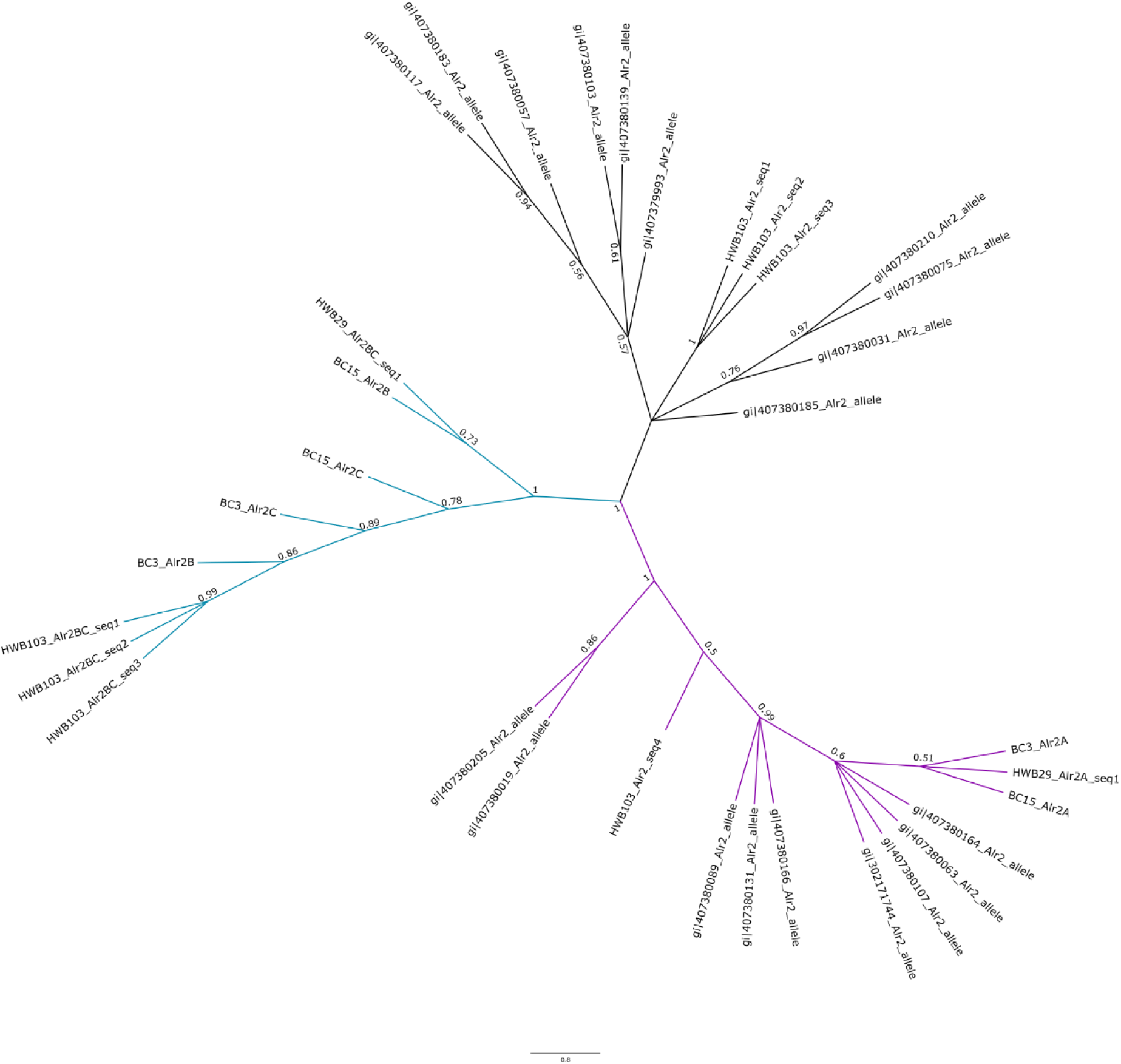
Genetic relationships among the Alr2-type proteins. Maximum Likelihood-based phylogenetic tree was based on a multiple alignment of the complete extracellular region (Ig-domains to the transmembrane region) of the Alr2-type proteins (Alr2A, Alr2B, Alr2C) and selected GenBank-reported Alr2A alleles. Bootstrap supports from 100 replicates are shown. Major clades are color shaded according to the location of Alr2A sequences (purple and black) and Alr2B and Alr2C sequences (blue). Reported Alr2A alleles are identified according to their GenBank accession number

### Sequence variation in additional genomic regions

The draft genome assemblies of BC-3 and BC-15 individuals provided complementary genomic data to preliminarily explore sequence variation outside of the ARC region that could potentially contribute to the allorecognition phenotypes of *Hydractinia*. The assemblies were generated from 188 million high-quality trimmed reads for BC-3 and 255 million high-quality reads for BC-15 (sequencing depths of ∼36X and ∼49X, respectively). This genomic data resulted from the paired-end and mate-pair libraries combined (Table S6). Total assembly lengths were approximately 436Mb and 420Mb for the BC-3 and BC-15 genomes, respectively, which account for ∼85% and ∼82% of the estimated genome size (514Mb) of

### H. symbiolongicarpus (Frank et al. 2020)

A total of 1897 and 5599 genes were predicted from 1767 and 4751 scaffolds equal or greater than 10 Kb in the genomes of BC-3 and BC-15, respectively. The almost three-fold difference in the number of predicted genes between the individuals is due to a more fragmented genome assembly of individual BC-3 compared to BC-15 (Table S6). Regardless, a total of 1406 homologous proteins were identified between BC-3 and BC-15 individuals, including 78 variable proteins not present in the ARC intervals. Among these proteins, 68 were classified based on their putative functional class (Fig. S2B and Table S13), while the remaining 10 could not be functionally annotated. Eight annotated proteins showed sequence variation potentially associated with allorecognition phenotypes among the genomic and transcriptomic sequences of individuals BC-3, BC-15, HWB29, and HWB103 (Table S14 and Fig. S18). Specifically, individuals HWB29 and BC-15 fused; HWB29 and BC-3 showed transitory fusion; and BC-3 and BC-15 rejected (Fig. 3B). Individual HWB103 rejected all individuals (HWB29, BC-3 and BC-15) from the backcross population. In this analysis, our eight candidate proteins showed high variability between individuals that rejected each other, and low variability between individuals that fused. For instance, BC-3 and BC-15 (rejection) had the highest genetic distances for the following proteins: CAP-containing protein (p-distance=0.59), importin alpha subunit protein (p-distance=0.52), HSP20-like protein (p-distance=0.49), and Fibrinogen-containing protein (p-distance=0.19) (Table S14). Meanwhile, Nidogen-Kazal-MAM-containing protein and Kazal- containing protein; Notch-like protein and Sushi-EGF-like-Ig-containing protein had the highest genetic distances between individuals BC-3 and HWB103 (rejection), and HWB29 and HWB103 (rejection), respectively (Table S14). In contrast, BC-15 and HWB29 (fusion) showed the lowest p-distances (0.00) for the eight proteins (Table S14). This subset of eight variable proteins contains domains such as Cysteine-rich secretory proteins, antigen 5, and pathogenesis-related 1 proteins (CAP), Importin beta binding (IBB), Armadillo repeats, Heat shock protein (HSP) 20-like, Fibrinogen-like (FBG), Nidogen, Kazal, and Meprin, A5 protein, and protein tyrosine phosphatase Mu (MAM), DSL (Delta/Serrate/Lag-2), Sushi, Epidermal growth factor (EGF)-like and Immunoglobulin (Ig) (Fig. S18). These candidate proteins identified here were not cloned in our backcross population and their possible polymorphism and actual correlation with the allorecognition phenotypes of *Hydractinia* will be established in future studies.

## DISCUSSION

The genomic, transcriptomic, and cloning evidence described here support a revised model for the control of allorecognition in *H. symbiolongicarpus.* This new model is composed of multiple *alr-type* and *IgSF*-like genes located mainly in the ARC region (Fig. 7). In agreement with this idea, Zárate-Potes *et al*., (2019) reported the gene expression of several highly variable *alr1*-type sequences that grouped into at least two clades, and hypothesized the presence of multiple transcriptionally active *alr1-type* genes. Additionally, their study showed that more than one *alr2-type* gene may be expressed due to the presence of five unique full-length *alr2-type* sequences (Zárate-Potes et al. 2019). Nicotra (2022) also suggested the presence of multiple *alr-type* genes in the ARC region that could participate in allorecognition phenotypes and other functions (Nicotra 2022). Accordingly, the *alr1-type*, *alr2-type*, and *IgSF-F* genes identified here correlated with allorecognition phenotypes within and between fusibility groups A and B of our backcross population. Moreover, these genes correlated with the allorecognition phenotypes between parentals and their offspring, although some deviations from these predictions were found. These results suggest that inheriting a haplotype is not always enough to establish a fusion phenotype. For instance, individual F1- 8 and its parentals HWB29 and HWB53 shared a haplotype, but they showed transitory fusion and rejection, respectively, instead of the predicted fusion. Thus, the two-locus model must be reviewed to explain the allorecognition phenotypes in *Hydractinia*. Our study describes new putative genes that could control the allorecognition phenotypes in this species and could be incorporated into a new model. Furthermore, the model should reconsider the assumption that sharing at least a haplotype between individuals is sufficient to establish a fusion. Because, even if the number of allorecognition genes that make up the haplotype increases to three or more genes, this model will always predict fusion, which has been shown to be incorrect for the allorecognition phenotypes between parent and offspring. Therefore, a revised model must incorporate not only new genes, but other molecular mechanisms of variation, such as alternative splicing and/or haplotype variation to provide a more holistic explanation of allorecognition in *Hydractinia*.

**Fig. 7.**
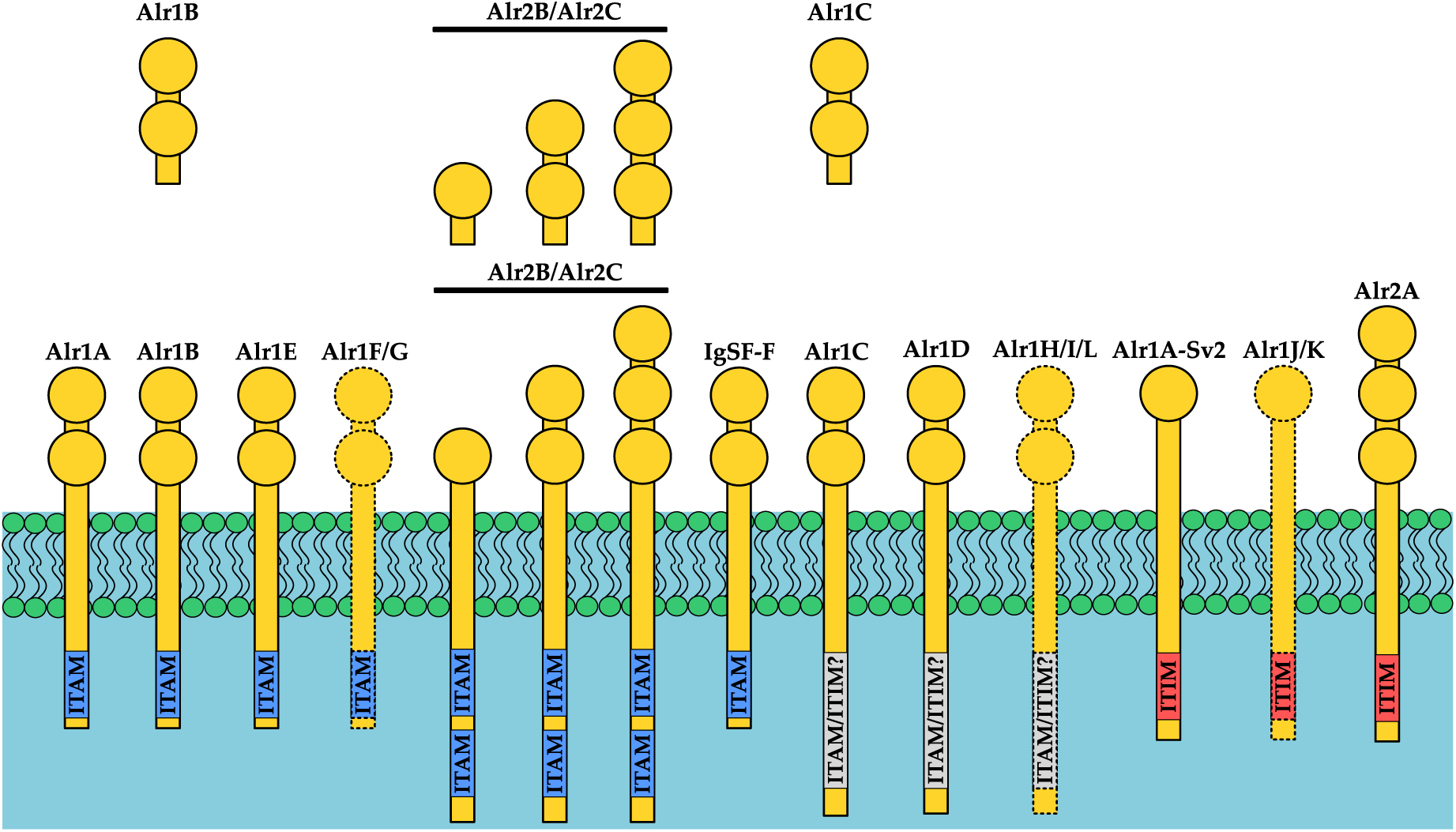
Multi-IgSF-like gene model of control of allorecognition in *H. symbiolongicarpus*. Alr1-type, Alr2-type and IgSF-like F proteins showed allorecognition- associated variability in *H. symbiolongicarpus*. Alr1F, Alr1G, Alr1H, Alr1I, Alr1L, Alr1J and Alr1K proteins only had transcriptomic evidence (dotted lines). ITAM/ITAM-like motifs and ITIM/ITIM-like motifs are represented in blue and red boxes, respectively. Alr1C, Alr1D, Alr1H, Alr1I and Alr1L proteins can have ITAM or ITIM motifs in the cytoplasmic region (see text). Alr1J and Alr1K proteins could have additional unidentified Ig-domains in the extracellular region. Secreted versions of Alr1B, Alr1C, Alr2B and Alr2C proteins were identified in our backcross population. Alr2B and Alr2C receptors and secreted isoforms can have one, two or three Ig-domains (the first highly polymorphic Ig-domain is always present)

According to our findings, the allorecognition system of *Hydractinia* could comprise putative inhibitory receptors, such as Alr1A-Sv2, Alr1J, Alr1K and Alr2A proteins; putative activating receptors, such as Alr1A, Alr1B, Alr1E, Alr1F, Alr1G, Alr2B, Alr2C and IgSF-F proteins; and putative activating/inhibitory receptors, such as Alr1C, Alr1D, Alr1H, Alr1I and Alr1L proteins (Fig. 7). This last category of receptors needs further characterization to establish whether ITAM or/and ITIM motifs are functional in the cytoplasmic regions of these proteins. Interestingly, Alr1A protein that has an ITAM-like motif in the cytoplasmic region, was found to use alternative splicing to generate a variant called Alr1A-Sv2 with a putative ITIM-like motif in the cytoplasmic region. A slightly similar example has been observed in human T-cells, where the inhibitory receptor SLAMF6 uses alternative splicing to skip a fragment of its extracellular Ig-domain and generate the costimulatory receptor SLAMF6^Δ17–65^ (Hajaj et al. 2021). A well-known example is the CD45 receptor that is a protein tyrosine phosphatase with multiple isoforms, which change their expression through T-cell differentiation and maturation. For instance, human naive T-cells express the splice form CD45RA, while the memory T-cells express the splice form CD45RO; or double negative and late single positive T-cells express the splicing variants CD45R**A**, CD45R**B**, and CD45R**C,** while double positive and early single positive T-cells express the splice variant CD45R**O** (Rheinländer et al. 2018). A similar regulation could be occurring in the Alr1A receptor of *Hydractinia* using alternative splicing. For example, the Alr1A receptor could be constitutively expressed, while the expression of the Alr1A-Sv2 splice form could be induced once compatible colonies come into contact to inhibit rejection through its putative ITIM-like motif. Conversely, if the colonies are incompatible, the Alr1A receptor would be the only molecule expressed, which would induce rejection through its putative ITAM-like motif. Moreover, a previous study has shown that the Alr1 and Alr2 proteins display an allele-specific homophilic *trans* interaction (Karadge et al. 2015), indicating that *Hydractinia* colonies use these receptors to recognize self-molecules. However, the cytoplasmic region of the Alr1A protein and other proteins described here have ITAM-like motifs that likely participate in an activating signaling pathway of allorecognition, which could induce rejection. But if these activating receptors recognized self-proteins, it would be contradictory to their function of activating rejection. Thus, it is possible that the inhibitory splice variant Alr1A-Sv2 is overexpressed compared with the activating Alr1A protein during fusion, while underexpression occurs during rejection. Nevertheless, this hypothesis involves several assumptions about the molecular mechanism of allorecognition in *Hydractinia* that still need to be investigated, so it should be carefully analyzed.

Furthermore, our results showed that the Alr2B and Alr2C receptors have two ITAM motifs in the cytoplasmic region, contrasting with the presence of an ITIM-like motif in the cytoplasmic region of the Alr2A protein (Fig. 7) (Nicotra et al. 2009). Moreover, the extracellular regions of the *alr2A*, *alr2B*, and *alr2C* genes are difficult to distinguish, suggesting that these genes could have evolved from an ancestral *alr2* gene. Following duplication of the *alr2-type* genes, these could have evolved independently to generate unique cytoplasmic regions. In addition, the extracellular regions of the *alr2-type* genes could be recombining since these regions are encoded by the same number of exons (five exons) and these genes have the same orientation in the genome. These conditions are known to enable recombination of the Killer Ig-like receptors (KIRs) in the human NK-cells (Pende et al. 2019). In our analysis, interallelic recombination for the *alr2A* gene was detected, as has been previously reported (Rosengarten et al. 2011; Gloria-Soria et al. 2012). However, further exploration will be required to detect recombination among the extracellular regions of different *alr2-type* genes.

The presence of ITAM and ITIM motifs in the allorecognition proteins of *Hydractinia* suggests that the molecular mechanism of allorecognition in this cnidarian can consist of activating and inhibitory signaling pathways, similar to the missing-self mechanism of NK-cells in humans (Myers and Miller 2021). Interestingly, the same molecular mechanism has also been suggested to explain the allorecognition response of the ascidian *B. schlosseri* (De Tomaso 2009). Determining if the genome of *Hydractinia* contains related proteins involved in the signaling pathways of NK-cells, such as ZAP70, SHP1/2, or SHIP proteins (Vivier et al. 2004) will be an essential question to understand the signaling pathways of allorecognition in this species. In this direction, a Syk protein-tyrosine kinase has been reported for *Hydra vulgaris* that showed a close phylogenetic relationship with the ZAP70 protein of humans (Steele et al. 1999).

Moreover, the secreted variants of the Alr1B, Alr1C, Alr2B, and Alr2C proteins detected in our backcross population are likely generated through an alternative splicing mechanism (Fig. 7). We identified Alr2B and Alr2C receptors with one, two or three Ig-domains, and secreted isoforms with the same variation in the number of Ig-domains. Remarkably, both Alr2B and Alr2C receptors and secreted isoforms keep the first Ig-domain, which is the most polymorphic domain compared to the second and third Ig-domains. A possible scenario in which these secreted proteins could function would be the contact zone between colonies. However, the interaction of these proteins in the extracellular space raises questions about the signaling process of these secreted proteins or the significance of these interactions. The *Fester* allorecognition receptor in *Botryllus* has shown extensive use of alternative splicing to generate individual-specific splice variants and putative secreted proteins, whose functions still need to be characterized (Nyholm et al. 2006). This evidence could indicate that alternative splicing is a common molecular mechanism that allorecognition proteins of *Hydractinia* and *Botryllus* use to generate variability. Thus, the function of these secreted proteins in the allorecognition system of *Hydractinia* will be an interesting question to explore in the future.

Our data showed that some *alr1-type* genes were expressed only in a specific group of individuals; for instance, the *alr1F* gene in individual HWB103; the *alr1G* gene in strain 291- 10; the *alr1B* gene in the individuals of our backcross population; the *alr1H* and *alr1I* genes in individuals HWB29 and HWB103; and the *alr1J*, *alr1K,* and *alr1L* genes in individual HWB29. These findings indicate that at least the *alr1-type* genes are expressed at an individual-specific level, and *alr-type* genes could display haplotypic variation. To confirm this hypothesis, it will be necessary to sequence the genomes of different *Hydractinia* individuals. Moreover, the fact that Alr1A, Alr1E, Alr1F, and Alr1G proteins share transmembrane and cytoplasmic regions suggests that these genes are products of recent duplication events and/or there is an evolutionary pressure on these cytoplasmic regions to be conserved as essential components of the signaling process of these receptors.

Regarding the *alr2-type* genes, the reported *Hydractinia* inbred strains have two *alr2-type* pseudogenes (Nicotra et al. 2009; Rosengarten et al. 2011; Gloria-Soria et al. 2012), while these sequences represented two putative functional *alr2-type* genes in our backcross population. This pseudogenization of the *alr2-type* genes can be explained by the homogenization process of the genetic background in the inbred lines, possibly generating premature stop codons in these genes. We did not detect haplotype variation of the *alr2- type* genes in our backcross population. However, haplotypes have been reported with two and three *alr2-type* pseudogenes, indicating that natural populations could have a variation in the number of functional *alr2-type* genes (Rosengarten et al. 2011). In addition, the *alr- type* genes characterized here could represent only a subset of the total *alr-type* genes in the genome of *Hydractinia*, as has been suggested by (Nicotra 2022). The idea of haplotypic variation in the ARC region is supported by a previous study that demonstrated a size variation of 1.6 to 30.5 cM in this genomic region. This variation could be explained by a different number of *alr-type* genes in this region across individuals (Powell et al. 2011). A well-known example of haplotype variation has been described for the KIRs receptors that are expressed on the surface of human NK-cells. Across human populations the number of KIRs receptors that are present is different and this variability can be classified into two haplotypes, namely A and B. Haplotypes A contain a single activating KIR receptor (surrounded by inhibitory receptors), while haplotypes B can have a higher number of activating KIR receptors (Pende et al. 2019). Moreover, the NK-cells of mice express analogous receptors, called Ly49s, that are based on C-type lectin-like domains, which display haplotype variation in the four inbred mouse strains analyzed to date (Carrillo-Bustamante et al. 2016). This suggests that haplotype variation can be a molecular mechanism extended throughout metazoa that generates variability in immune receptors, such as KIRs, Ly49s, and Alrs.

In our analysis of variable genes outside the ARC region, we identified eight candidate proteins with domains other than IgSF-like domains (Fig. S18). Several of these domains are involved in immunity, recognition, protein binding, and/or cell-cell adhesion. For instance, in the ascidian *Halocynthia roretzi* the PR (plant pathogenesis-related 1) domain (belonging to the CAP domain family) is present in the HrUrabin protein that indirectly participates in the self-incompatibility system of this species (Urayama et al. 2008). Additionally, an Importin alpha subunit protein (with importin beta binding -IBB- domain and Armadillo repeats) was detected in our analysis, which has a similar domain composition to importin-α proteins. These proteins participate in gametogenesis and recognition during fertilization in *Drosophila melanogaster, Caenorhabditis elegans,* and *Mus musculus* (Tewari et al. 2010). Furthermore, a HSP20-like protein was found among our candidates, which is of interest because HSP proteins have been observed in other allorecognition systems. For example, in *B. schlosseri* the HSP40-like protein is a variable molecule that correlates with fusion and rejection phenotypes (Nydam et al. 2013). Similarly, a HSP70 protein has been found that participates in the self-incompatibility system of *Ciona intestinalis* (Marino et al. 1998). On the other hand, we found a fibrinogen-containing protein among our candidates, these proteins have shown to be highly polymorphic in several invertebrates, displaying multiple gene copies and recombination events that can promote functional diversity (Hanington and Zhang 2011). For instance, fibrinogen domains are present in the v-Themis proteins of *Ciona intestinalis* type A, which have been implicated in the self-incompatibility system of this ascidian (Ben-Shlomo 2008; Harada and Sawada 2008). Moreover, one of our candidate proteins contains EGF, Sushi, and Ig domains, indicating a putative function in cell adhesion, ligand-receptor binding, or recognition. These three domains are found separately in allorecognition proteins of *B. schlosseri*, namely FuHc-Tm (Ig-like domains), FuHc-Sec (EGF-like domain), Fester, and Uncle fester (Sushi domain) proteins (Taketa and De Tomaso 2015). Lastly, the remaining domains found in our analysis (i.e., Kazal, Nidogen, DSL and MAM) have not been reported in proteins involved in allorecognition. Related to that, it is difficult to establish whether genes other than *alr-type* genes could control the allorecognition in *Hydractinia*. Nonetheless, our genomic comparison of two histo- incompatible siblings (individuals BC-3 and BC-15) clearly shows variable genes other than *alr-type* genes that could be associated with allorecognition in *Hydractinia*, and the characterization of these genes will require more experiments.

Several challenges were encountered in this study to clone the *alr-type* genes due to the high similarity of their IgSF-like domains and/or transmembrane and cytoplasmic regions (for instance, *alr2B* and *alr2C* genes). Additionally, the likely existence of primer-bias increased the difficulties in cloning these genes, because some alleles were amplified more frequently than others. However, future work should use long-read sequencing technologies that can enable simultaneously genotyping and haplotyping gene complexes, such as the *Hydractinia* ARC. The new *alr-type* and *IgSF-like* genes identified in this study represent a step towards explaining the allorecognition phenotypes of *Hydractinia*. Therefore, it is necessary to combine different molecular mechanisms, such polymorphism, alternative splicing, haplotype variation, recombination between different genes, interallelic recombination, among others, in a new model that allows explaining allorecognition phenotypes in this cnidarian.

## ACKNOWLEDGEMENTS

We wish to thank Dr. Andrés Pinzón for sharing his server to handle the genomic and transcriptomic data of *Hydractinia*. We would also like to thank Dr. Uri Frank for sharing the high-molecular weight DNA isolation protocol used in his laboratory. This study was funded by grant numbers 014-2013 from COLCIENCIAS, and Hermes 37463 from the Universidad Nacional de Colombia to Luis F. Cadavid.

