## Supplementary material for "Multiple *alr* genes exhibit allorecognition-associated variation in the colonial cnidarian *Hydractinia*"

**Table S1** BAC clones used to assemble the Allorecognition Complex (ARC) reference (Intervals alr1, alr2, Interval 1, Interval 2, and Interval 3)

| BAC clone ID | GenBank accession | Length (bp) | ARC interval assembled (Total length in bp) |
|---|---|---|---|
| YU0092J03 | AC234888.1 | 108925 | alr1 (1225305) |
| YU0089N10 | AC234887.1 | 171723 | |
| YU0084I16 | AC234883.1 | 151185 | |
| YU0062M14 | AC234878.1 | 144062 | |
| YU0060F17 | AC234877.1 | 134186 | |
| YU0031K01 | AC234871.1 | 131056 | |
| YU00081B18 | AC234869.1 | 98434 | |
| YU00066K10 | AC234868.1 | 120907 | |
| YU00048G22 | AC234867.1 | 93389 | |
| YU00025C05 | AC234865.1 | 137755 | |
| YU0001F08 | AC234864.1 | 113159 | |
| YU00018B13 | AC234863.1 | 100329 | |
| YU0087P24 | AC234885.1 | 174175 | alr2 (522055) |
| YU0077A14 | AC234881.1 | 93377 | |
| YU0064J23 | AC234880.1 | 153466 | |
| YU0063P12 | AC234879.1 | 99738 | |
| YU0025P04 | AC234870.1 | 109343 | |
| YU0089C02 | AC234886.1 | 96041 | Interval 1 (586429) |
| YU0086A23 | AC234884.1 | 212725 | |
| YU0060B22 | AC234876.1 | 111793 | |
| YU0002M08 | AC234866.1 | 89120 | |
| YU00012A13 | AC234862.1 | 101389 | |
| YU00006C08 | AC234861.1 | 119205 | |
| YU0033F02 | AC234872.1 | 111111 | Interval 2 (207512) |
| YU0028G22 | AC234793.1 | 104311 | |
| YU007J18 | AC234882.1 | 93641 | Interval 3 (214692) |
| YU0051G19 | AC234874.1 | 148550 | |

**Table S2** Primers used in RT-PCR for amplifying candidate allorecognition-associated genes

| Protein | | Primer sequence (5'-3') | Expected size (bp) |
|---|---|---|---|
| *IgSF-like F* | Forward | TCATTTCATTTCTCGCTGTTGC | 1252 |
| | Reverse | GCCGTACCCTCCATGTTTTC | |
| *dnaJ-like* | Forward | ATGTGGGCGGAATCCTCTTG | 1463 |
| | Reverse | AATCCACCGCCACCAAAATC | |
| *CAP-containing* | Forward | CCAGAACGGCCAATGAGAAG | 653 |
| | Reverse | GGAACGTAAAGACCTCCGTT | |
| *FBG-containing* | Forward | AAGAGATCAGGAGGTGCTACT | 945 |
| | Reverse | CACTTGTGGTTGGTCGTAGC | |
| *hsp20-like* | Forward | TGTACCATTTCCAGTCAGTGC | 824 |
| | Reverse | TTCTAAGGTTCTTGGCTGCG | |
| *alr1A* | Forward[1] | ATAGAACCGTAATCAATGGGAAT | 1573 |
| | Reverse[2] | TTGTATTGCGATCGATATTGC | |
| *alr1A-Sv2* | Reverse | CTATGCCAACCACAATCAGTG | 512 |
| *alr1B* | Forward | TGGCACTAAGTTGGAATTGG | 1041 |
| | Reverse | GCCTCGTAATCGTTAGTACGTT | |
| *alr1C* | Forward | GTGGATATCCTAACCCTGAAGTT | 807-880 |
| | Reverse | TCCCTGATATTGGAGTATGTGTC | |
| *alr1D* | Forward | CGATACAGGTCAAACAATTAACAG | 1124 |
| | Reverse | AGCGTAAATACTCGGTTACACAG | |
| *alr1E* | Forward | TGTACCGTTGGTAGCGGTATT | 1510 |
| *alr1A and alr1E* | Reverse | CCTTCATGATCATCCTCATCTG | |
| *alr2A* | Forward[3] | ACCAAATGATGAAAATATGCTGAAC | 1865-1901 |
| | Reverse[4] | CTTCAGGAGGGTCACAAACTT | |
| *alr2B and alr2C* | Forward | TCCTGTKTATCTSTCACCTCTCC | 1411(most alleles) |
| | Reverse | TGTCTTTGTTAGACCTGTGTATAAATC | |

[1]Primer modified from LH07-026/LH07-046 5'end primer. [2]Primer used previously to amplify f and r alleles of the *alr1* gene (Rosa et al. 2010). [3]Primer modified from P3402 published primer. [4]Primer modified from P3981 published primer (Gloria-Soria et al. 2012).

**Table S3** Genomic mapping and assembly statistics of the Allorecognition Complex (Intervals alr1, alr2, Interval 1, Interval 2, and Interval 3) for individuals BC-3 and BC-15

| Individual | Statistic | ARC Interval | | | | |
|---|---|---|---|---|---|---|
| | | alr1 | alr2 | Int 1 | Int 2 | Int 3 |
| BC-3 | Mapped reads [a] (%) | 0.77 | 0.57 | 0.64 | 0.17 | 0.25 |
| | Mapped scaffolds (%) | 7.79 | 6.47 | 6.54 | 3.09 | 4.47 |
| | Sequence[b] length (bp) | 1226896 | 522047 | 590140 | 216169 | 214576 |
| | Gaps, Ns (%) | 8.96 | 5.46 | 3.04 | 4.02 | 6.35 |
| | Undefined bases in sequence[c] (%) | 1.43 | 1.10 | 1.73 | 0.51 | 3.37 |
| BC-15 | Mapped reads [a] (%) | 1.24 | 0.89 | 1.06 | 0.31 | 0.49 |
| | Mapped scaffolds (%) | 3.51 | 2.51 | 2.63 | 1.14 | 1.8 |
| | Consensus sequence [b] length (bp) | 1225296 | 529880 | 589488 | 212109 | 214692 |
| | Gaps, Ns (%) | 7.28 | 1.40 | 0.60 | 1.69 | 0.25 |
| | Undefined bases in sequence[c] (%) | 1.28 | 0.52 | 1.32 | 1.13 | 1.24 |

[a] Paired-end and mate-pair libraries combined
[b] Merged sequence from independently mapped reads and scaffolds
[c] R, Y, S, W, K, M; according to the IUPAC nucleotide code

**Table S6** Statistics of the draft genome assemblies for BC-3 and BC-15 individuals of *H. symbiolongicarpus*

| Individual | | Scaffolds ≥ 500 pb | Scaffolds ≥ 10 Kb |
|---|---|---|---|
| BC-3 | Number of scaffolds | 93859 | 1767 |
| | Total span (Mb) | 436.47 | 26.16 |
| | Scaffold N50 (bp) | 3598 | |
| | Largest scaffold (bp) | 86698 | 86698 |
| | Average scaffold length (bp) | 2197 | 14802 |
| | GC content (%) | 34.75 | |
| BC-15 | Number of scaffolds | 40525 | 4751 |
| | Total span (Mb) | 420.31 | 93.42 |
| | Scaffold N50 (bp) | 11203 | |
| | Largest scaffold (bp) | 130686 | 130686 |
| | Average scaffold length (bp) | 4259 | 19662 |
| | GC content (%) | 33.21 | |

**Table S7** Allorecognition phenotypes of the backcross population of *H. symbiolongicarpus*

| | Backcross population | | | | | | | | | | | | | | | | | | | | | | | | | | | | | | | | |
|---|---|---|---|---|---|---|---|---|---|---|---|---|---|---|---|---|---|---|---|---|---|---|---|---|---|---|---|---|---|---|---|---|---|
| | **15** | **23** | **36** | **40** | **70** | **3** | **9** | **22** | **50** | **53** | **43** | **5** | **7** | **11** | **24** | **25** | **26** | **29** | **31** | **33** | **38** | **41** | **44** | **51** | **56** | **59** | **61** | **62** | **64** | **66** | **67** | **68** | **71** |
| **15** | | F | F | F | F | R | R | R | R | R | F | TF | F | F | R | R | TF | F | TF | F | TF | TF | F | R | F | F | F | F | F | F | F | F | TF |
| **23** | | | F | F | F | R | R | R | R | R | F | | TF | TF | | | | R | | | | F | F | R | | R | F | | | | | R | |
| **36** | | | | F | F | R | R | R | R | R | F | | TF | F | | | | R | | | | TF | F | R | | F | F | | | | | R | |
| **40** | | | | | F | R | R | R | R | R | F | | R | R | | | | R | | | | TF | F | R | | R | F | | | | | | |
| **70** | | | | | | R | R | R | R | R | F | | | R | | | | | | | | | | R | | TF | F | | | | | R | |
| **3** | | | | | | | F | F | F | F | F | TF | R | R | TF | F | F | R | TF | TF | TF | TF | R | F | R | F | R | R | R | F | F | F | F |
| **9** | | | | | | | | F | F | F | F | TF | R | R | F | TF | F | R | F | TF | F | F | R | F | | F | R | | | | | F | |
| **22** | | | | | | | | | F | F | F | | R | R | | | | R | | | | R | R | F | | F | R | | | | | F | |
| **50** | | | | | | | | | | F | TF | | R | R | | | | R | | | | R | R | F | | F | R | | | | | F | |
| **53** | | | | | | | | | | | TF | | R | R | | | | R | | | | R | R | F | | TF | R | | | | | F | |
| **43** | | | | | | | | | | | | | F | R | | | | | | | | | | F | | F | F | | | | | | |
| **5** | | | | | | | | | | | | | R | R | | | | | | | | | | | | | | | | | | | |
| **7** | | | | | | | | | | | | | | R | F | R | R | TF | F | F | TF | R | TF | R | | TF | | | | | | | |
| **11** | | | | | | | | | | | | | | | R | R | TF | R | R | R | TF | R | F | R | | F | R | | | | | F | |
| **24** | | | | | | | | | | | | | | | | | | | | | | | | | | | | | | | | | |
| **25** | | | | | | | | | | | | | | | | | | | | | | | | | | | | | | | | | |
| **26** | | | | | | | | | | | | | | | | | | | | | | | | | | | | | | | | | |
| **29** | | | | | | | | | | | | | | | | | | | | | | R | F | R | | | | | | | | | |
| **31** | | | | | | | | | | | | | | | | | | | | | | | | | | | | | | | | | |
| **33** | | | | | | | | | | | | | | | | | | | | | | | | | | | | | | | | | |
| **38** | | | | | | | | | | | | | | | | | | | | | | | | | | | | | | | | | |
| **41** | | | | | | | | | | | | | | | | | | | | | | | R | R | | | | | | | | | |
| **44** | | | | | | | | | | | | | | | | | | | | | | | | R | | | | | | | | | |
| **51** | | | | | | | | | | | | | | | | | | | | | | | | | | F | | | | | | | |
| **56** | | | | | | | | | | | | | | | | | | | | | | | | | | | | | | | | | |
| **59** | | | | | | | | | | | | | | | | | | | | | | | | | | | TF | | | | | F | |
| **61** | | | | | | | | | | | | | | | | | | | | | | | | | | | | | | | | F | |
| **62** | | | | | | | | | | | | | | | | | | | | | | | | | | | | | | | | | |
| **64** | | | | | | | | | | | | | | | | | | | | | | | | | | | | | | | | | |
| **66** | | | | | | | | | | | | | | | | | | | | | | | | | | | | | | | | | |
| **67** | | | | | | | | | | | | | | | | | | | | | | | | | | | | | | | | | |
| **68** | | | | | | | | | | | | | | | | | | | | | | | | | | | | | | | | | |
| **71** | | | | | | | | | | | | | | | | | | | | | | | | | | | | | | | | | |

F: Fusion, R: Rejection, TF: Transitory fusion. Individuals from the fusibility groups A and B are highlighted in blue and red colors, respectively.

**Table S8 Allorecognition phenotypes per backcross individual**

| Colony | n | Fusions (%) | Rejections (%) | TF (%) |
|---|---|---|---|---|
| **BC-3** | 32 | 13 (40.6) | 13 (40.6) | 6 (18.8) |
| **BC-7** | 23 | 5 (21.7) | 12 (52.2) | 6 (26.1) |
| **BC-9** | 26 | 13 (50.0) | 10 (38.5) | 3 (11.5) |
| **BC-11** | 26 | 5 (19.2) | 18 (69.2) | 3 (11.5) |
| **BC-15** | 32 | 18 (56.3) | 8 (25.0) | 6 (18.8) |
| **BC-22** | 19 | 8 (42.1) | 11 (57.9) | 0 (0.0) |
| **BC-23** | 19 | 8 (42.1) | 9 (47.4) | 2 (10.5) |
| **BC-29** | 14 | 2 (14.3) | 11 (78.6) | 1 (7.1) |
| **BC-36** | 19 | 9 (47.4) | 8 (42.1) | 2 (10.5) |
| **BC-40** | 18 | 7 (38.9) | 10 (55.6) | 1 (5.6) |
| **BC-41** | 14 | 2 (14.3) | 8 (57.1) | 4 (28.6) |
| **BC-44** | 14 | 6 (42.9) | 7 (50.0) | 1 (7.1) |
| **BC-50** | 19 | 7 (36.8) | 11 (57.9) | 1 (5.3) |
| **BC-53** | 19 | 6 (31.6) | 11 (57.9) | 2 (10.5) |
| **BC-51** | 17 | 7 (41.2) | 10 (58.8) | 0 (0.0) |
| **BC-59** | 16 | 10 (62.5) | 2 (12.5) | 4 (25.0) |
| **BC-61** | 14 | 7 (50.0) | 6 (42.9) | 1 (7.1) |
| **BC-68** | 12 | 9 (75.0) | 3 (25.0) | 0 (0.0) |
| **BC-70** | 15 | 6 (40.0) | 8 (53.3) | 1 (6.7) |
| **Mean** | | 7.8 (40.4) | 9.3 (48.5) | 2.3 (11.1) |

**Table S9** Identity matrix for the first Ig-domain of the Alr2BC proteins

**Alr2BC Alleles – First Ig-domain**

| | | | | | | | |
|---|---|---|---|---|---|---|---|
| **Alr2BC-01** | **Alr2BC-01** | | | | | | |
| **Alr2BC-02** | 73.7 | **Alr2BC-02** | | | | | |
| **Alr2BC-03** | 76.4 | 74.5 | **Alr2BC-03** | | | | |
| **Alr2BC-04** | 72.7 | 76.7 | 71.5 | **Alr2BC-04** | | | |
| **Alr2BC-05** | 68.6 | 70.7 | 66.6 | 66.6 | **Alr2BC-05** | | |
| **Alr2BC-06** | 63.7 | 64.7 | 60.9 | 64.7 | 59.8 | **Alr2BC-06** | |
| **Alr2BC-07** | 66.0 | 66.0 | 64.0 | 61.0 | 62.0 | 84.3 | **Alr2BC-07** |
| **Alr2BC-08** | 72.5 | 84.1 | 81.3 | 71.5 | 67.6 | 60.0 | 62.1 |

**Table S10** Identity matrix for the second Ig-domain of the Alr2BC proteins

**Alr2BC Alleles – Second Ig-domain**

| | | | | | |
|---|---|---|---|---|---|
| **Alr2BC-01** | **Alr2BC-01** | | | | |
| **Alr2BC-02** | 88.6 | **Alr2BC-02** | | | |
| **Alr2BC-03** | 88.6 | 88.6 | **Alr2BC-03** | | |
| **Alr2BC-04** | 91.1 | 97.4 | 88.6 | **Alr2BC-04** | |
| **Alr2BC-05** | 91.1 | 97.4 | 89.8 | 97.4 | **Alr2BC-05** |
| **Alr2BC-06** | 88.6 | 94.9 | 86.0 | 97.4 | 94.9 |

**Table S11** Identity matrix for the third Ig-domain of the Alr2BC proteins

**Alr2BC Alleles – Third Ig-domain**

| | | | | |
|---|---|---|---|---|
| **Alr2BC-01** | **Alr2BC-01** | | | |
| **Alr2BC-02** | 90.9 | **Alr2BC-02** | | |
| **Alr2BC-04** | 88.3 | 94.8 | **Alr2BC-04** | |
| **Alr2BC-05** | 83.1 | 84.4 | 84.4 | **Alr2BC-05** |
| **Alr2BC-06** | 87.0 | 93.5 | 94.8 | 85.7 |

**Table S12** Identity matrix for the transmembrane (TM) and cytoplasmic regions of the Alr2BC proteins

**Alr2BC Alleles – TM and cytoplasmic regions**

| | | | | | | |
|---|---|---|---|---|---|---|
| **Alr2BC-01** | **Alr2BC-01** | | | | | |
| **Alr2BC-02** | 88.5 | **Alr2BC-02** | | | | |
| **Alr2BC-03** | 91.4 | 92.8 | **Alr2BC-03** | | | |
| **Alr2BC-04** | 95.7 | 90.0 | 92.8 | **Alr2BC-04** | | |
| **Alr2BC-05** | 85.7 | 94.2 | 92.8 | 87.1 | **Alr2BC-05** | |
| **Alr2BC-06** | 90.0 | 98.5 | 94.2 | 91.4 | 95.7 | **Alr2BC-06** |
| **Alr2BC-07** | 90.0 | 98.5 | 94.2 | 91.4 | 95.7 | 100 |

**Table S14** Genetic distances for the eight candidate allorecognition-proteins outside of the ARC

| Pairwise comparison (variation measured as p-distance[a]) | | | | | | |
|---|---|---|---|---|---|---|
| | BC-3 vs. | | | BC-15 vs. | | HWB29 vs. |
| | BC15 | HWB29 | HWB103 | HWB29 | HWB103 | HWB103 |
| **Phenotype[b]/Protein** | R | TF | R | F | R | R |
| CAP-containing protein | **0.59[c,d]** | 0.02 | 0.58[d] | 0.00 | 0.04 | 0.02 |
| Importin alpha subunit protein | **0.52[d]** | 0.00 | 0.01 | 0.00 | 0.01 | 0.01 |
| HSP20-like protein | **0.49[d]** | 0.00 | 0.48[d] | 0.00 | 0.01 | 0.01 |
| Fibrinogen-containing protein | **0.19[e]** | 0.01 | 0.02 | 0.00 | 0.01 | 0.02 |
| Nidogen-Kazal-MAM-containing protein | 0.02 | 0.01 | **0.05** | 0.00 | 0.02 | 0.01 |
| Kazal-containing protein | 0.02 | 0.02 | **0.22** | 0.00 | 0.18 | 0.02 |
| Notch-like protein | 0.01 | 0.01 | 0.01 | 0.00 | 0.01 | **0.03** |
| Sushi-EGF-like-Ig-containing protein | 0.01 | 0.01 | 0.01 | 0.00 | **0.02** | **0.02** |

**a.** The genetic distance was measured as Nei's p-distance based on pairwise protein alignments.

**b.** Allorecognition phenotypes: Fusion (F), rejection (R) and transitory fusion (TF).

**c.** Highest p-distances for each protein are highlighted in bold.

**d.** This p-distance is attributed to variation across different genes that encode the same type of protein.

**e.** This p-distance is attributed to variation due to recombination of protein variants.

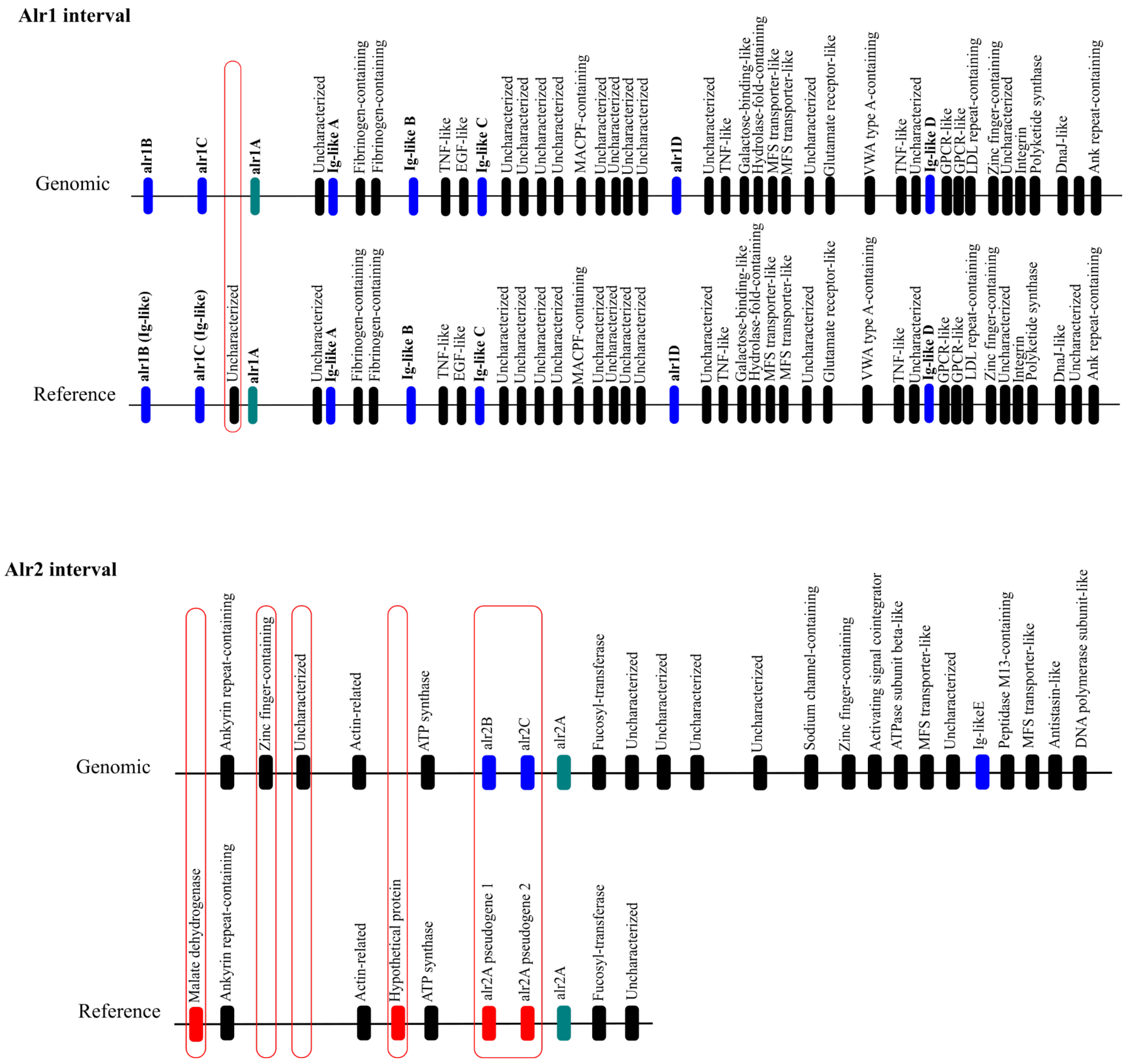


**Fig. S1 Comparison of gene content of the alr1 and alr2 intervals between the genomic scaffolds sequenced and the reference regions.** Reference sequences were assembled from BAC clones available in GenBank. Genomic scaffolds from individual BC-15 are shown on top of each comparison. Top: Genomic (BC-15) and reference Alr1 interval comparison. Bottom: Genomic (BC-15) and reference Alr2 interval comparison. Genes are depicted by boxes and the annotation is provided above each box. *alr1A* and *alr2A* genes are shown in green boxes; *Ig-like* genes are shown in blue boxes, including new characterized *alr-type* genes; absent genes in the genomic sequences (BC-15 or reference) are outlined in red

(a)

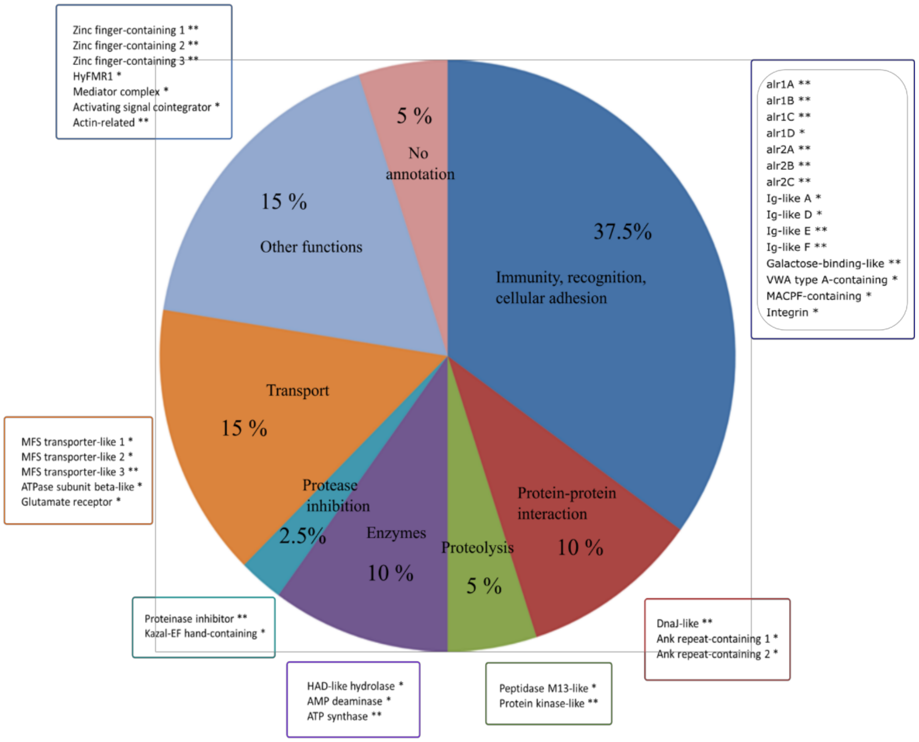

Zinc finger-containing 1 **
Zinc finger-containing 2 **
Zinc finger-containing 3 **
HyFMR1 *
Mediator complex *
Activating signal cointegrator *
Actin-related **
5 %
No annotation
15 %
Other functions
37.5%
Immunity, recognition, cellular adhesion
alr1A **
alr1B **
alr1C **
alr1D *
alr2A **
alr2B **
alr2C **
Ig-like A *
Ig-like D *
Ig-like E **
Ig-like F **
Galactose-binding-like **
VWA type A-containing *
MACPF-containing *
Integrin *
Transport
15 %
MFS transporter-like 1 *
MFS transporter-like 2 *
MFS transporter-like 3 **
ATPase subunit beta-like *
Glutamate receptor *
Protease inhibition
2.5%
Enzymes
10 %
Proteolysis
5 %
Protein-protein interaction
10 %
Proteinase inhibitor **
Kazal-EF hand-containing *
HAD-like hydrolase *
AMP deaminase *
ATP synthase **
Peptidase M13-like *
Protein kinase-like **
DnaJ-like **
Ank repeat-containing 1 *
Ank repeat-containing 2 *


(b)

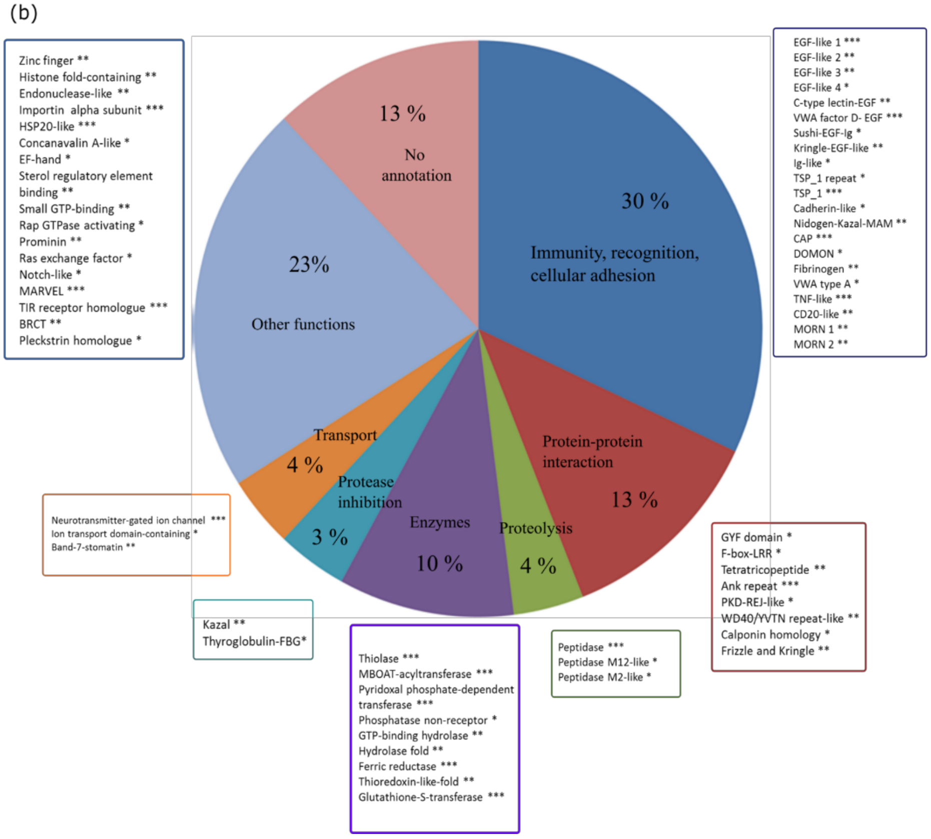

Zinc finger **
Histone fold-containing **
Endonuclease-like **
Importin alpha subunit ***
HSP20-like ***
Concanavalin A-like *
EF-hand *
Sterol regulatory element binding **
Small GTP-binding **
Rap GTPase activating *
Prominin **
Ras exchange factor *
Notch-like *
MARVEL ***
TIR receptor homologue ***
BRCT **
Pleckstrin homologue *
13 %
No annotation
30 %
Immunity, recognition, cellular adhesion
23%
Other functions
EGF-like 1 ***
EGF-like 2 **
EGF-like 3 **
EGF-like 4 *
C-type lectin-EGF **
VWA factor D- EGF ***
Sushi-EGF-Ig *
Kringle-EGF-like **
Ig-like *
TSP_1 repeat *
TSP_1 ***
Cadherin-like *
Nidogen-Kazal-MAM **
CAP ***
DOMON *
Fibrinogen **
VWA type A *
TNF-like ***
CD20-like **
MORN 1 **
MORN 2 **
Transport
4 %
Protease inhibition
3 %
Enzymes
10 %
Proteolysis
4 %
Protein-protein interaction
13 %
Neurotransmitter-gated ion channel ***
Ion transport domain-containing *
Band-7-stomatin **
Kazal **
Thyroglobulin-FBG*
Thiolase ***
MBOAT-acyltransferase ***
Pyridoxal phosphate-dependent transferase ***
Phosphatase non-receptor *
GTP-binding hydrolase **
Hydrolase fold **
Ferric reductase ***
Thioredoxin-like-fold **
Glutathione-S-transferase ***
Peptidase ***
Peptidase M12-like *
Peptidase M2-like *
GYF domain *
F-box-LRR *
Tetratricopeptide **
Ank repeat ***
PKD-REJ-like *
WD40/YVTN repeat-like **
Calponin homology *
Frizzle and Kringle **

**Fig. S2 (A) Variable proteins between BC-3 and BC-15 individuals within the ARC. (B) Variable proteins between BC-3 and BC-15 individuals outside of the ARC.** Proteins are classified according to their putative function based on predicted domain architecture. Asterisks represent sequence variation levels (Nei's p-distance) between encoded proteins in individuals BC-3 and BC-15: *** (p-distance $\geq 0.20$), ** ($0.02 \leq$ p-distance $< 0.20$), * (p-distance $< 0.02$)

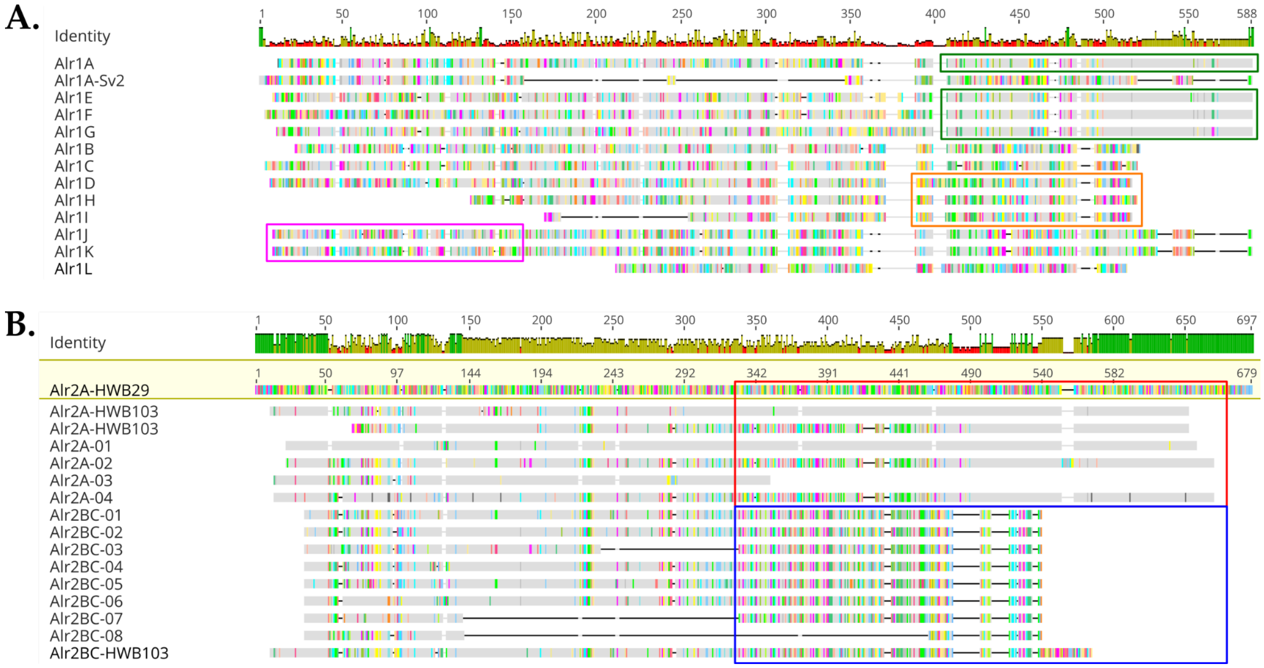


**Fig. S3 Summary of Alr1-type and Alr2-type proteins. (A) Alignment of Alr1-type proteins.** Alr1A, Alr1E, Alr1F and Alr1G proteins have similar transmembrane and cytoplasmic regions (green rectangles). Alr1D, Alr1H and Alr1I proteins have similar transmembrane and cytoplasmic regions (orange rectangle). Alr1J and Alr1K proteins differ only in the N-terminal region (pink rectangle). **(B) Alignment of Alr2-type proteins.** Alr2A and Alr2BC proteins have different stem, transmembrane and cytoplasmic regions that are highlighted in the red (Alr2A protein) and blue rectangles (Alr2BC proteins). However, these three proteins have a similar N-terminal region that contains Ig-domains

A.

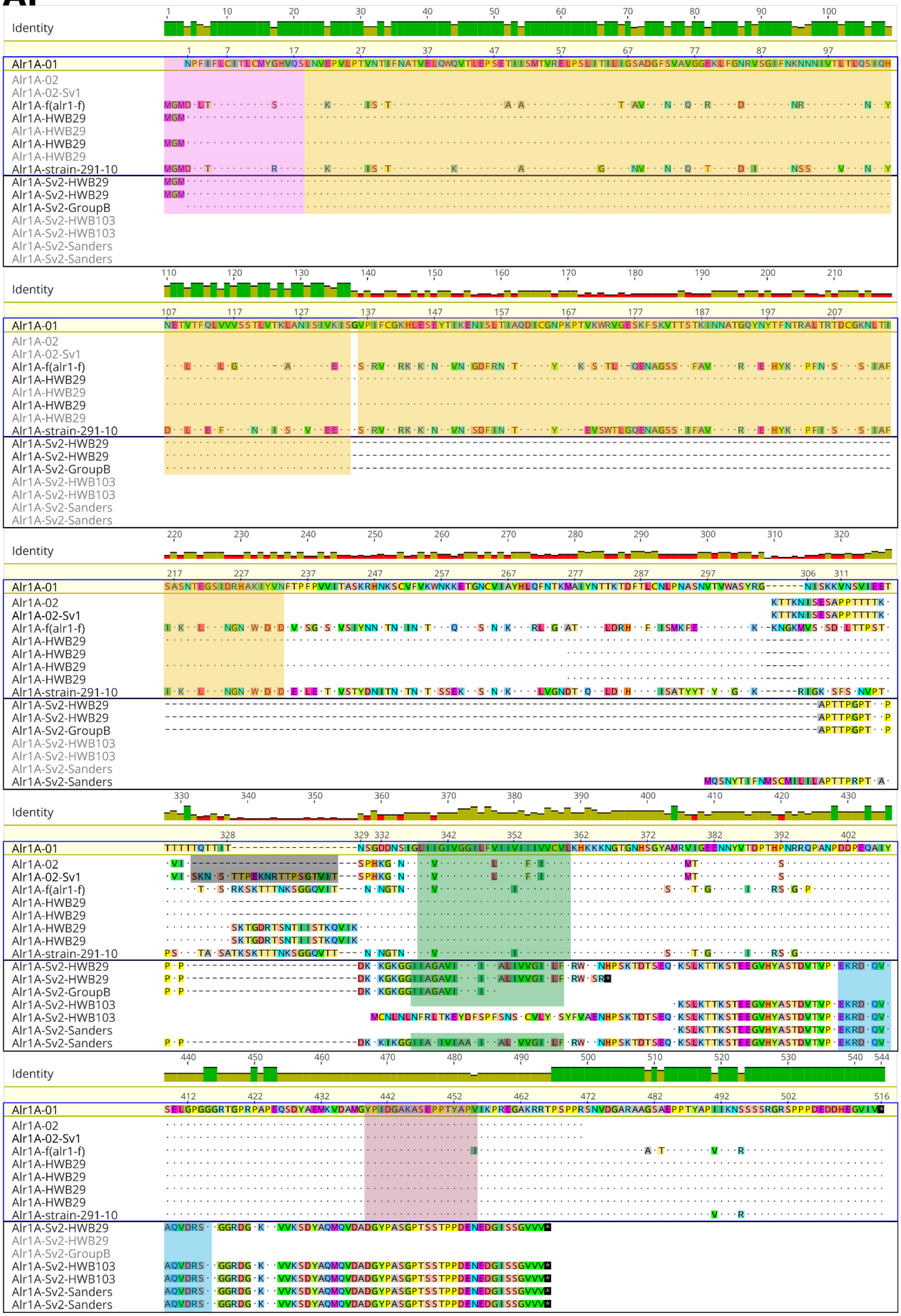
Identity
Alr1A-01
Alr1A-02
Alr1A-02-Sv1
Alr1A-f(alr1-f)
Alr1A-HWB29
Alr1A-strain-291-10
Alr1A-Sv2-HWB29
Alr1A-Sv2-GroupB
Alr1A-Sv2-HWB103
Alr1A-Sv2-Sanders

**B.**

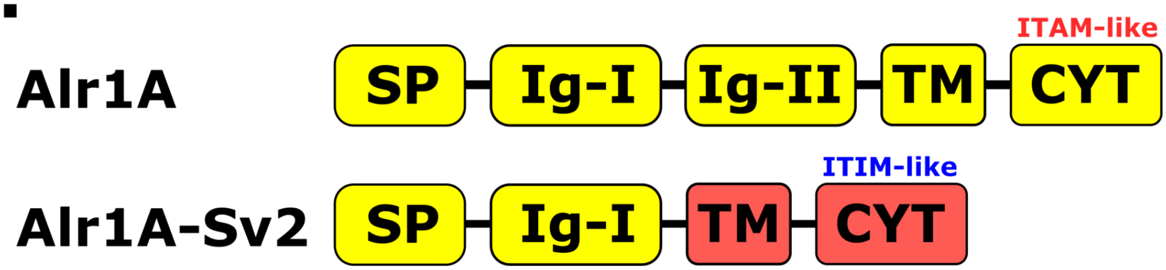


**Fig. S4 (A) Alignment of Alr1A and Alr1A-Sv2.** Two alleles (Alr1A-01 and Alr1A-02) and two splicing variants were detected in our backcross population for the *alr1A* gene. The Alr1A-Sv1 splicing variant was detected for allele 2 and corresponds to a small insertion (highlighted in black) before the TM-domain. The Alr1A-Sv2 splicing variant shares the signal peptide and the first Ig-domain with the Alr1A protein. However, the Alr1A-Sv2 protein does not have the second Ig-domain, and it has different transmembrane and cytoplasmic regions compared with the Alr1A protein. The cytoplasmic region of the Alr1A-Sv2 protein has a putative ITIM-like motif instead of an ITAM-like motif as in the Alr1A protein. Signal peptides, Ig-domains, TM-Domains, ITAM-like and ITIM-like motifs are highlighted in pink, yellow, green, red, and blue, respectively. Dots in the alignment indicate residue conservation with respect to the reference sequence. Reported *alr1-f* allele (renamed *alr1A-f*) was included in the alignment. **(B) Summary of Alr1A and Alr1A-Sv2 protein architecture.** Alr1A and Alr1A-Sv2 proteins share the signal peptide and first Ig-domain, but Alr1A-Sv2 does not have a second Ig-domain and has different transmembrane and cytoplasmic regions compared with the Alr1A protein. Note that the Alr1A-Sv2 protein is a putative inhibitory receptor in contrast to the Alr1A protein that is a putative activating receptor. SP: Signal peptide, Ig-I and Ig-II: First and second Immunoglobulin domains, TM: Transmembrane domain, CYT: Cytoplasmic region

**A.**

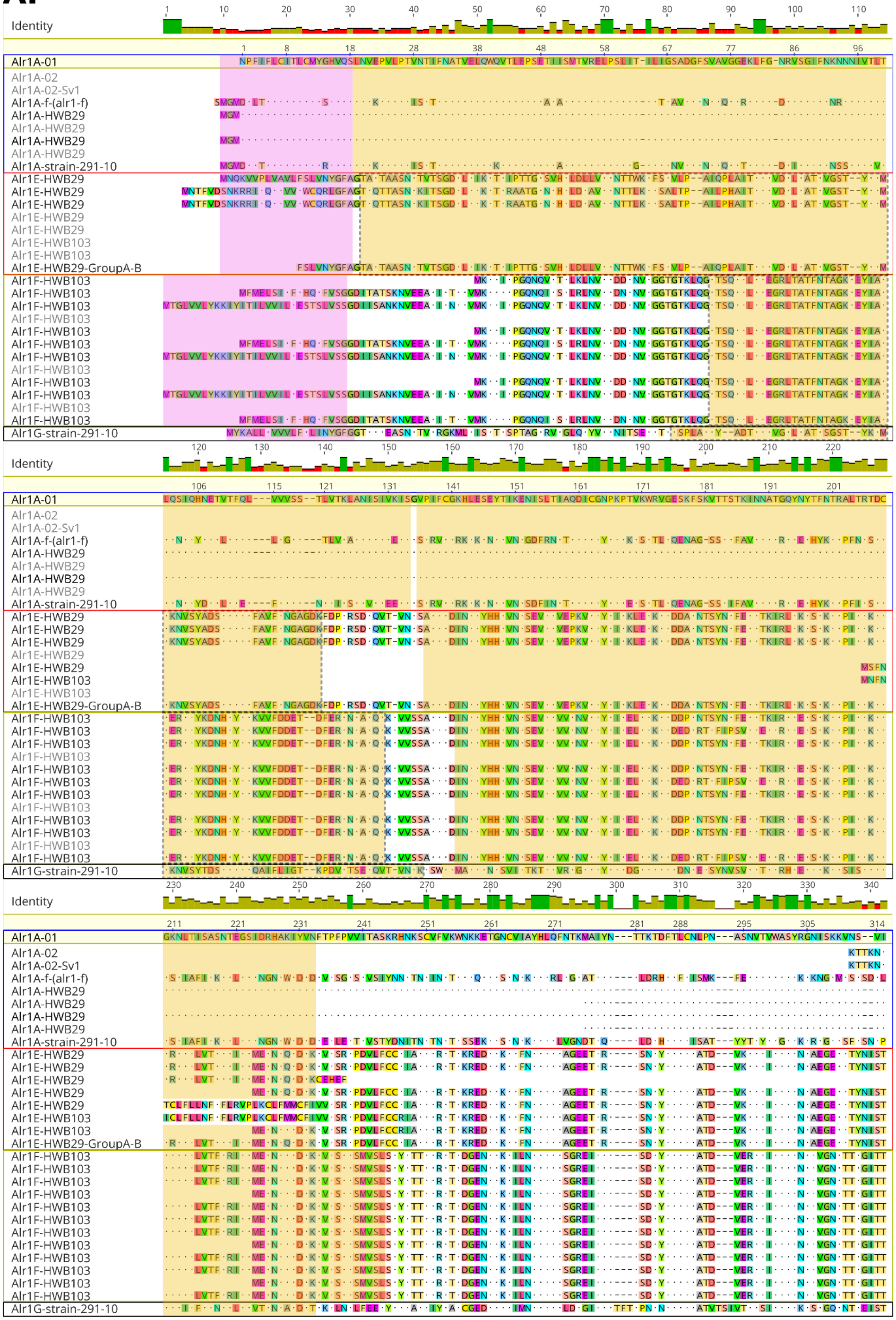
Identity
Alr1A-01
Alr1A-02
Alr1A-02-Sv1
Alr1A-f-(alr1-f)
Alr1A-HWB29
Alr1A-strain-291-10
Alr1E-HWB29
Alr1E-HWB103
Alr1E-HWB29-GroupA-B
Alr1F-HWB103
Alr1G-strain-291-10

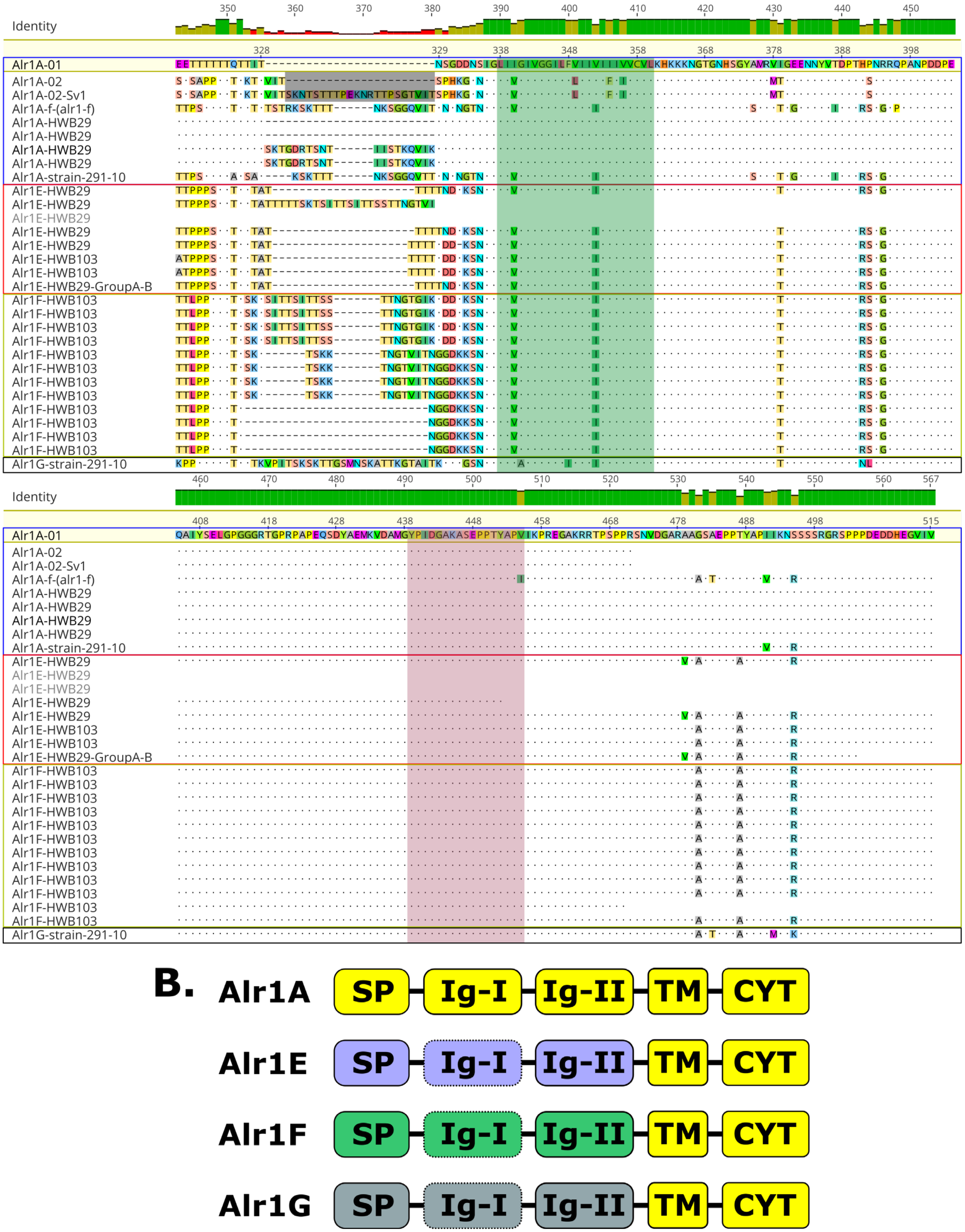
Identity
Alr1A-01
Alr1A-02
Alr1A-02-Sv1
Alr1A-f-(alr1-f)
Alr1A-HWB29
Alr1A-strain-291-10
Alr1E-HWB29
Alr1E-HWB103
Alr1E-HWB29-GroupA-B
Alr1F-HWB103
Alr1G-strain-291-10
B.
Alr1A
Alr1E
Alr1F
Alr1G
SP
Ig-I
Ig-II
TM
CYT

**Fig. S5 (A) Alignment of Alr1A, Alr1E, Alr1F and Alr1G proteins.** These sequences show multiple differences in the extracellular region (Signal peptide and two Ig-domains). However, these proteins share the same transmembrane domain and cytoplasmic region. Signal peptides, Ig-domains, TM-Domains, and ITAM-like motif are highlighted in pink, yellow, green, and red, respectively. Hypothetical first Ig-domains of the Alr1E, Alr1F and Alr1G proteins are indicated by yellow rectangles with dotted lines. Alr1A protein has a splicing variant (Alr1A-02-Sv1) that is highlighted in the black rectangle before TM-domain. **(B) Summary of the architecture of Alr1A, Alr1E, Alr1F and Alr1G proteins.** These proteins share the same transmembrane and cytoplasmic regions (highlighted in yellow), but they have different signal peptides and Ig-domains, represented by different colors for each gene. SP: Signal peptide; Ig-I and Ig-II: First and second Immunoglobulin domains; TM: Transmembrane domain; CYT: cytoplasmic region

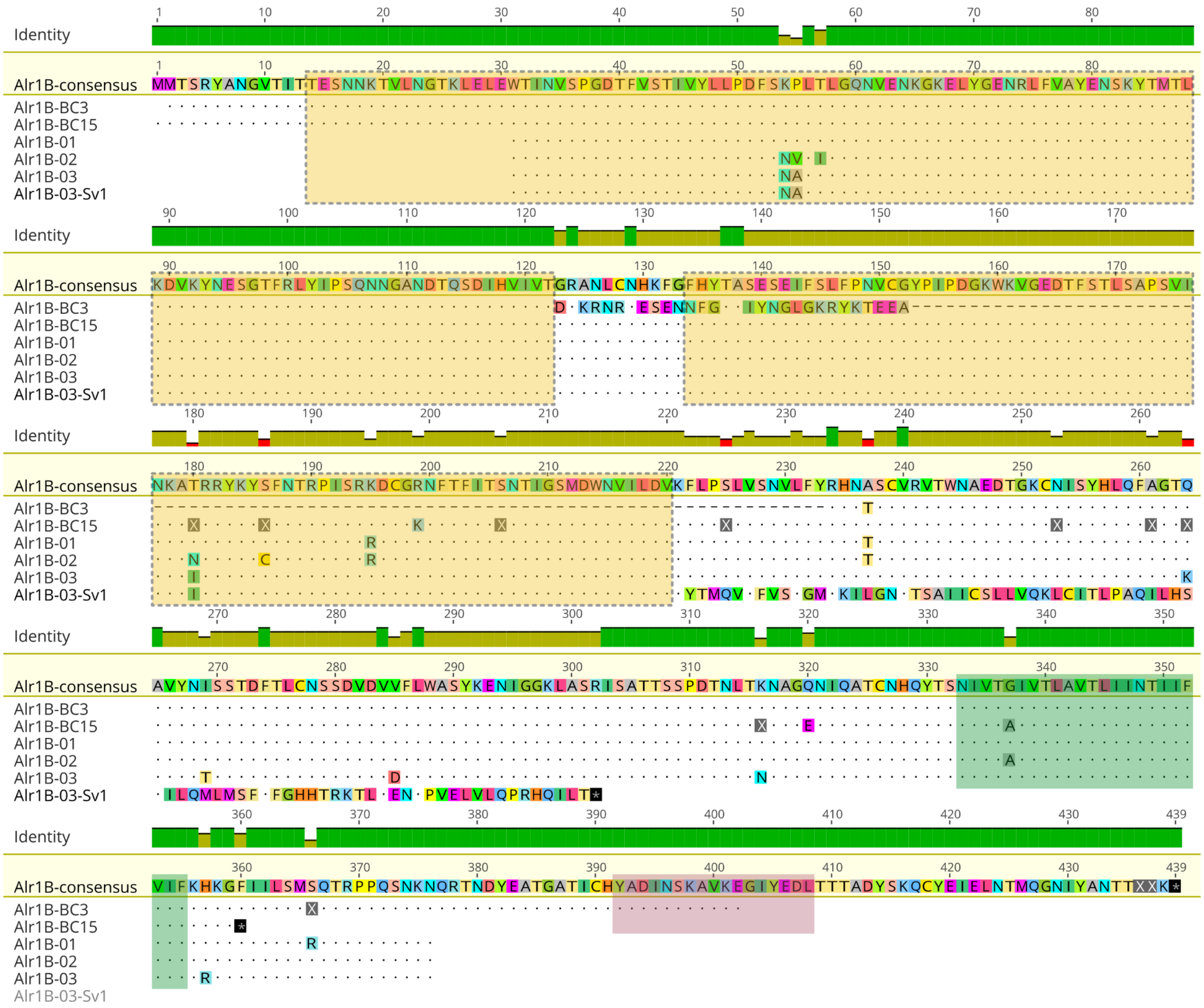


**Fig. S6 Alignment of Alr1B protein.** Predicted Alr1B proteins from the genomes of individuals BC-3 and BC-15 are compared with the sequences cloned from our backcross population. An Alr1B-consensus sequence was created with these sequences and used as reference. Alr1B transcripts could not be identified from the transcriptomes analyzed in this study. A splicing variant (Alr1B-03-Sv1) of allele 3 was identified in the individual HWB53, which has a premature stop codon before the TM domain. The signal peptide for the Alr1B protein was not detected in our sequences. Two Ig-domains were identified by Pfam software with non-significant e-values. These hypothetical Ig-domains are highlighted in yellow rectangles with dotted lines. TM domain and ITAM motif are highlighted in green and red rectangles, respectively

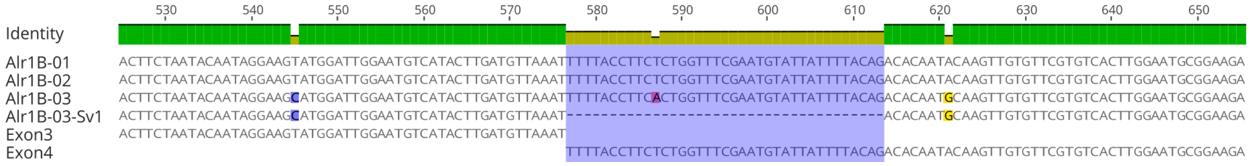


**Fig. S7 Splicing variant of the *alr1B* gene.** A splicing variant was observed for allele 3 of the *alr1B* gene. All sequences have a full-length exon 3, but the splicing variant (Alr1B-03-Sv1) of allele 3 lost the first region of exon 4 (37bp, highlighted in blue). This deletion creates a premature stop codon before the transmembrane domain, as shown in the previous figure

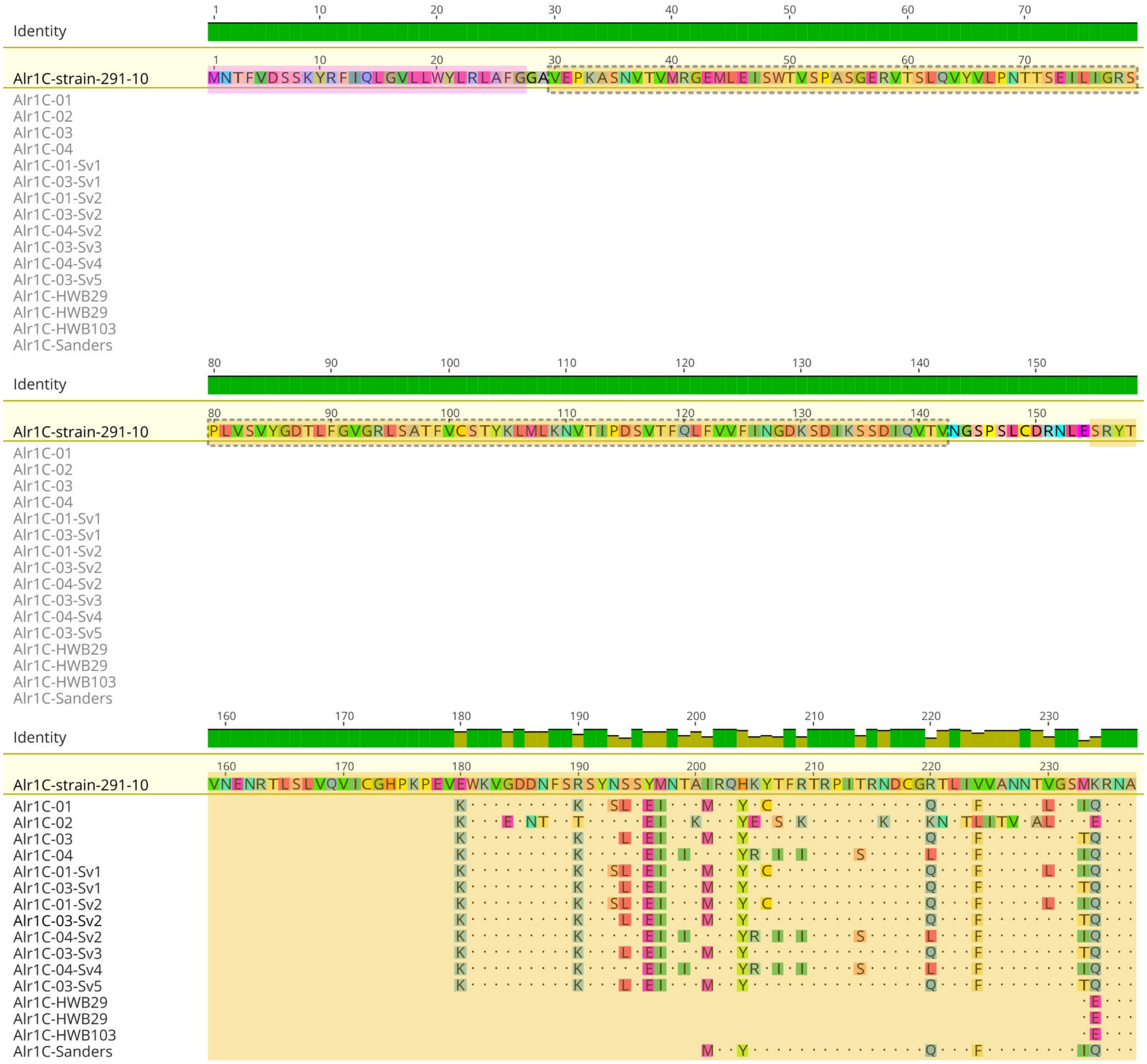

Identity
Alr1C-strain-291-10
Alr1C-01
Alr1C-02
Alr1C-03
Alr1C-04
Alr1C-01-Sv1
Alr1C-03-Sv1
Alr1C-01-Sv2
Alr1C-03-Sv2
Alr1C-04-Sv2
Alr1C-03-Sv3
Alr1C-04-Sv4
Alr1C-03-Sv5
Alr1C-HWB29
Alr1C-HWB29
Alr1C-HWB103
Alr1C-Sanders

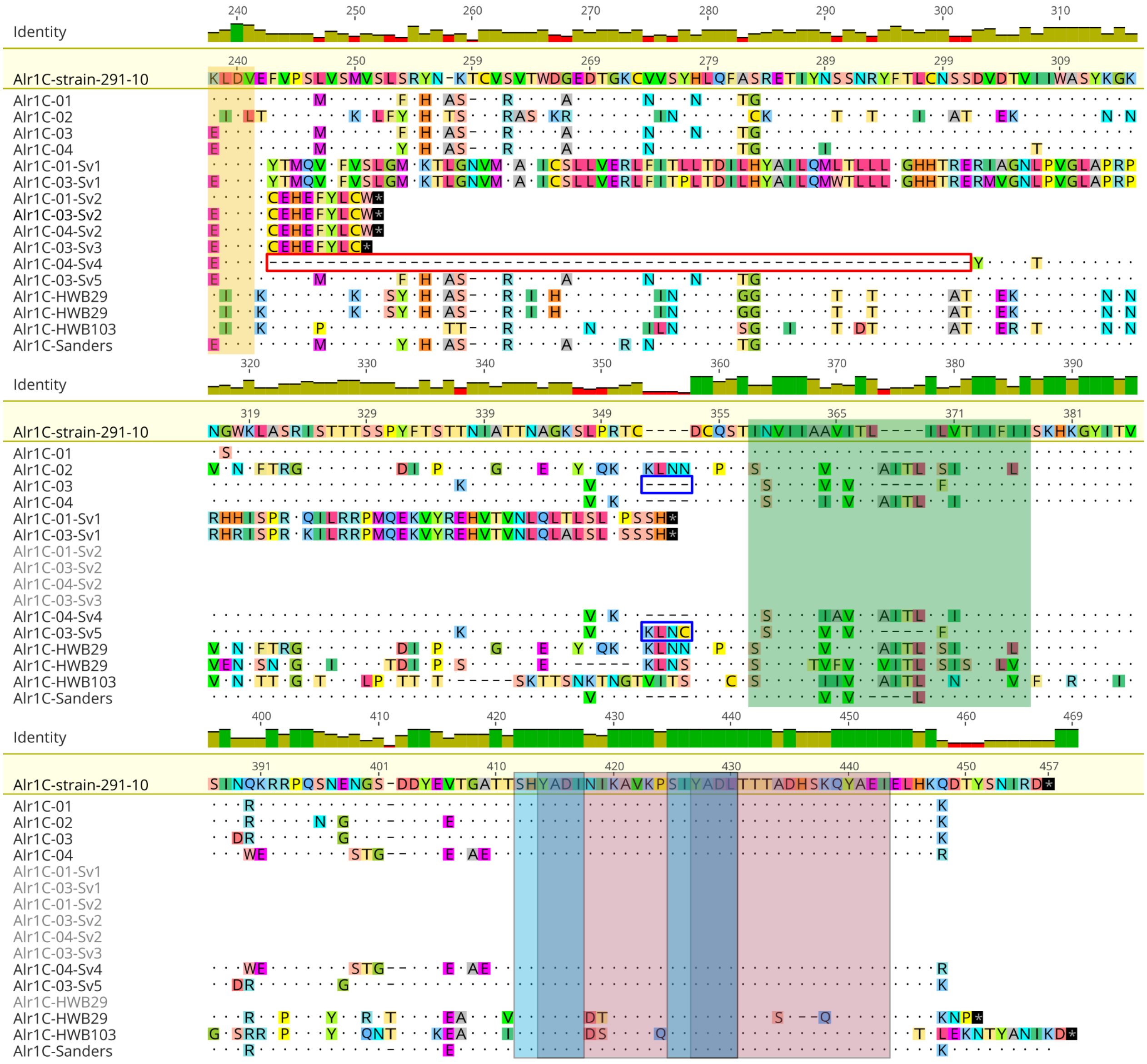

**Fig. S8 Alignment of Alr1C protein.** Alr1C sequences isolated from our backcross population (Alr1C-01, -02, -03 and -04) and the transcriptomes analyzed in this study are compared. The putative first region of Alr1C protein was only identified in the transcriptome of strain 291-10. Five splicing variants (Sv1 to Sv5) of Alr1C protein were identified in our backcross population. Sv1, Sv2 and Sv3 have premature stop codons before the transmembrane domain. The cytoplasmic region of Alr1C protein has two putative ITIM motifs, or two overlapping putative ITAM motifs. Signal peptide, Ig-domains, TM-Domain, ITAM and ITIM motifs are highlighted in pink, yellow, green, red, and blue, respectively. The first Ig-domain was predicted by Pfam software with a non-significant e-value, and it is represented by a yellow rectangle with dotted lines. The second Ig-domain was predicted normally with Interproscan software

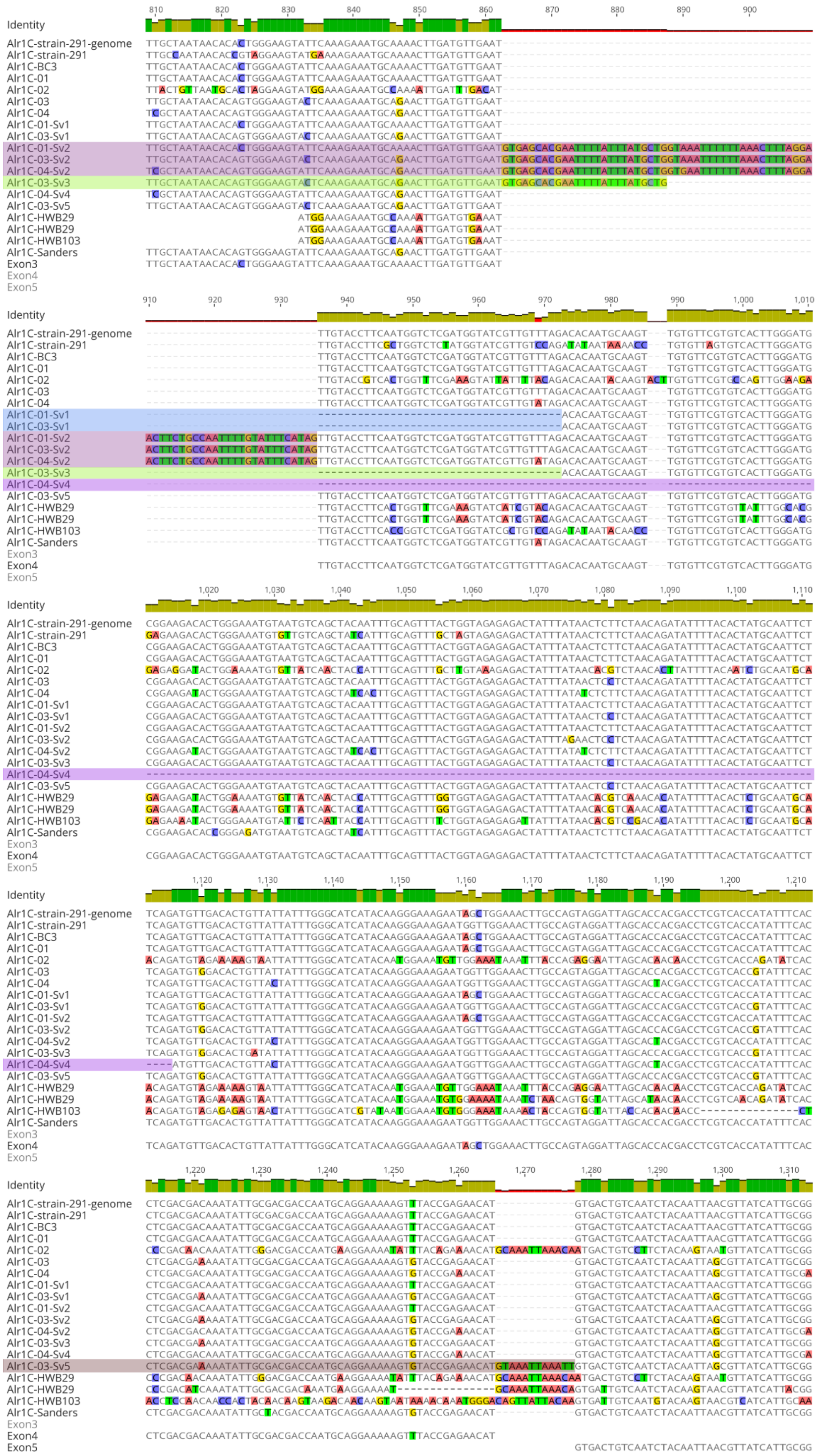

Identity
Alr1C-strain-291-genome
Alr1C-strain-291
Alr1C-BC3
Alr1C-01
Alr1C-02
Alr1C-03
Alr1C-04
Alr1C-01-Sv1
Alr1C-03-Sv1
Alr1C-01-Sv2
Alr1C-03-Sv2
Alr1C-04-Sv2
Alr1C-03-Sv3
Alr1C-04-Sv4
Alr1C-03-Sv5
Alr1C-HWB29
Alr1C-HWB29
Alr1C-HWB103
Alr1C-Sanders
Exon3
Exon4
Exon5

**Fig. S9 Splicing variants of the *alr1C* gene.** Sequences of the *alr1C* gene isolated from the backcross population (Alr1C-01, -02, -03 and -04) and the transcriptomes are compared. In addition, the sequences of exons 3, 4 and 5 are shown at the bottom of the alignment. Five splicing variants (Sv1 to Sv5) were detected for the *alr1C* gene. Splicing variant 1 (Sv1) of the *alr1C* gene loses a fragment of exon 4 (37bp, highlighted in blue). Sv2 retains a fragment of intron 3 (73bp, highlighted in red). Sv3 retains a fragment of intron 3 (25bp) and loses a fragment of exon 4 (37bp, highlighted in green). These three splicing variants have premature stop codons before the transmembrane domain, as shown in the previous figure. Sv4 loses a fragment of exon 4 (177bp, highlighted in pink), and Sv5 retains a fragment of intron 4 (12bp, highlighted in gray). These two splicing variants encode transmembrane proteins, as shown in the previous figure

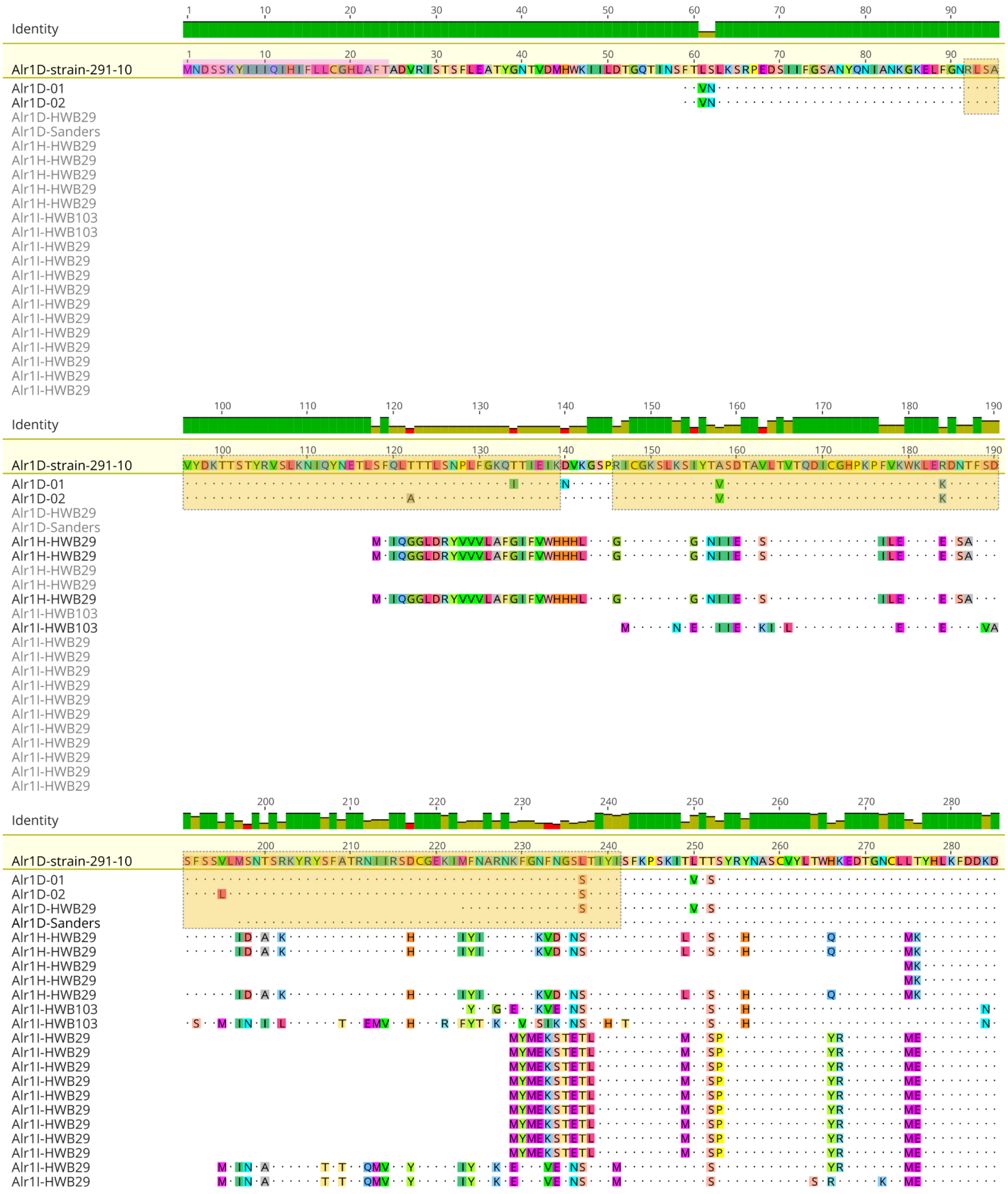

Identity
Alr1D-strain-291-10
Alr1D-01
Alr1D-02
Alr1D-HWB29
Alr1D-Sanders
Alr1H-HWB29
Alr1I-HWB103
Alr1I-HWB29

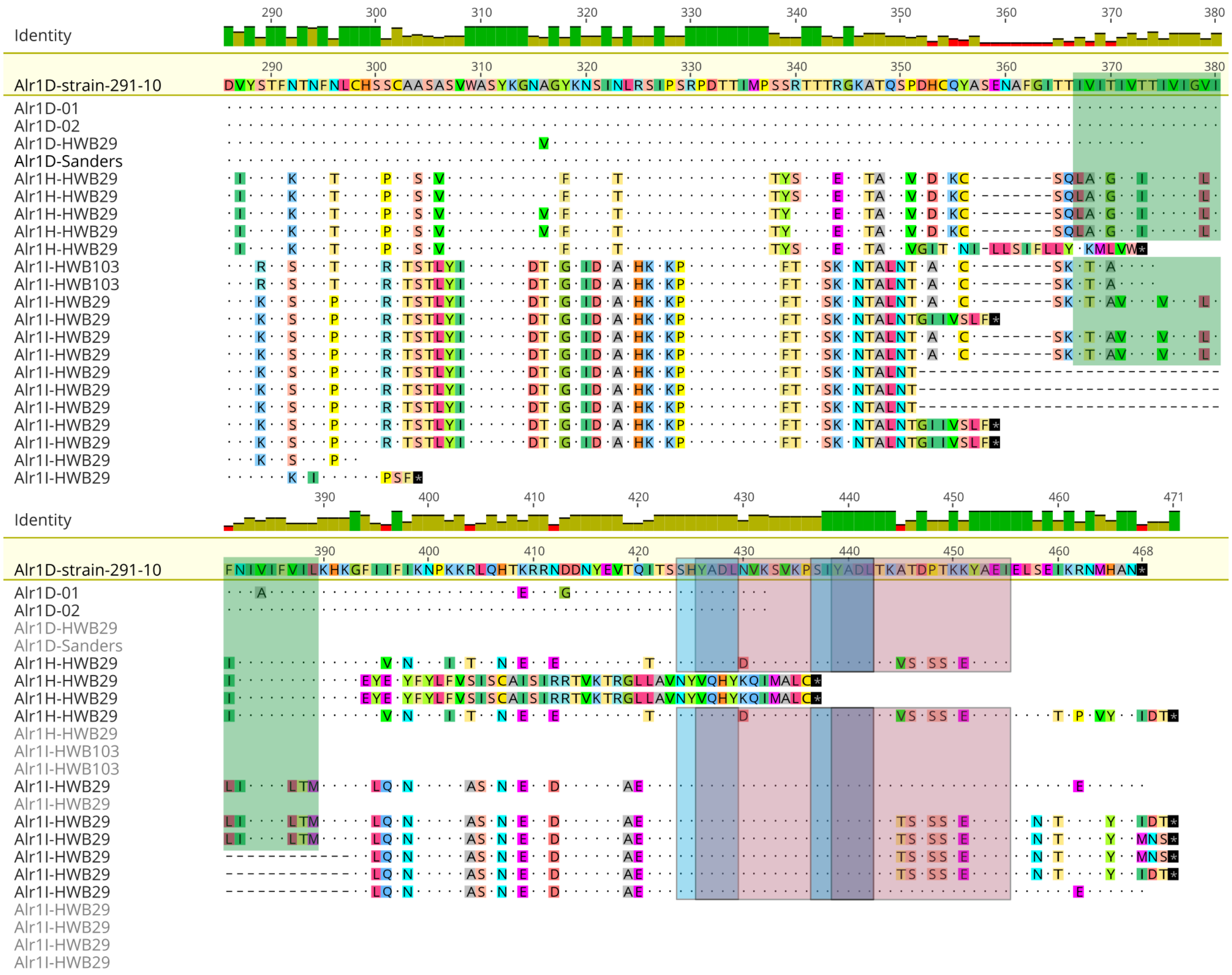

**Fig. S10 Alignment of Alr1D, Alr1H and Alr1I proteins.** Two alleles of Alr1D proteins were identified in our backcross population (Alr1D-01 and -02). Alr1D transcripts were isolated from HWB29, Sanders and strain 291-10 transcriptomes. Putative *alr1H* and *alr1I* genes were isolated from the HWB29 transcriptome. Transcripts for the *alr1I* gene were found in the HWB103 transcriptome. Two hypothetical Ig-domains were predicted in the extracellular region of Alr1D protein by Pfam software with non-significant e-values. The cytoplasmic regions of Alr1D, Alr1H and Alr1I proteins have two putative ITIM motifs, or two overlapping putative ITAM motifs. Signal peptide, Ig-domains, TM-Domain, ITAM and ITIM motifs are highlighted in pink, yellow (with dotted lines), green, red, and blue, respectively

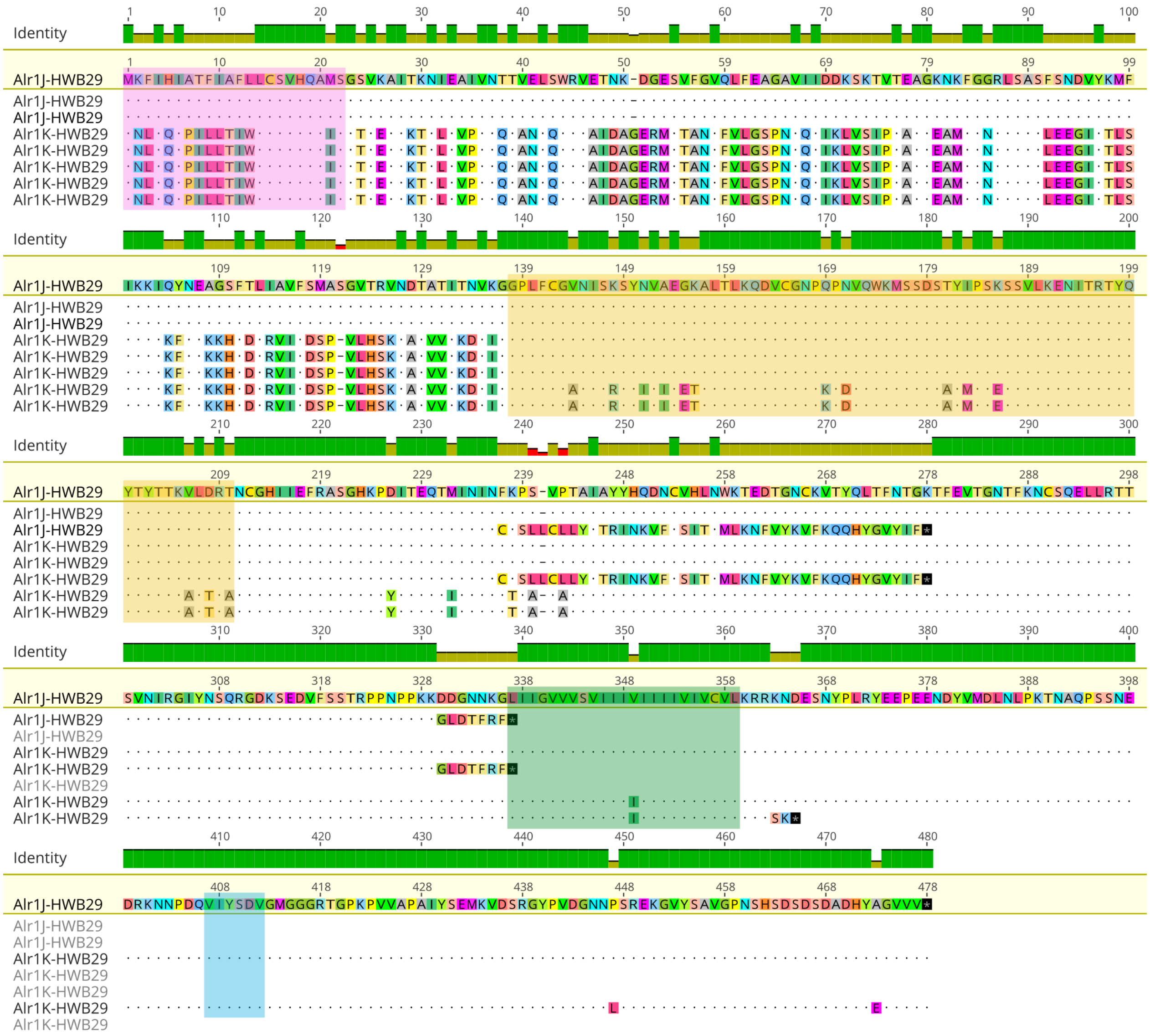

**Fig. S11 Alignment of the Alr1J and Alr1K proteins.** Alr1J and Alr1K proteins were identified in the transcriptome of the individual HWB29. These sequences show differences in the N-terminal region, but they share the sequence for the Ig-domain, transmembrane and cytoplasmic regions. These two proteins have an ITIM motif in the cytoplasmic region. Signal peptide, Ig-domain, TM-Domain and ITIM motifs are highlighted in pink, yellow, green, and blue, respectively

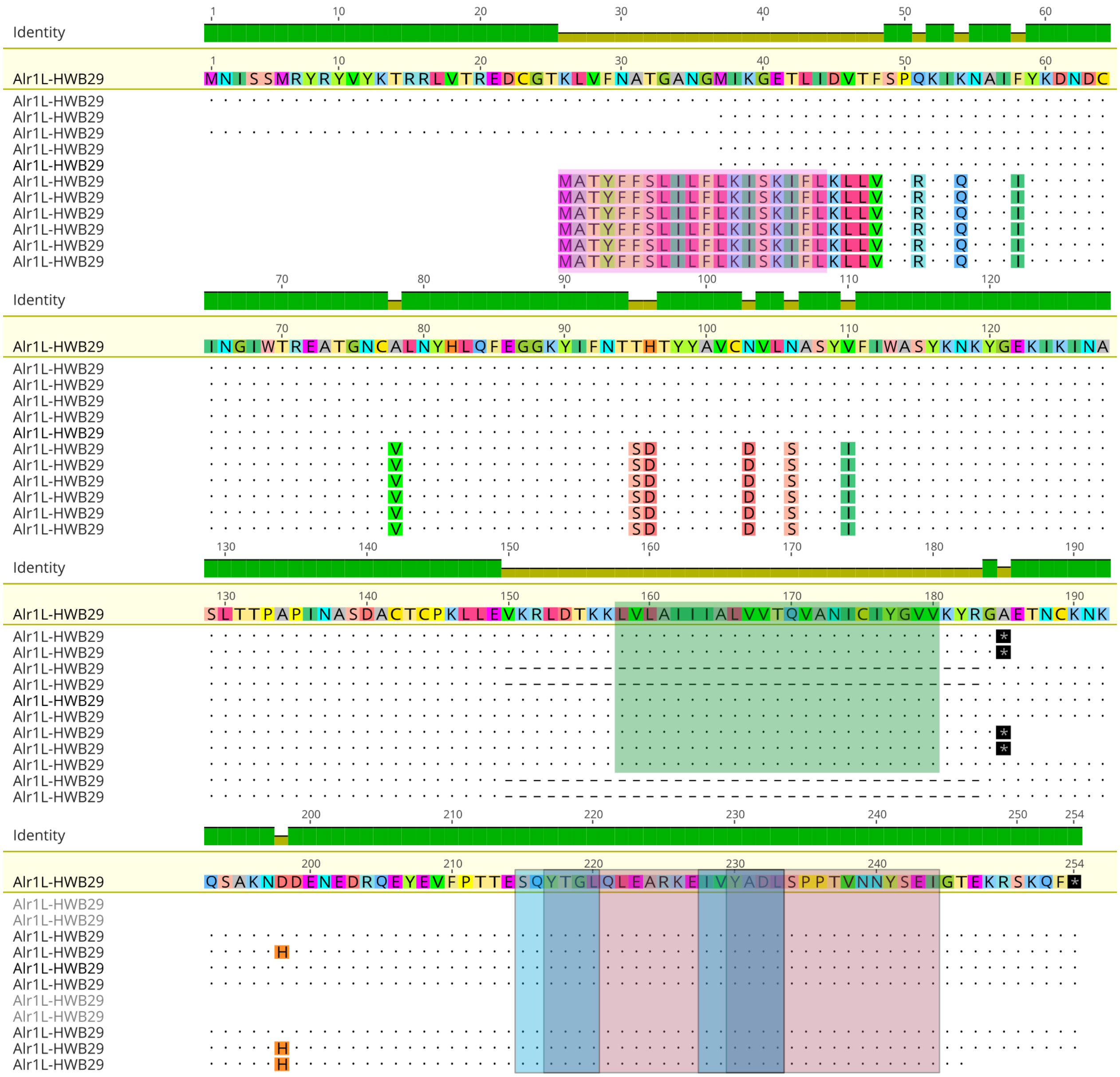

**Fig. S12 Alignment of the Alr1L protein.** Alr1L protein was identified in the transcriptome of the individual HWB29. The cytoplasmic region of Alr1L protein has two putative ITIM motifs, or two overlapping putative ITAM motifs. Signal peptide, TM-Domain, ITAM and ITIM motifs are highlighted in pink, green, red, and blue, respectively

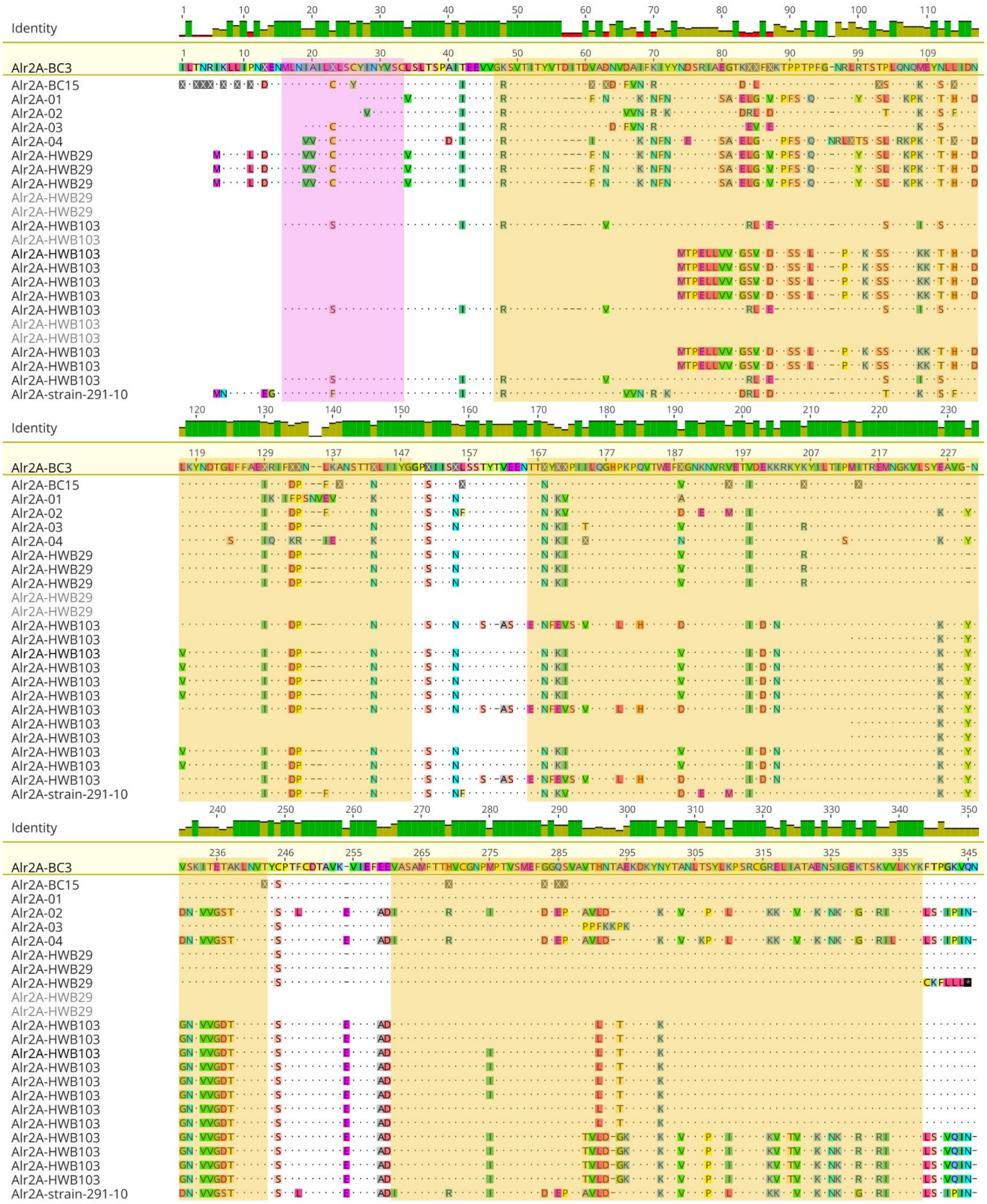
Identity
Alr2A-BC3
Alr2A-BC15
Alr2A-01
Alr2A-02
Alr2A-03
Alr2A-04
Alr2A-HWB29
Alr2A-HWB103
Alr2A-strain-291-10

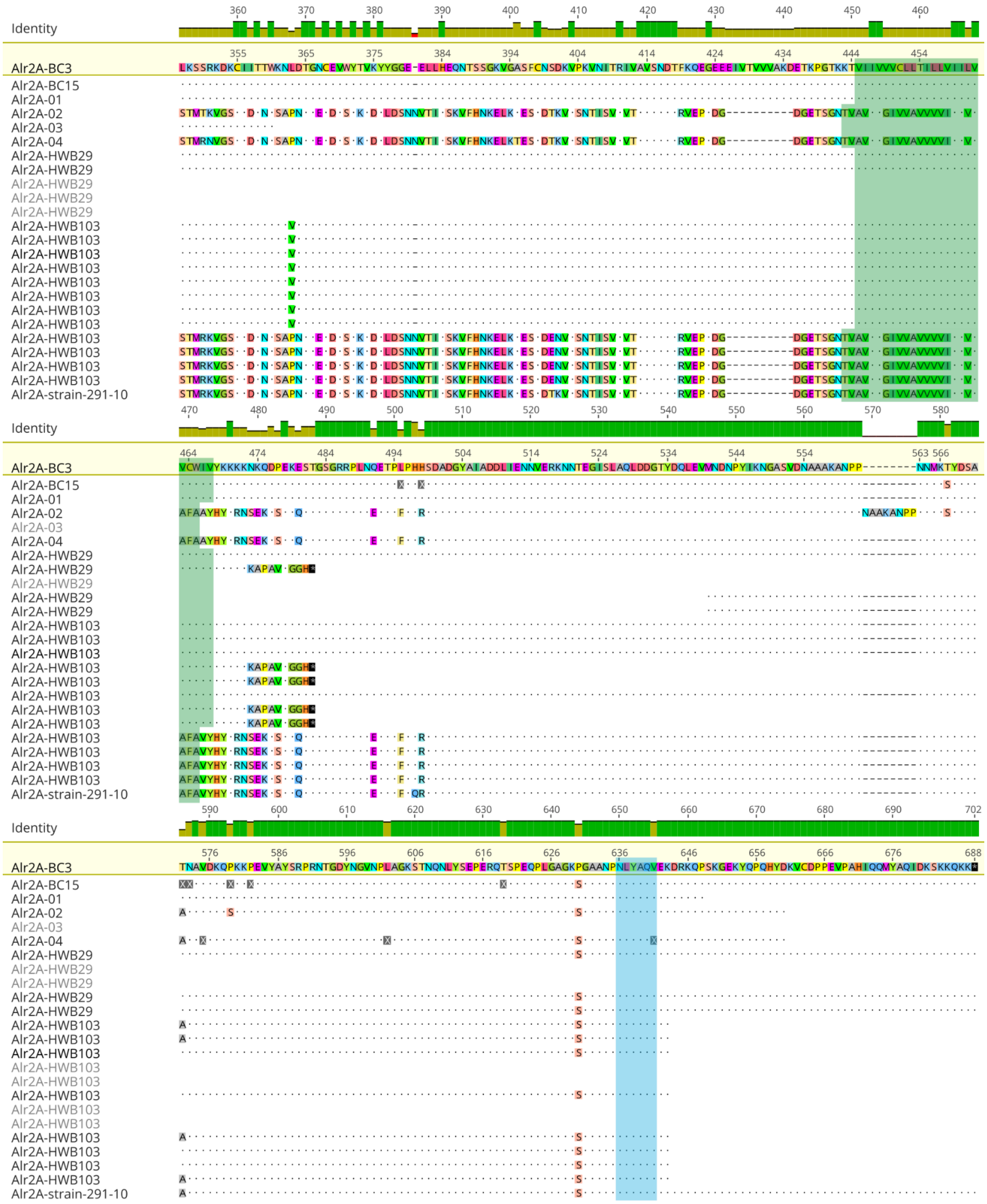
Identity
Alr2A-BC3
Alr2A-BC15
Alr2A-01
Alr2A-02
Alr2A-03
Alr2A-04
Alr2A-HWB29
Alr2A-HWB103
Alr2A-strain-291-10

**Fig. S13 Alignment of the Alr2A protein.** Alr2A sequences isolated from BC-3 and BC-15 genomes, our backcross population (Alr2A-01, -02, -03 and -04), and HWB29, HWB103 and strain 291-10 transcriptomes are compared. Alleles Alr2A-02 and Alr2A-03 are more similar to each other in the signal peptide and the first Ig-domain. Meanwhile, Alr2A-02 and Alr2A-04 are more similar to each other in the second and third Ig-domains, and the transmembrane and cytoplasmic regions. Signal peptide, Ig-domains, TM-Domain, and ITIM-like motif are highlighted in pink, yellow, green, and blue, respectively. X: Non-determined residues

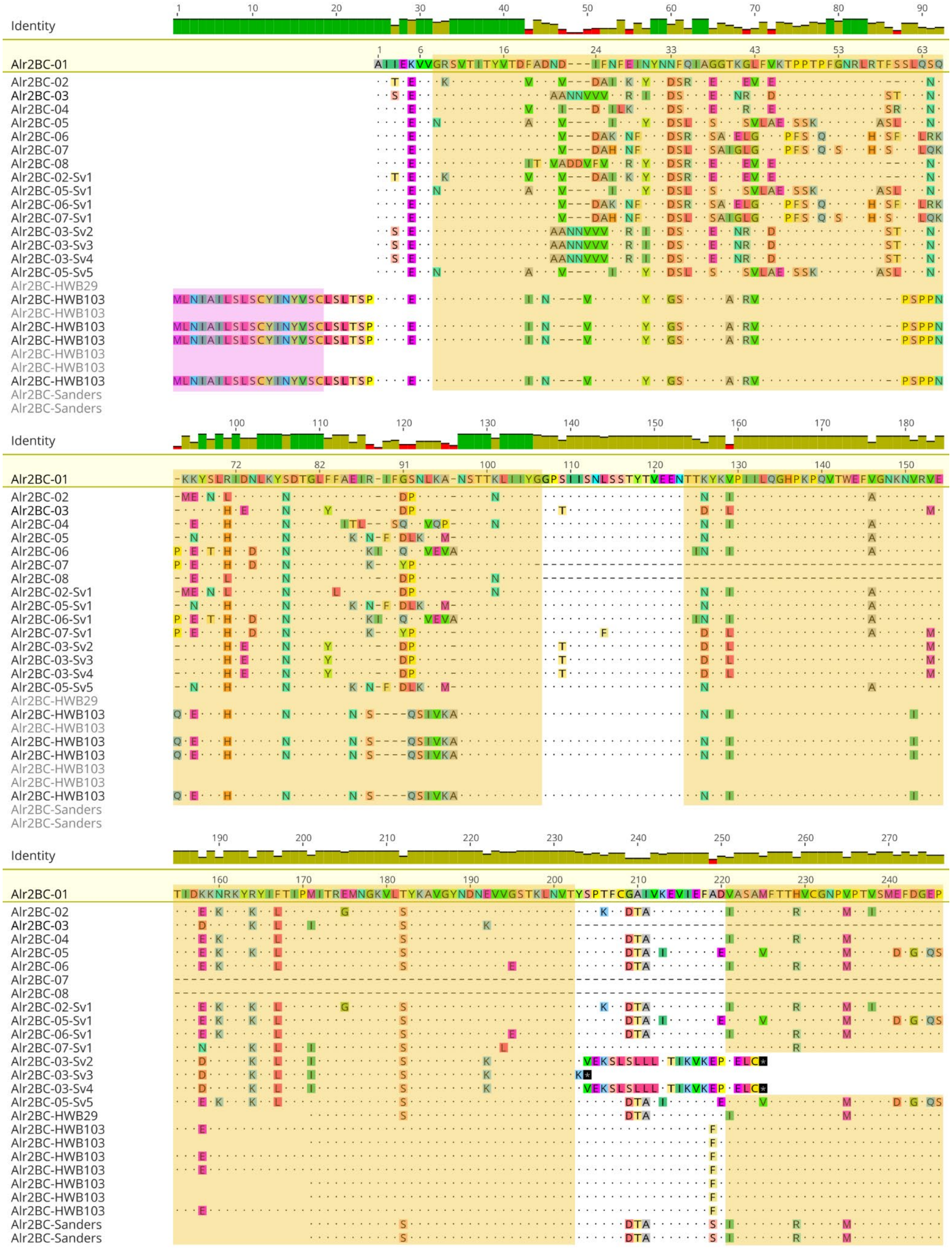
Identity
Alr2BC-01
Alr2BC-02
Alr2BC-03
Alr2BC-04
Alr2BC-05
Alr2BC-06
Alr2BC-07
Alr2BC-08
Alr2BC-02-Sv1
Alr2BC-05-Sv1
Alr2BC-06-Sv1
Alr2BC-07-Sv1
Alr2BC-03-Sv2
Alr2BC-03-Sv3
Alr2BC-03-Sv4
Alr2BC-05-Sv5
Alr2BC-HWB29
Alr2BC-HWB103
Alr2BC-Sanders

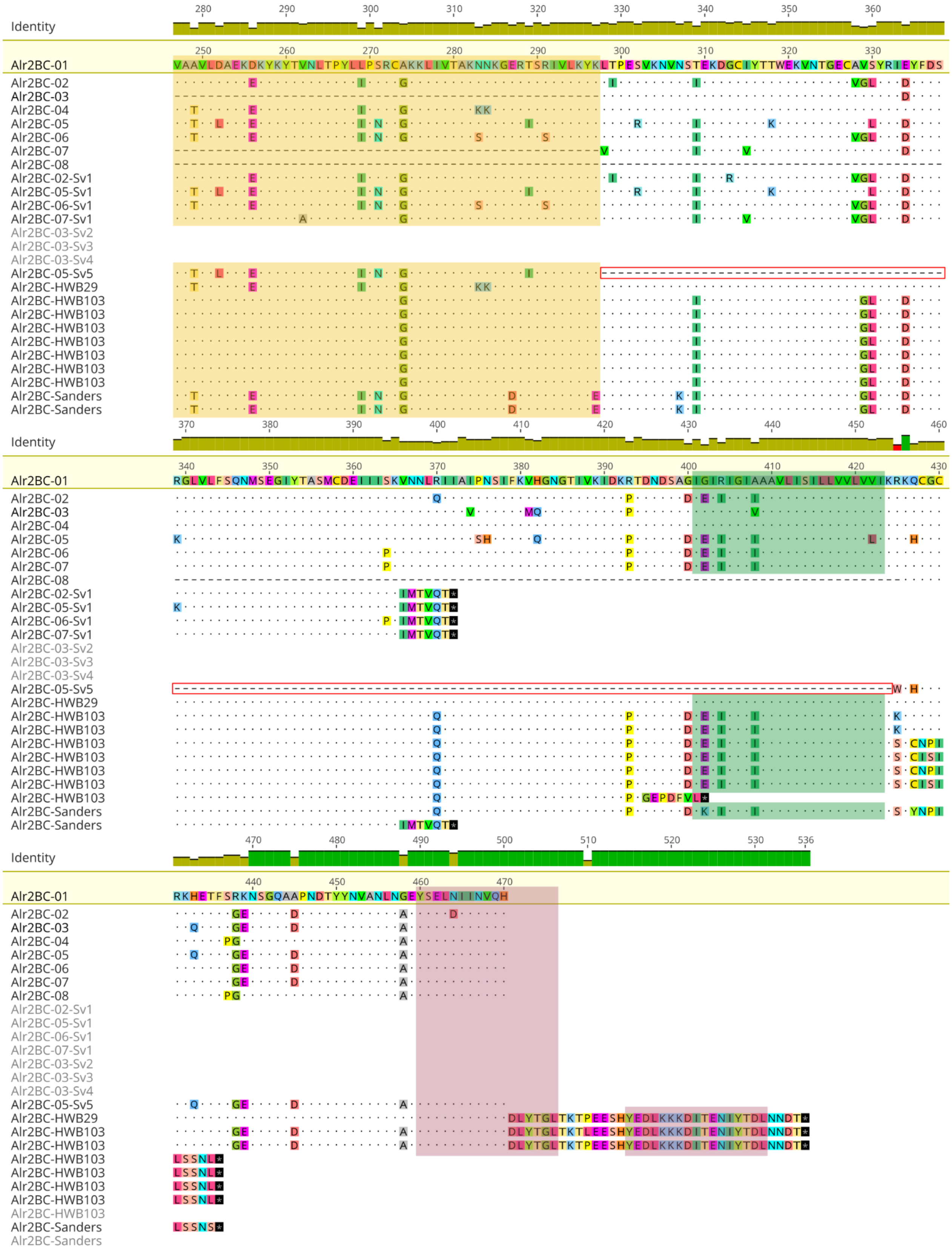
Identity
Alr2BC-01
VAAVLDAEKDKYKYTVNLTPYLLPSRCAKKLIVTAKNNKGERTSRIVLKYKLTPESVKNVNSTEKDGCIYTTWEKVNTGECAVSYRIEYFDS
Alr2BC-02
Alr2BC-03
Alr2BC-04
Alr2BC-05
Alr2BC-06
Alr2BC-07
Alr2BC-08
Alr2BC-02-Sv1
Alr2BC-05-Sv1
Alr2BC-06-Sv1
Alr2BC-07-Sv1
Alr2BC-03-Sv2
Alr2BC-03-Sv3
Alr2BC-03-Sv4
Alr2BC-05-Sv5
Alr2BC-HWB29
Alr2BC-HWB103
Alr2BC-Sanders
Identity
Alr2BC-01
RGLVLFSQNMSEGIYTASMCDEIIISKVNNLRIIAIPNSIFKVHGNGTIVKIDKRTDNDSAGIGIRIGIAAAVLISILLVVLVVIKRKQCGC
Identity
Alr2BC-01
RKHETFSRKNSGQAAPNDTYYNVANLNGEYSELNIINVQH
DLYTGLTKTPEESHYEDLKKKDITENIYTDLNNDT*
LSSNL*
LSSNS*

**Fig. S14 Alignment of the Alr2BC proteins.** Eight Alr2BC alleles were identified in the backcross population (Alr2BC-01 to Alr2BC-08). Five splicing variants (Alr2BC-Sv1 to Alr2BC-Sv5) were detected for the Alr2BC proteins in our backcross population. Alr2BC-Sv1, -Sv2, -Sv3 and -Sv4 splicing variants have premature stop codons before the transmembrane domain. Alr2BC-Sv1 encodes a putative secreted protein with three Ig-domains, while Alr2BC-Sv2, -Sv3 and -Sv4 encode putative secreted proteins with two Ig-domains. Alr2BC-Sv5 skips the stem region and transmembrane domain (highlighted in the empty red rectangle), which encodes a putative secreted protein with three Ig-domains. The cytoplasmic region of the Alr2BC protein has two putative ITAM motifs. Signal peptide, Ig-domains, TM-Domain and ITAM motifs are highlighted in pink, yellow, green, and red, respectively

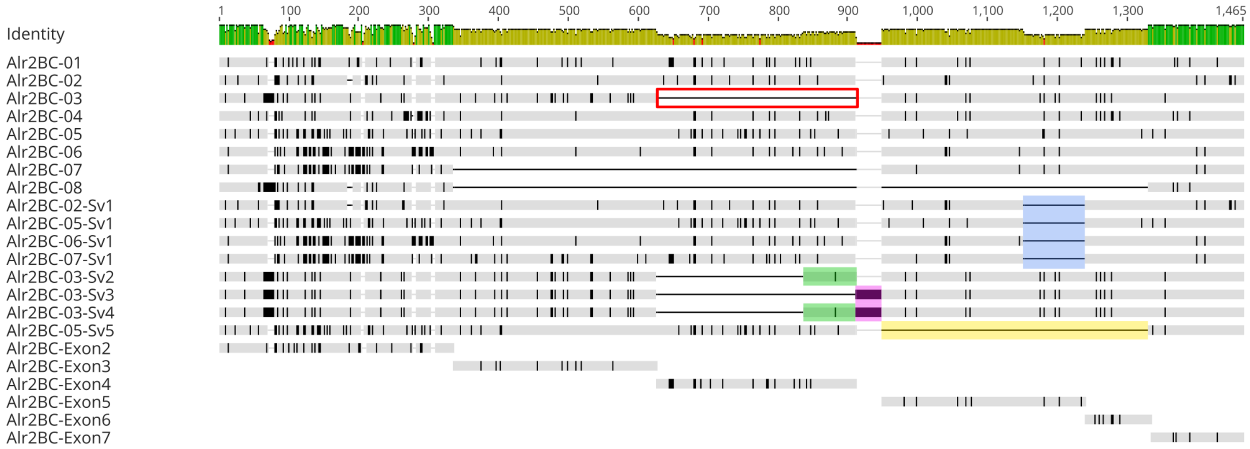


**Fig. S15 Alignment of splicing variants of the *alr2BC* genes.** Five splicing variants (Alr2BC-Sv1 to Alr2BC-Sv5) were detected for the Alr2BC proteins in our backcross population. Alr2BC-Sv1 was detected for alleles 2, 5, 6 and 7. This consists of a deletion in the exon 5 (highlighted in blue). Alr2BC-Sv2, -Sv3 and -Sv4 were detected for allele 3. The "full-length" allele 3 skips the exon 4 (empty red rectangle). Alr2BC-Sv2 retains a fragment of exon 4 (highlighted in green). Alr2BC-Sv3 retains a fragment of intron 4 (highlighted in pink). Alr2BC-Sv4 retains a fragment of exon 4 and intron 4 (highlighted in green and pink). Alr2BC-Sv5 was detected for allele 5. This variant loses the exon 5 and most of the exon 6 (highlighted in yellow). Variable nucleotide positions are noted as vertical black lines in the alignment

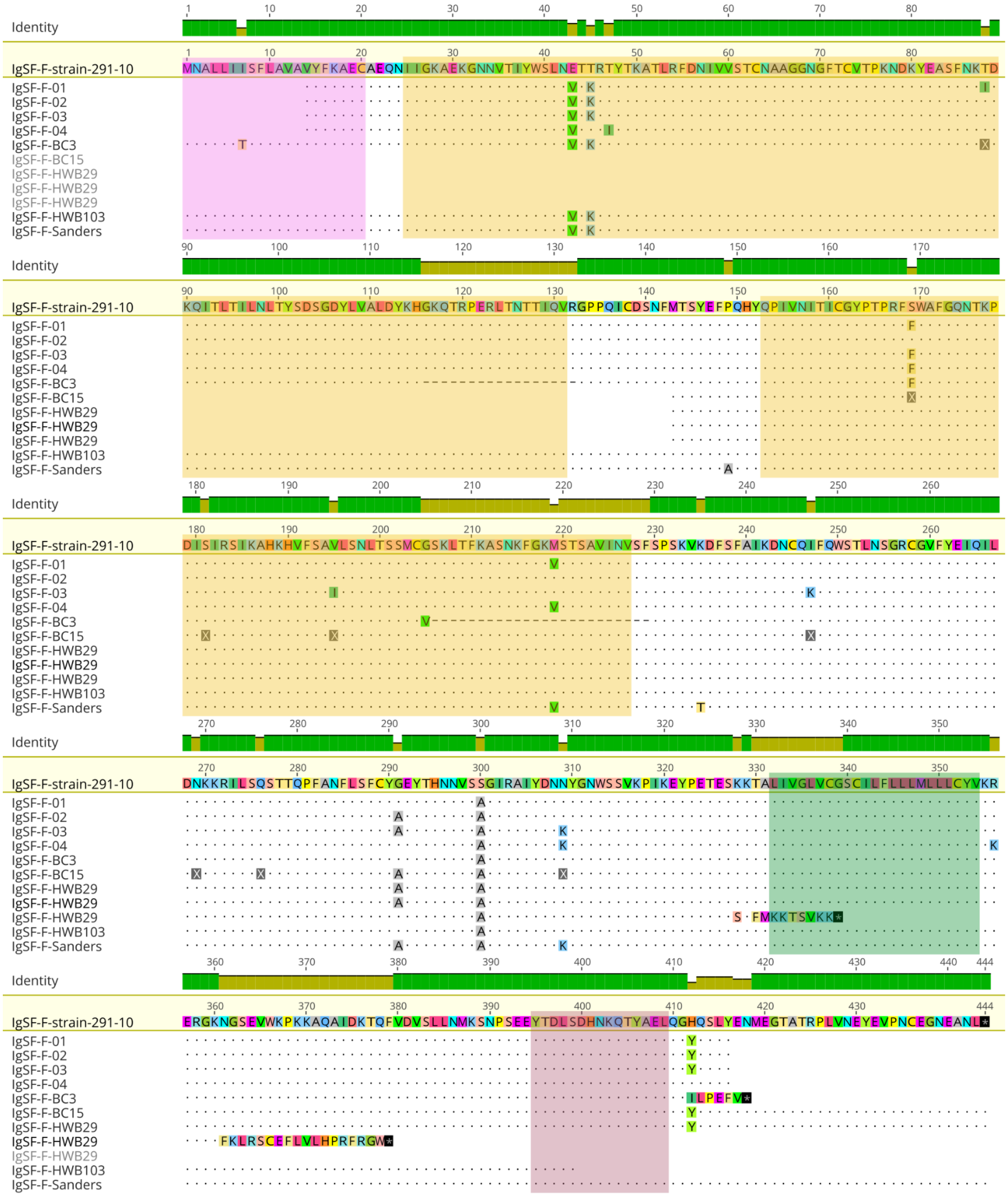

**Fig. S16 Alignment of the IgSF-F protein.** Alleles of the *IgSF-F* gene isolated from our backcross population (IgSF-F-01 to IgSF-F-04), sequences predicted from the genomes of the individuals BC-3 and BC-15, and the sequences from the transcriptomes analyzed (HWB29, HWB103,

Sanders and strain 291-10) were compared. The cytoplasmic region of the IgSF-F protein has an ITAM motif. Signal peptide, Ig-domains, TM-Domain and the ITAM motif are highlighted in pink, yellow, green, and red, respectively

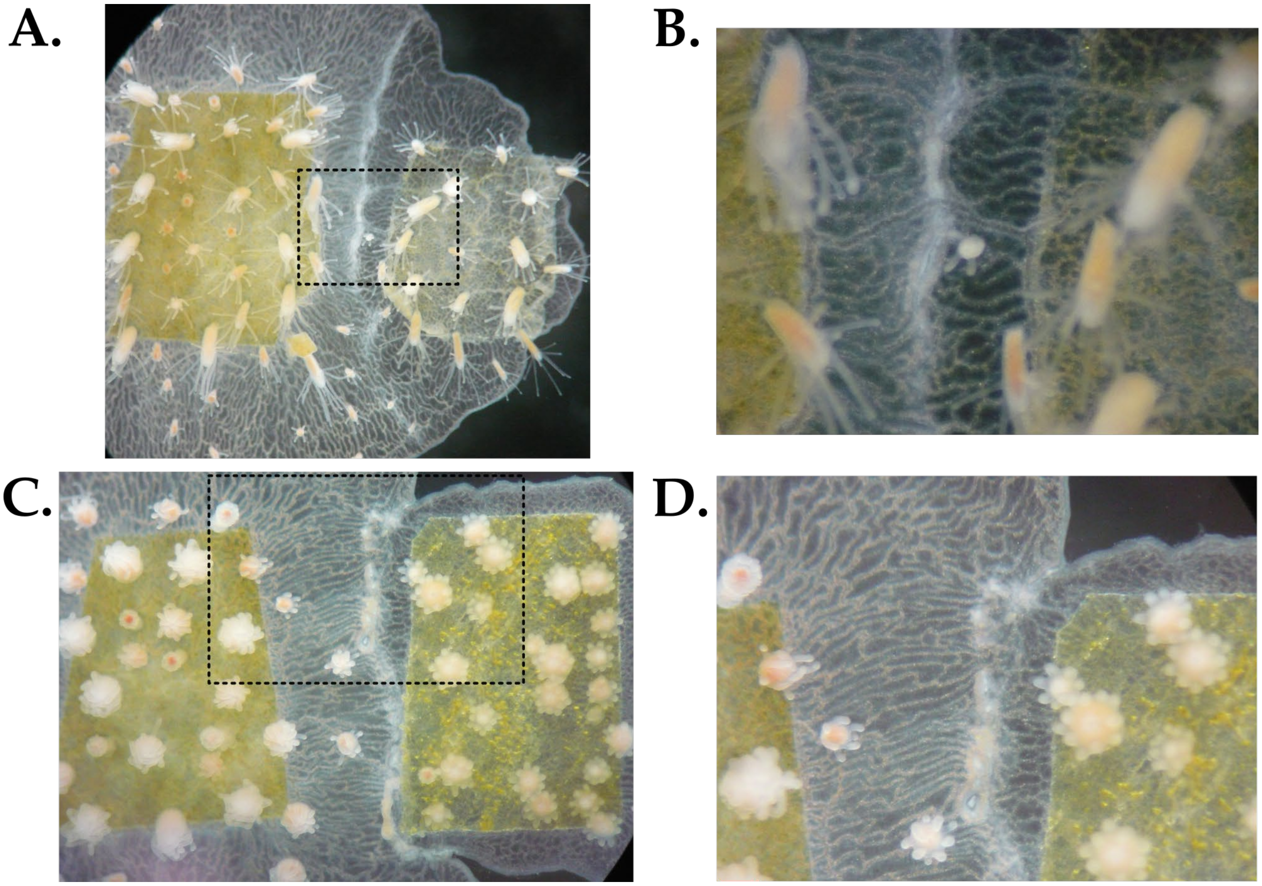


**Fig. S17 Transitory fusions in our backcross population.** Transitory fusion represented a heterogeneous collection of phenotypes in our backcross population. **A.** The contact region between the colonies shows fusion and rejection coexisting at the same time. **B.** Detail of the dotted rectangle in part A. A vessel of the gastrovascular system is observed that connects the colonies, and it is surrounded by rejection-like tissue. **C.** The contact region between colonies resembles a rejection phenotype. **D.** Detail of the dotted rectangle in part C. Rejection was observed predominantly in the contact region, but some regions still have a continuous gastrovascular system between the colonies

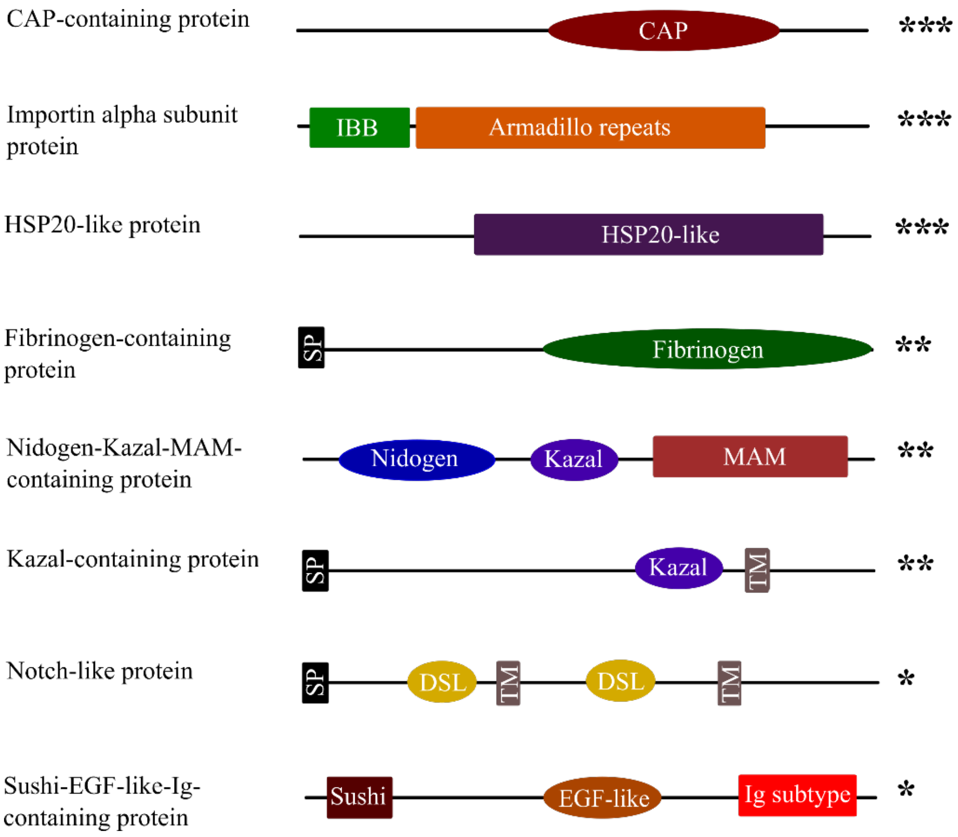


**Fig. S18 Domain architectures of the eight candidate allorecognition-proteins outside of the ARC.** These proteins showed variation potentially associated with allorecognition phenotypes among individuals BC3, BC15, HWB29, and HWB103. These proteins were identified outside ARC intervals. Domain abbreviations are CAP: Cysteine-rich secretory proteins, antigen 5, and pathogenesis-related 1; IBB: Importin beta binding; MAM: Meprin, A5 protein, and protein tyrosine phosphatase Mu; DSL: Delta/Serrate/Lag-2; Ig-subtype: Immunoglobulin-like domain; SP: Signal peptide; and TM: Transmembrane domain. Asterisks represent sequence variation levels (Nei's p-distance) between encoded proteins in individuals BC-3 and BC-15: *** (p-distance ≥ 0.20), ** (0.02 ≤ p-distance < 0.20), * (p-distance < 0.02)
