## Supplementary material for "Multiple *alr* genes exhibit allorecognition-associated variation in the colonial cnidarian *Hydractinia*": Table S4

**Table S4**. Predicted genes in the alr1, alr2, Interval 1, Interval 2, and Interval 3 of the Allorecognition Complex in individuals BC-3 and BC-15. Gene coordinates are given based on the genome of individual BC-15. The homology-based and functional annotations are described. Variable proteins between individuals are shown in bold and Nei’s p-distance is given

| INTERVAL | PROTEIN NUMBER | | PROTEIN NAME | LENGTH BC-3 (AA) | LENGTH BC-15 (AA) | GENE COORDINATES (DIRECTION) | HOMOLOGY-BASED ANNOTATION | | FUNCTIONAL ANNOTATION | | P-DISTANCE |
| --- | --- | --- | --- | --- | --- | --- | --- | --- | --- | --- | --- |
|  |  |  |  |  |  |  | **DESCRIPTION** | **E-VALUE** | **PREDICTED PROTEIN ARCHITECTURE** | **FUNCTIONAL DESCRIPTION** |  |
| ALR1 | | 1 | **alr1B** | **320** | **359** | **15147-21502**  **(-)** | **gi\|302457161\|gb\|ADL39772.1\| allorecognition 1 [Hydractinia symbiolongicarpus]** | **1.27E-68** | **Ig-like fold; TM** | **Recognition, membrane receptor; protein-protein binding; alr1-like** | **0.09** |
|  |  | 2 | **alr1C** | **332** | **356** | **83271-86898**  **(-)** | **gi\|302457161\|gb\|ADL39772.1\| allorecognition 1 [Hydractinia symbiolongicarpus]** | **7.97E-67** | **Ig-like fold; TM** | **Membrane receptor; recognition; protein-protein binding; alr1-like** | **0.18** |
|  |  | 3 | Uncharacterized protein | 1038 | 946 | 263066-269949  (+) | NO ANNOTATION | | NO PREDICTION | |  |
|  |  | 4 | **Ig-likeA** | **365** | **365** | **279228-286758**  **(+)** | **gi\|302457159\|gb\|ADL39771.1\| allorecognition 1 [Hydractinia symbiolongicarpus]** | **1.29E-24** | **TM X3** | **Membrane protein** | **0.01** |
|  |  | 5 | **alr1A** | **545** | **545** | **210195-221972**  **(+)** | **gi\|302457191\|gb\|ADL39787.1\| allorecognition 1 [Hydractinia symbiolongicarpus]** | **1.11E-55** | **SP; Ig-like fold; TM** | **Recognition, membrane receptor; protein-protein binding; alr1** | **0.13** |
|  |  | 6 | Fibrinogen-containing | 146 | 339 | 295730-301220  (+) | gi\|562886455\|ref\|XP_006170660.1\| PREDICTED: fibrinogen alpha chain [Tupaia chinensis] | 1.65E-14 | SP; Fibrinogen, alpha/beta/gamma chain, C-terminal globular domain | Receptor binding |  |
|  |  | 7 | Fibrinogen-containing | N/A | 163 | 314576-315999  (+) | gi\|260788738\|ref\|XP_002589406.1\| hypothetical protein BRAFLDRAFT_77850 [Branchiostoma floridae] | 2.33E-14 | Fibrinogen, alpha/beta/gamma chain, C-terminal globular domain | Receptor binding |  |
|  |  | 8 | **Ig-likeB** | **434** | **469** | **331198-336481**  **(+)** | **gi\|498944129\|ref\|XP_004521996.1\| PREDICTED: neogenin-like [Ceratitis capitata]** | **1.70E-06** | **Ig-like fold; LDL receptor class A repeat; Ig-like fold** | **Protein-protein binding** | **0.001** |
|  |  | 9 | TNF-like | 133 | 202 | 381102-384636  (-) | NO ANNOTATION | | TNF-like | Immune response; Receptor binding of tumoral necrosis factor |  |
|  |  | 10 | Uncharacterized secreted protein | 130 | 130 | 465720-466772  (+) | NO ANNOTATION | | SP | Secreted protein |  |
|  |  | 11 | Uncharacterized protein | 97 | 143 | 469277-470053  (+) | NO ANNOTATION | | NO PREDICTION | |  |
|  |  | 12 | EGF-like | 373 | 373 | 421620-424602  (+) | NO ANNOTATION | | SP; EGF-like | Protein-protein binding; cell adhesion; Secreted protein |  |
|  |  | 13 | Ig-likeC | N/A | 419 | 429335-433392  (-) | gi\|617419059\|ref\|XP_007557872.1\| PREDICTED: uncharacterized protein LOC103141974 isoform X2 [Poecilia formosa] | 1.09E-07 | Ig-like fold X2; LDL receptor class A repeat; FN3 X2 | Protein-protein binding |  |
|  |  | 14 | Uncharacterized protein | 125 | 85 | 510174-511648  (+) | NO ANNOTATION | | NO PREDICTION | |  |
|  |  | 15 | Uncharacterized secreted protein | 96 | 98 | 526032-526788  (+) | NO ANNOTATION | | SP | Secreted protein |  |
|  |  | 16 | Uncharacterized protein | N/A | 92 | 572446-572959  (+) | NO ANNOTATION | | NO PREDICTION | |  |
|  |  | 17 | **MACPF-containing** | **315** | **345** | **555212-561165**  **(-)** | **gi\|283137504\|gb\|ADB11402.1\| MAC/perforin- and kringle-domains-containing protein, partial [Pomatoceros lamarckii]** | **5.42E-43** | **MACPF** | **Immunity; lysis activity; lysis pore formation** | **0.01** |
|  |  | 18 | Uncharacterized secreted protein | N/A | 92 | 611247-611760  (+) | NO ANNOTATION | | SP | Secreted protein |  |
|  |  | 19 | Uncharacterized secreted protein | 110 | 68 | 649344-650210  (+) | NO ANNOTATION | | SP | Secreted protein |  |
|  |  | 20 | Uncharacterized protein | 123 | 585 | 667332-684741  (+) | NO ANNOTATION | | NO PREDICTION | |  |
|  |  | 21 | Uncharacterized membrane protein | 246 | 453 | 748389-756038  (+) | NO ANNOTATION | | TM | Membrane protein |  |
|  |  | 22 | **alr1D** | **283** | **445** | **701242-705191**  **(+)** | **gi\|302457159\|gb\|ADL39771.1\| allorecognition 1 [Hydractinia symbiolongicarpus]** | **1.85E-99** | **Ig-like fold; TM** | **Recognition; membrane receptor; protein-protein binding; alr1-like** | **0.001** |
|  |  | 23 | TNF-like | 126 | 160 | 759992-762057  (+) | NO ANNOTATION | | TNF-like | Immune response; Receptor binding of tumor necrosis factor; protein-protein binding |  |
|  |  | 24 | **Galactose-binding-like** | **264** | **240** | **767425-771246**  **(+)** | **gi\|551548771\|ref\|XP_005762028.1\| regulator of telomere elongation helicase [Emiliania huxleyi CCMP1516]** | **3.53E-08** | **SP; ShKT; Galactose binding-like** | **Cell adhesion** | **0.04** |
|  |  | 25 | Hydrolase-fold-containing | 322 | 322 | 795912-799197  (-) | gi\|449683348\|ref\|XP_004210331.1\| PREDICTED: uncharacterized protein LOC101241114 [Hydra vulgaris] | 2.55E-107 | SP; alpha/beta hydrolase fold | Lipid metabolism |  |
|  |  | 26 | MFS transporter-like | 628 | 628 | 804352-809728  (-) | gi\|449667661\|ref\|XP_002154002.2\| PREDICTED: monocarboxylate transporter 10-like [Hydra vulgaris] | 0.00 | MFS_general substrate transporter X2 | Transmembrane transport; integral membrane component |  |
|  |  | 27 | Uncharacterized membrane protein | 140 | 171 | 869896-876162  (-) | NO ANNOTATION | | SP; TM | Membrane protein |  |
|  |  | 28 | **MFS transporter-like** | **278** | **281** | **826269-830975**  **(-)** | **gi\|449686970\|ref\|XP_002162044.2\| PREDICTED: monocarboxylate transporter 10-like [Hydra vulgaris]** | **3.20E-111** | **MFS_general substrate transporter X2** | **Transmembrane transport; integral membrane component** | **0.001** |
|  |  | 29 | **Glutamate receptor-like** | **519** | **709** | **887674-893120**  **(+)** | **gi\|585651030\|ref\|XP_006814676.1\| PREDICTED: glutamate receptor ionotropic, NMDA 2A-like [Saccoglossus kowalevskii]** | **2.10E-36** | **Glutamate receptor L-glutamate/glycine binding; TM X2** | **Transporter activity; ionotropic receptor activity; Membrane protein** | **0.001** |
|  |  | 30 | Uncharacterized secreted protein | 170 | 170 | 1011435-1013370  (+) | NO ANNOTATION | | SP | Secreted protein |  |
|  |  | 31 | TNF-like | 253 | 120 | 1031640- 1032559  (+) | NO ANNOTATION | | TM; TNF-like | Immune response; Receptor binding of tumor necrosis factor; Membrane protein |  |
|  |  | 32 | Ig-likeD | 242 | 261 | 1037711-1047563  (+) | gi\|512842688\|ref\|XP_002931559.2\| PREDICTED: thrombospondin-2 [Xenopus (Silurana) tropicalis] | 4.36E-07 | Ig-like fold; TSP_1; TM | Protein binding; cell adhesion; Membrane protein |  |
|  |  | 33 | GPCR-like | 95 | 95 | 1051000-1051512  (-) | NO ANNOTATION | | GPCR, family 2, secretin-like; TM X3 | G-coupling receptor activity; Membrane protein |  |
|  |  | 34 | GPCR-like | 96 | 78 | 1052484-1052817  (-) | NO ANNOTATION | | GPCR, family 2, secretin-like; TM X3 | G-coupling receptor activity; Membrane protein |  |
|  |  | 35 | VWA typeA-containing | 263 | 185 | 990164-993615  (-) | gi\|449663000\|ref\|XP_004205667.1\| PREDICTED: collagen alpha-1(XII) chain-like [Hydra vulgaris] | 4.00E-20 | SP; VWA_type A | Ligand interaction; multi-protein complexes; cell adhesion, cell migration |  |
|  |  | 36 | LDL repeat-containing | 338 | 499 | 1054271-1059724  (-) | NO ANNOTATION | | LDL receptor class A repeat; TM | Protein-protein binding |  |
|  |  | 37 | Uncharacterized protein | 230 | 240 | 1166927-1170330  (+) | sp\|Q5PQJ7\|TBCEL_RAT Tubulin-specific chaperone cofactor E-like protein | 3.70E-63 | NO PREDICTION | |  |
|  |  | 38 | **Ank repeat-containing** | **380** | **380** | **1206304-1210477**  **(+)** | **gi\|528504032\|ref\|XP_694014.5\| PREDICTED: B-cell lymphoma 3 protein homolog [Danio rerio]** | **4.55E-23** | **Ank_repeat containing** | **Protein-protein binding** | **0.001** |
|  |  | 39 | **Uncharacterized protein** | **443** | **443** | **1110102-1115469**  **(-)** | **gi\|156359434\|ref\|XP_001624774.1\| predicted protein [Nematostella vectensis]** | **2.69E-72** | **NO PREDICTION** | | **0.001** |
|  |  | 40 | Zinc finger-containing | 1975 | 1975 | 1083750-1108372  (-) | gi\|585683356\|ref\|XP_006812408.1\| PREDICTED: NFX1-type zinc finger-containing protein 1-like [Saccoglossus kowalevskii] | 0.00 | P-loop containing nucleoside triphosphate hydrolase; Zinc finger, NF-X1-type | DNA-binding transcription factor activity |  |
|  |  | 41 | **Integrin** | **790** | **790** | **1118049-1130726**  **(-)** | **gi\|10998792\|gb\|AAG25994.1\|AF308652_1 integrin beta chain [Podocoryna carnea]** | **0.00** | **Integrin beta subunit; TM** | **Receptor activity; Membrane protein** | **0.001** |
|  |  | 42 | Polyketide synthase | 398 | 398 | 1133395-1140665  (-) | gi\|449692707\|ref\|XP_004213141.1\| PREDICTED: synaptic vesicle membrane protein VAT-1 homolog, partial [Hydra vulgaris] | 1.49E-119 | Polyketide synthase, enoylreductase | Redox activity; oxidoreductase activity, transferase |  |
|  |  | 43 | **DnaJ-like** | **511** | **496** | **1158245-1166145**  **(-)** | **gi\|449664494\|ref\|XP_002163923.2\| PREDICTED: dnaJ homolog subfamily C member 3-like [Hydra vulgaris]** | **0.00** | **SP; Tetratricopeptide-like; DnaJ** | **Protein-protein binding** | **0.05** |
| ALR2 | | 44 | **Ank repeat-containing** | **1720** | **1720** | **64111-75333**  **(+)** | **gi\|164609120\|gb\|ABY62780.1\| ankyrin repeat protein 17-like protein [Hydractinia symbiolongicarpus]** | **0.00** | **Ankyrin repeat-containing; K Homology; Sterile alpha motif/pointed** | **RNA binding; protein-protein binding** | **0.001** |
|  |  | 45 | **Zinc finger-containing** | **405** | **301** | **27190-41430**  **(+)** | **sp\|A6NGD5\|ZSA5C_HUMAN Zinc finger and SCAN domain-containing protein** | **5.95E-07** | **Chromo domain-like; Zinc finger C2H2-type/integrase DNA-binding** | **DNA binding; metal ion binding** | **0.06** |
|  |  | 46 | **Uncharacterized protein** | **204** | **153** | **24078-25480**  **(-)** | **NO ANNOTATION** | | **NO PREDICTION** |  | **0.13** |
|  |  | 47 | **ATP synthase** | **164** | **248** | **116898-122656**  **(+)** | **gi\|313105484\|gb\|ADR32100.1\| ATP synthase [Hydractinia symbiolongicarpus]** | **1.54E-143** | **Orthogonal Bundle domain in ATP12** | **ATP complex synthase; ion transport** | **0.15** |
|  |  | 48 | **Actin-related** | **571** | **564** | **86474-92069**  **(-)** | **gi\|164609121\|gb\|ABY62781.1\| actin related protein 8-like protein [Hydractinia symbiolongicarpus]** | **0.00** | **Actin-related protein family** | **Chromatin remodeling** | **0.12** |
|  |  | 49 | **alr2C** | **348** | **178** | **181018-192598**  **(+)** | **gi\|225423264\|gb\|ACN91138.1\| allorecognition 2 [Hydractinia symbiolongicarpus]** | **0.00** | **SP; Ig-like fold X3; TM** | **Recognition; protein-protein binding** | **0.08** |
|  |  | 50 | Uncharacterized protein | 196 | 218 | 287301-288931  (+) | NO ANNOTATION | | NO PREDICTION | |  |
|  |  | 51 | **alr2A** | **680** | **680** | **206551-222431**  **(+)** | **gi\|225423264\|gb\|ACN91138.1\| allorecognition 2 [Hydractinia symbiolongicarpus]** | **0.00** | **SP; Ig-like fold X3; TM** | **Recognition; protein-protein binding** | **0.03** |
|  |  | 52 | Fucosyl-transferase | 512 | 512 | 233718-236264  (-) | gi\|313105490\|gb\|ADR32105.1\| fucosyl_transferase [Hydractinia symbiolongicarpus] | 0.00 | TM; Glycosyltransferase family 23 (GT23) | Cell adhesion |  |
|  |  | 53 | Uncharacterized protein | 227 | 138 | 254849-257136  (-) | gi\|164609125\|gb\|ABY62785.1\| unknown protein [Hydractinia symbiolongicarpus] | 3.17E-13 | NO PREDICTION | |  |
|  |  | 54 | Uncharacterized protein | 257 | 482 | 245447-250207  (-) | gi\|164609125\|gb\|ABY62785.1\| unknown protein [Hydractinia symbiolongicarpus] | 0.00 | NO PREDICTION | |  |
|  |  | 55 | **alr2B** | **733** | **516** | **157099-179062**  **(+)** | **gi\|225423264\|gb\|ACN91138.1\| allorecognition 2 [Hydractinia symbiolongicarpus]** | **0.00** | **SP; Ig-like fold X3; TM** | **Recognition; protein-protein binding** | **0.17** |
|  |  | 56 | Uncharacterized membrane protein | 231 | 279 | 336679-340229  (+) | NO ANNOTATION | | TM | Membrane protein |  |
|  |  | 57 | Sodium channel-containing | 280 | 525 | 383246-392500  (+) | gi\|449671675\|ref\|XP_004207541.1\| PREDICTED: uncharacterized protein LOC101235583 [Hydra vulgaris] | 3.56E-100 | Epithelial sodium channel | Sodium channel activity; sodium ions transport |  |
|  |  | 58 | **Activating signal cointegrator** | **354** | **354** | **406650-415662**  **(+)** | **gi\|449668128\|ref\|XP_004206716.1\| PREDICTED: activating signal cointegrator 1 complex subunit 1-like, partial [Hydra vulgaris]** | **8.93E-93** | **K Homology; RNA ligase/cyclic nucleotide phosphodiesterase** | **RNA binding; catalytic activity** | **0.01** |
|  |  | 59 | Zinc finger-containing | 570 | 856 | 394570-404890  (+) | sp\|P10072\|HKR1_HUMAN Krueppel-related zinc finger protein 1 OS=Homo sapiens GN=HKR1 PE=2 SV=4 | 4.09E-14 | Chromo domain-like; Zinc finger C2H2-type/integrase DNA-binding X2 | DNA binding; metal ion binding |  |
|  |  | 60 | **Ig-likeE** | **466** | **368** | **474857-481398**  **(+)** | **gi\|302457155\|gb\|ADL39769.1\| allorecognition 1 [Hydractinia symbiolongicarpus]** | **3.74E-07** | **SP** | **Secreted protein** | **0.13** |
|  |  | 61 | DNA polymerase subunit | 308 | 531 | 515279-524977  (+) | gi\|532074904\|ref\|XP_005323251.1\| PREDICTED: DNA polymerase delta subunit 3 [Ictidomys tridecemlineatus] | 1.46E-30 | DNA polymerase subunit Cdc27 | DNA replication |  |
|  |  | 62 | Uncharacterized protein | 778 | 778 | 458216-462496  (+) | gi\|156403588\|ref\|XP_001639990.1\| predicted protein [Nematostella vectensis] | 6.66E-51 | NO PREDICTION | |  |
|  |  | 63 | Antistasin-like | 213 | 223 | 499687-503301  (-) | gi\|524873557\|ref\|XP_005093523.1\| PREDICTED: antistasin-like [Aplysia californica] | 2.81E-12 | SP; Insulin-like growth factor-binding protein, IGFBP; Antistasin-like domainX2 | Cell growth regulation; growth factor binding; inhibitor of serine-type endopeptidase |  |
|  |  | 64 | **Peptidase M13-containing** | **678** | **714** | **485339-491975**  **(-)** | **gi\|564245265\|ref\|XP_006261756.1\| PREDICTED: endothelin-converting enzyme 1 [Alligator mississippiensis]** | **7.06E-124** | **TM;Peptidase M13** | **Membrane protein; proteolysis; metallopeptidase activity** | **0.01** |
|  |  | 65 | **MFS transporter-like** | **707** | **526** | **431388-453495**  **(-)** | **gi\|449663924\|ref\|XP_002168497.2\| PREDICTED: major facilitator superfamily domain-containing protein 1-like [Hydra vulgaris]** | **0.00** | **MFS_general substrate transporter X2** | **Transmembrane transport; integral membrane component** | **0.13** |
|  |  | 66 | **ATPase subunit beta-like** | **299** | **330** | **417924-423201**  **(-)** | **gi\|449684222\|ref\|XP_002167147.2\| PREDICTED: sodium/potassium-transporting ATPase subunit beta-1-like [Hydra vulgaris]** | **9.92E-68** | **Sodium/potassium-transporting ATPase subunit beta** | **Sodium and potassium ions transport** | **0.01** |
|  |  | 67 | MFS transporter-like | 335 | 366 | 496551-498747  (-) | gi\|254728979\|gb\|ACT79639.1\| transposase [Acropora millepora] | 1.23E-32 | MFS_general substrate transporter X2 | Transmembrane transport; integral membrane component |  |
| INTERVAL 1 | | 68 | Zinc finger-containing | 678 | 699 | 26817-35686  (-) | gi\|449680951\|ref\|XP_002158120.2\| PREDICTED: U2 small nuclear ribonucleoprotein auxiliary factor 35 kDa subunit-related protein 2-like [Hydra vulgaris] | 7.82E-42 | Zinc finger, CCCH-type; Nucleotide-binding, alpha-beta plait | DNA, RNA and metal ion binding |  |
|  |  | 69 | **Mediator complex** | **1355** | **1404** | **3460- 21540**  **(-)** | **gi\|449671974\|ref\|XP_002162802.2\| PREDICTED: mediator of RNA polymerase II transcription subunit 14-like [Hydra vulgaris]** | **0.00** | **Mediator complex, subunit Med14** | **RNA polymerase II transcription cofactor activity** | **0.01** |
|  |  | 70 | **Zinc finger-containing** | **452** | **459** | **36233-52106**  **(-)** | **gi\|449680951\|ref\|XP_002158120.2\| PREDICTED: U2 small nuclear ribonucleoprotein auxiliary factor 35 kDa subunit-related protein 2-like [Hydra vulgaris]** | **8.32E-81** | **Zinc finger, CCCH-type; Nucleotide-binding, alpha-beta plait** | **DNA, RNA and metal ion binding** | **0.04** |
|  |  | 71 | Zinc finger-containing | 519 | 519 | 66999-76533  (-) | gi\|449680951\|ref\|XP_002158120.2\| PREDICTED: U2 small nuclear ribonucleoprotein auxiliary factor 35 kDa subunit-related protein 2-like [Hydra vulgaris] | 2.18E-151 | Zinc finger, CCCH-type; Nucleotide-binding, alpha-beta plait | DNA, RNA and metal ion binding |  |
|  |  | 72 | Homeodomain-like | 174 | 174 | 250-1163  (-) | gi\|222478322\|gb\|ACM62738.1\| NK2b homeodomain transcription factor protein [Clytia hemisphaerica] | 2.31E-21 | Homeodomain-like | DNA binding |  |
|  |  | 73 | Arf GTPase activating | 527 | 584 | 97808-107584  (+) | sp\|Q2TA45\|AGFG1_BOVIN Arf-GAP domain and FG repeat-containing protein 1 OS=Bos taurus GN=AGFG1 PE=2 SV=1 | 1.65E-65 | Arf GTPase activating protein | GTPase activation; zinc binding |  |
|  |  | 74 | Uncharacterized secreted protein | 319 | 319 | 126648-127765  (+) | gi\|49146397\|ref\|YP_025505.1\| putative phage-related minor tail protein [Caedibacter taeniospiralis] | 9.58E-22 | SP | Secreted protein |  |
|  |  | 75 | Uncharacterized protein | 96 | 87 | 180800-184570  (-) | gi\|449672060\|ref\|XP_004207623.1\| PREDICTED: regulator complex protein LAMTOR4 homolog [Hydra vulgaris] | 9.14E-29 | NO PREDICTION | |  |
|  |  | 76 | **MFS transporter-like** | **445** | **445** | **151223-152560**  **(-)** | **gi\|221124576\|ref\|XP_002168703.1\| PREDICTED: equilibrative nucleoside transporter 1-like [Hydra vulgaris]** | **3.22E-165** | **MFS_general substrate transporter X2** | **Transmembrane transport; integral membrane component** | **0.01** |
|  |  | 77 | **AMP deaminase** | **812** | **866** | **163604-175019**  **(-)** | **gi\|585708691\|ref\|XP_006899389.1\| PREDICTED: AMP deaminase 2 isoform X3 [Elephantulus edwardii]** | **0.00** | **Adenosine/AMP deaminase domain** | **Monophosphate purine biosynthesis; deaminase activity** | **0.01** |
|  |  | 78 | **HyFMR1** | **784** | **753** | **131739-140437**  **(-)** | **gi\|51889276\|emb\|CAH25437.1\| HyFMR1 protein [Hydractinia echinata]** | **0.00** | **Agenet-like domain X2; K Homology_type 1** | **RNA binding** | **0.02** |
|  |  | 79 | Uncharacterized protein | 85 | 75 | 286022-286761  (+) | NO ANNOTATION | | NO PREDICTION | |  |
|  |  | 80 | **HAD-like hydrolase** | **242** | **242** | **255276-259499**  **(+)** | **gi\|221123627\|ref\|XP_002156375.1\| PREDICTED: pseudouridine-5'-monophosphatase-like [Hydra vulgaris]** | **2.45E-102** | **HAD-like hydrolase** | **Hydrolase activity** | **0.01** |
|  |  | 81 | **Zinc finger-containing** | **454** | **436** | **241729-247101**  **(+)** | **gi\|449668312\|ref\|XP_002159766.2\| PREDICTED: nuclear fragile X mental retardation-interacting protein 1-like [Hydra vulgaris]** | **4.64E-50** | **Zinc finger, C2H2-like** | **DNA binding; metal ion binding** | **0.02** |
|  |  | 82 | **Ig-likeF** | **417** | **443** | **211177-217594**  **(+)** | **gi\|302171734\|gb\|ADK97768.1\| allorecognition 2-like protein [Hydractinia polyclina]** | **1.04E-10** | **SP; Ig-like fold X2; TM** | **Recognition; protein-protein binding; alr2-like** | **0.03** |
|  |  | 83 | Uncharacterized protein | 106 | 147 | 278659-281028  (+) | gi\|340383707\|ref\|XP_003390358.1\| PREDICTED: hypothetical protein LOC100638403 [Amphimedon queenslandica] | 2.71E-19 | NO PREDICTION | |  |
|  |  | 84 | **VWA typeA-containing** | **321** | **321** | **199861-201761**  **(-)** | **sp\|Q923P0\|COKA1_MOUSE Collagen alpha-1(XX) chain OS=Mus musculus GN=Col20a1 PE=2 SV=3** | **1.20E-14** | **VWA_type A** | **Ligand interaction; multi-protein complexes; cell adhesion** | **0.01** |
|  |  | 85 | Zinc finger-containing | 184 | 619 | 217907-228830  (-) | gi\|449672333\|ref\|XP_004207690.1\| PREDICTED: vacuolar segregation protein PEP7-like, partial [Hydra vulgaris] | 3.25E-174 | SP; Zinc finger, RING/FYVE/PHD-type X2 | Metal ion binding |  |
|  |  | 86 | **Proteinase inhibitor** | **902** | **1361** | **185705-198886**  **(-)** | **sp\|P81439\|EQST_ACTEQ Equistatin OS=Actinia equina PE=1 SV=1** | **2.97E-18** | **Proteinase inhibitor I2, Kunitz; Thyroglobulin type-1** | **Inhibitory activity of serine-type endopeptidase** | **0.07** |
|  |  |  |  | **638** |  |  |  |  |  |  |  |
|  |  | 87 | PapD-like | 215 | 215 | 252170-255908  (-) | gi\|449663382\|ref\|XP_002155118.2\| PREDICTED: motile sperm domain-containing protein 1-like [Hydra vulgaris] | 1.47E-84 | PapD-like; TM X2 | Structural activity; periplasmic adhesion chaperones |  |
|  |  | 88 | Neurotransmitter-gated ion channel | 315 | 140 | 273358-274379  (-) | gi\|557777992\|ref\|XP_005188629.1\| PREDICTED: CHRNA7-FAM7A fusion protein-like, partial [Musca domestica] | 7.07E-24 | Neurotransmitter-gated ion-channel ligand-binding | Ligand-mediated extracellular ion channel activity |  |
|  |  | 89 | Neurotransmitter-gated ion channel | 213 | 114 | 260150-269695  (-) | gi\|449663525\|ref\|XP_002156390.2\| PREDICTED: transcription initiation factor TFIID subunit 9B-like [Hydra vulgaris] | 3.28E-28 | Histone-fold | Protein heterodimerization activity |  |
|  |  | 90 | Histone-fold-containing | 135 | 137 | 274615-281467  (-) | sp\|P02717\|ACHD_CHICK Acetylcholine receptor subunit delta OS=Gallus gallus GN=CHNRD PE=3 SV=1 | 1.92E-08 | Neurotransmitter-gated ion-channel ligand-binding | Ligand-mediated extracellular ion channel activity |  |
|  |  | 91 | Peptidase M28-containing | 737 | 717 | 321850-330438  (+) | sp\|P70627\|FOLH1_RAT Glutamate carboxypeptidase 2 OS=Rattus norvegicus GN=Folh1 PE=2 SV=1 | 1.70E-146 | TM; Peptidase M28; Transferrin receptor-like, dimerisation | Catalytic activity/peptidase; recognition; Protein binding |  |
|  |  | 92 | Ank repeat-containing | 891 | 854 | 523822-528045  (+) | gi\|602646222\|ref\|XP_007429544.1\| PREDICTED: NF-kappa-B inhibitor epsilon [Python bivittatus] | 1.00E-30 | Ank_repeat containing | Protein-protein binding |  |
|  |  | 93 | Porphobilinogen deaminase | 386 | 386 | 469957-474578  (+) | sp\|P08397\|HEM3_HUMAN Porphobilinogen deaminase OS=Homo sapiens GN=HMBS PE=1 SV=2 | 5.60E-111 | Porphobilinogen deaminase | Hydroxymethylbilane synthase activity |  |
|  |  | 94 | BRCT-containing | 879 | 872 | 551600-562338  (+) | gi\|443727325\|gb\|ELU14128.1\| hypothetical protein CAPTEDRAFT_176435 [Capitella teleta] | 0.00 | BRCT domain X2; Dbl homology (DH) | GTPase activation; Guanyl nucleotide exchange factor activity |  |
|  |  | 95 | RCC1 / BTB-containing | 381 | 392 | 385568-397319  (+) | gi\|221113828\|ref\|XP_002154529.1\| PREDICTED: RCC1 and BTB domain-containing protein 1-like, partial [Hydra vulgaris] | 0.00 | RCC1/beta-lactamase-inhibitor protein II; BTB/POZ fold | Protein-protein interaction |  |
|  |  | 96 | **Protein kinase-like** | **1092** | **939** | **575566-586421**  **(+)** | **gi\|221108130\|ref\|XP_002169972.1\| PREDICTED: putative serine protease K12H4.7-like [Hydra vulgaris]** | **9.57E-168** | **SP; Alpha/Beta hydrolase fold; Protein kinase-like** | **Serin-peptidase activity; transferase activity; ATP binding** | **0.04** |
|  |  | 97 | TNF-like | 268 | 268 | 482489-485954  (-) | NO ANNOTATION | | TM; TNF-like | Immune response; Receptor binding of tumor necrosis factor |  |
|  |  | 98 | Uncharacterized protein | 833 | 833 | 533298-536217  (-) | NO ANNOTATION | | NO PREDICTION | |  |
|  |  | 99 | Protein kinase-like | 384 | 388 | 437896-447437  (-) | gi\|449672319\|ref\|XP_002166025.2\| PREDICTED: serine/threonine-protein kinase 25-like, partial [Hydra vulgaris] | 6.45E-169 | Protein kinase-like | Kinase activity; phosphotransferase activity |  |
|  |  | 100 | Cullin repeat-like-containing | 154 | 126 | 376671-380599  (-) | gi\|584010127\|ref\|XP_006799999.1\| PREDICTED: cullin-3-like [Neolamprologus brichardi] | 3.07E-21 | Cullin repeat-like-containing | Protein catabolism, ubiquitin-dependent |  |
| Interval 2 | | 101 | **Kazal / EF-hand-containing** | **272** | **290** | **762- 5316**  **(+)** | **gi\|449679846\|ref\|XP_002170609.2\| PREDICTED: follistatin-related protein 1-like, partial [Hydra vulgaris]** | **2.81E-62** | **SP; Kazal; EF-hand domain X2** | **Protein-protein binding; calcium binding** | **0.01** |
|  |  | 102 | **Synthase beta subunit-like** | **613** | **355** | **9206-20212**  **(+)** | **gi\|449679284\|ref\|XP_002163391.2\| PREDICTED: cystathionine beta-synthase-like [Hydra vulgaris]** | **3.76E-163** | **Tryptophan synthase beta subunit-like PLP-dependent enzyme** | **Biosynthesis process of cysteine from serine** | **0.01** |
|  |  | 103 | Uncharacterized membrane protein | 81 | 83 | 44442-55853  (+) | gi\|221113401\|ref\|XP_002170443.1\| PREDICTED: negative elongation factor A-like [Hydra vulgaris] | 1.88E-116 | TM X2 | Membrane protein |  |
|  |  | 104 | Uncharacterized protein | 159 | 188 | 61385-62907  (+) | NO ANNOTATION | | NO PREDICTION | |  |
|  |  | 105 | Peptidase M13-containing | 735 | 752 | 64695-79468  (-) | gi\|591388068\|ref\|XP_007068447.1\| PREDICTED: endothelin-converting enzyme 2-like [Chelonia mydas] | 2.18E-155 | TM; Peptidase M13 | Proteolysis; metallopeptidase activity; Secreted protein |  |
|  |  | 106 | Metallopeptidase-like | 319 | 319 | 29185-32026  (-) | gi\|449691922\|ref\|XP_002163948.2\| PREDICTED: 72 kDa type IV collagenase-like [Hydra vulgaris] | 3.87E-69 | SP; Metallopeptidase, catalytic domain; ShKT | Proteolysis; metallopeptidase activity; Secreted protein |  |
|  |  | 107 | Uncharacterized membrane protein | 265 | 222 | 20755-26427  (-) | NO ANNOTATION | | TM | Membrane protein |  |
|  |  | 108 | Pyruvate formate lyase-like | 1285 | 1285 | 198968-206560  (+) | gi\|260803037\|ref\|XP_002596398.1\| hypothetical protein BRAFLDRAFT_76215 [Branchiostoma floridae] | 0.00 | Pyruvate formate lyase | Redox activity; catalytic activity, peroxidase activity |  |
|  |  | 109 | TNF receptor-associated factor | 288 | 325 | 149578-151137  (+) | gi\|548507603\|ref\|XP_005693417.1\| PREDICTED: TNF receptor-associated factor 4, partial [Capra hircus] | 5.89E-14 | TNF receptor-associated factor 6 | Signal transduction activity; protein-protein binding; transferase activity |  |
|  |  | 110 | Cation/H+ exchanger-containing | 828 | 545 | 188126-194273  (+) | gi\|449662021\|ref\|XP_002158626.2\| PREDICTED: transmembrane and coiled-coil domain-containing protein 3-like [Hydra vulgaris] | 1.80E-108 | Cation/H+ exchanger | Transmembrane transport; antiport activity, integral membrane component |  |
|  |  | 111 | Peptidase M13-containing | 110 | 110 | 160670-162125  (-) | gi\|25245872\|gb\|AAN73018.1\| endothelin-converting enzyme [Locusta migratoria] | 3.28E-27 | Peptidase M13 | Proteolysis; metallopeptidase activity; Secreted protein |  |
|  |  | 112 | EGF-like | 328 | 355 | 113781-116602  (-) | gi\|156406546\|ref\|XP_001641106.1\| predicted protein [Nematostella vectensis] | 4.12E-06 | EGF-like | Protein-protein binding |  |
|  |  | 113 | Uncharacterized protein | 339 | 302 | 176492-179895  (-) | gi\|449675774\|ref\|XP_004208486.1\| PREDICTED: uncharacterized protein LOC101235187 [Hydra vulgaris] | 3.53E-20 | NO PREDICTION | |  |
|  |  | 114 | Uncharacterized protein | 597 | 342 | 139930-143983  (-) | NO ANNOTATION | | NO PREDICTION | |  |
|  |  | 115 | Peptidase M13-containing | 211 | 532 | 162533-166003  (-) | gi\|327279269\|ref\|XP_003224379.1\| PREDICTED: endothelin-converting enzyme 2-like [Anolis carolinensis] | 5.25E-42 | Peptidase M13 | Proteolysis; metallopeptidase activity |  |
| Interval 3 | | 116 | RNA polymerase | 1949 | 1967 | 13291-31698  (+) | gi\|444722932\|gb\|ELW63604.1\| DNA-directed RNA polymerase II subunit RPB1 [Tupaia chinensis] | 0.00 | RNA polymerase domains | RNA polymerase II promoter-based transcription; RNA polymerase activity; DNA binding |  |
|  |  | 117 | Negative elongation factor | 603 | 591 | 60547-66639  (-) | sp\|Q86NP2\|NELFA_DROME Negative elongation factor A OS=Drosophila melanogaster GN=Nelf-A PE=1 SV=2 | 2.56E-33 | Negative elongation factor A | Negative regulation of transcription elongation |  |
|  |  | 118 | Uncharacterized membrane protein | 234 | 234 | 105314-108676  (+) | gi\|449678480\|ref\|XP_002159311.2\| PREDICTED: uncharacterized protein LOC100202149 [Hydra vulgaris] | 6.53E-73 | TM X2 | Membrane protein |  |
|  |  | 119 | E6-AP HECT-like | 331 | 331 | 186273-187906  (-) | NO ANNOTATION | | Homologous to the E6-AP Carboxyl Terminus HECT; TM | Transferase activity, ubiquitin |  |
