## Supplementary material for "Multiple *alr* genes exhibit allorecognition-associated variation in the colonial cnidarian *Hydractinia*": Table S5

**Table S5**. Predicted proteins in the reference alr1 interval of the Allorecognition Complex assembled from BAC clones listed in Table S1. The homology-based and functional annotations are described. The predicted protein in bold is not present in the Alr1 Interval of individual BC-15

| PROTEIN NUMBER | PROTEIN NAME | LENGTH  (AA) | GENE COORDINATES (DIRECTION) | HOMOLOGY-BASED ANNOTATION | | FUNCTIONAL ANNOTATION | |
| --- | --- | --- | --- | --- | --- | --- | --- |
|  |  |  |  | **DESCRIPTION** | **EVALUE** | **PREDICTED PROTEIN ARCHITECTURE** | **FUNCTIONAL DESCRIPTION** |
| 1 | alr1B | 257 | 14520-17187 (-) | gi\|302457161\|gb\|ADL39772.1\| allorecognition 1 [Hydractinia symbiolongicarpus] | 1.27E-68 | Ig-like fold X2; TM | Recognition, membrane receptor; protein-protein binding; alr1-like |
| 2 | alr1C | 324 | 83271-90100 (-) | gi\|302457161\|gb\|ADL39772.1\| allorecognition 1 [Hydractinia symbiolongicarpus] | 7.97E-67 | Ig-like fold; TM | Membrane receptor; recognition; protein-protein binding; alr1-like |
| 3 | **Uncharacterized membrane protein** | **148** | **149202-159681**  **(-)** | **ADL39773.1 allorecognition 1 [Hydractinia symbiolongicarpus]** | **5.0E-63** | **Cytoplasmic domain; TM; non-cytoplasmic domain** | **Membrane protein** |
| 4 | Uncharacterized protein | 1126 | 263066-272025  (+) | NO ANNOTATION | | NO PREDICTION | |
| 5 | Ig-likeA | 365 | 279228-286758  (+) | gi\|302457159\|gb\|ADL39771.1\| allorecognition 1 [Hydractinia symbiolongicarpus] | 1.29E-24 | Ig-like fold; TM X3 | Membrane protein |
| 6 | alr1A | 742 | 208931-221972  (+) | gi\|302457191\|gb\|ADL39787.1\| allorecognition 1 [Hydractinia symbiolongicarpus] | 1.11E-55 | SP; Ig-like fold X2; TM | Recognition, membrane receptor; protein-protein binding; alr1 |
| 7 | Fibrinogen-containing | 339 | 295730-301220  (+) | gi\|562886455\|ref\|XP_006170660.1\| PREDICTED: fibrinogen alpha chain [Tupaia chinensis] | 1.65E-14 | SP; Fibrinogen, alpha/beta/gamma chain, C-terminal globular domain | Receptor binding |
| 8 | Fibrinogen-containing | 339 | 310410-315999  (+) | gi\|260788738\|ref\|XP_002589406.1\| hypothetical protein BRAFLDRAFT_77850 [Branchiostoma floridae] | 2.33E-14 | Fibrinogen, alpha/beta/gamma chain, C-terminal globular domain | Receptor binding |
| 9 | Ig-likeB | 497 | 331198-336481  (+) | gi\|498944129\|ref\|XP_004521996.1\| PREDICTED: neogenin-like [Ceratitis capitata] | 1.70E-06 | Ig-like fold; LDL receptor class A repeat; Ig-like fold | Protein-protein binding |
| 10 | TNF-like | 303 | 381102-384636  (-) | NO ANNOTATION | | TNF-like | Immune response; Receptor binding of tumoral necrosis factor |
| 11 | Uncharacterized secreted protein | 156 | 463859-466772  (+) | NO ANNOTATION | | SP | Secreted protein |
| 12 | Uncharacterized protein | 143 | 469277-470053  (+) | NO ANNOTATION | | NO PREDICTION | |
| 13 | EGF-like | 369 | 421620-424602  (+) | NO ANNOTATION | | SP; EGF-like | Protein-protein binding; cell adhesion; Secreted protein |
| 14 | Ig-likeC | 419 | 429335-433392  (-) | gi\|617419059\|ref\|XP_007557872.1\| PREDICTED: uncharacterized protein LOC103141974 isoform X2 [Poecilia formosa] | 1.09E-07 | Ig-like fold X2; LDL receptor class A repeat; FN3 X2 | Protein-protein binding |
| 15 | Uncharacterized protein | 102 | 510174-511648  (+) | NO ANNOTATION | | NO PREDICTION | |
| 16 | Uncharacterized secreted protein | 221 | 518250-526796  (+) | NO ANNOTATION | | SP | Secreted protein |
| 17 | Uncharacterized protein | 89 | 572283-572915  (+) | NO ANNOTATION | | NO PREDICTION | |
| 18 | MACPF-containing | 345 | 555212-561165  (-) | gi\|283137504\|gb\|ADB11402.1\| MAC/perforin- and kringle-domains-containing protein, partial [Pomatoceros lamarckii] | 5.42E-43 | MACPF | Immunity; lysis activity; lysis pore formation |
| 19 | Uncharacterized secreted protein | 92 | 611247-611760  (+) | NO ANNOTATION | | SP | Secreted protein |
| 20 | Uncharacterized secreted protein | 91 | 649376-650210  (+) | NO ANNOTATION | | SP | Secreted protein |
| 21 | Uncharacterized protein | 715 | 667332-684888  (+) | NO ANNOTATION | | NO PREDICTION | |
| 22 | Uncharacterized membrane protein | 453 | 748389-756038  (+) | NO ANNOTATION | | TM | Membrane protein |
| 23 | alr1D | 424 | 701242-705191  (+) | gi\|302457159\|gb\|ADL39771.1\| allorecognition 1 [Hydractinia symbiolongicarpus] | 1.85E-99 | Ig-like fold X2; TM | Recognition; membrane receptor; protein-protein binding; alr1-like |
| 24 | TNF-like | 209 | 759992-763253  (+) | NO ANNOTATION | | TNF-like | Immune response; Receptor binding of tumor necrosis factor; protein-protein binding |
| 25 | Galactose-binding-like | 279 | 767425-771292  (+) | gi\|551548771\|ref\|XP_005762028.1\| regulator of telomere elongation helicase [Emiliania huxleyi CCMP1516] | 3.53E-08 | SP; ShKT; Galactose binding-like | Cell adhesion |
| 26 | Hydrolase-fold-containing | 322 | 795912-799197  (-) | gi\|449683348\|ref\|XP_004210331.1\| PREDICTED: uncharacterized protein LOC101241114 [Hydra vulgaris] | 2.55E-107 | SP; alpha/beta hydrolase fold | Lipid metabolism |
| 27 | MFS transporter-like | 628 | 804352-809728  (-) | gi\|449667661\|ref\|XP_002154002.2\| PREDICTED: monocarboxylate transporter 10-like [Hydra vulgaris] | 0.00 | MFS_general substrate transporter X2 | Transmembrane transport; integral membrane component |
| 28 | Uncharacterized membrane protein | 171 | 869896-876162  (-) | NO ANNOTATION | | SP; TM | Membrane protein |
| 29 | MFS transporter-like | 557 | 826181-830975  (-) | gi\|449686970\|ref\|XP_002162044.2\| PREDICTED: monocarboxylate transporter 10-like [Hydra vulgaris] | 3.20E-111 | MFS_general substrate transporter X2 | Transmembrane transport; integral membrane component |
| 30 | Glutamate receptor-like | 632 | 887674-893905  (+) | gi\|585651030\|ref\|XP_006814676.1\| PREDICTED: glutamate receptor ionotropic, NMDA 2A-like [Saccoglossus kowalevskii] | 2.10E-36 | Glutamate receptor L-glutamate/glycine binding; TM X2 | Transporter activity; ionotropic receptor activity; Membrane protein |
| 31 | Uncharacterized secreted protein | 170 | 1011435-1013370  (+) | NO ANNOTATION | | SP | Secreted protein |
| 32 | TNF-like | 93 | 1031640- 1032562  (+) | NO ANNOTATION | | TM; TNF-like | Immune response; Receptor binding of tumor necrosis factor; Membrane protein |
| 33 | Ig-likeD | 373 | 1037628-1047563  (+) | gi\|512842688\|ref\|XP_002931559.2\| PREDICTED: thrombospondin-2 [Xenopus (Silurana) tropicalis] | 4.36E-07 | Ig-like fold; TSP_1; TM | Protein binding; cell adhesion; Membrane protein |
| 34 | GPCR-like | 95 | 1051000-1051512  (-) | NO ANNOTATION | | GPCR, family 2, secretin-like; TM X3 | G-coupling receptor activity; Membrane protein |
| 35 | GPCR-like | 83 | 1052483-1053177  (-) | NO ANNOTATION | | GPCR, family 2, secretin-like; TM X3 | G-coupling receptor activity; Membrane protein |
| 36 | VWA typeA-containing | 265 | 990261-993615  (-) | gi\|449663000\|ref\|XP_004205667.1\| PREDICTED: collagen alpha-1(XII) chain-like [Hydra vulgaris] | 4.00E-20 | SP; VWA_type A | Ligand interaction; multi-protein complexes; cell adhesion, cell migration |
| 37 | LDL repeat-containing | 354 | 1054271-1057893  (-) | NO ANNOTATION | | LDL receptor class A repeat; TM | Protein-protein binding |
| 38 | Uncharacterized protein | 240 | 1166927-1170329  (+) | sp\|Q5PQJ7\|TBCEL_RAT Tubulin-specific chaperone cofactor E-like protein | 3.70E-63 | NO PREDICTION | |
| 39 | Ank repeat-containing | 380 | 1206304-1211123  (+) | gi\|528504032\|ref\|XP_694014.5\| PREDICTED: B-cell lymphoma 3 protein homolog [Danio rerio] | 4.55E-23 | Ank_repeat containing | Protein-protein binding |
| 40 | Uncharacterized protein | 410 | 1110102-1115351  (-) | gi\|156359434\|ref\|XP_001624774.1\| predicted protein [Nematostella vectensis] | 2.69E-72 | NO PREDICTION | |
| 41 | Zinc finger-containing | 1975 | 1083750-1108372  (-) | gi\|585683356\|ref\|XP_006812408.1\| PREDICTED: NFX1-type zinc finger-containing protein 1-like [Saccoglossus kowalevskii] | 0.00 | P-loop containing nucleoside triphosphate hydrolase; Zinc finger, NF-X1-type | DNA-binding transcription factor activity |
| 42 | Integrin | 790 | 1118049-1130726  (-) | gi\|10998792\|gb\|AAG25994.1\|AF308652_1 integrin beta chain [Podocoryna carnea] | 0.00 | Integrin beta subunit; TM | Receptor activity; Membrane protein |
| 43 | Polyketide synthase | 409 | 1133395-1141118  (-) | gi\|449692707\|ref\|XP_004213141.1\| PREDICTED: synaptic vesicle membrane protein VAT-1 homolog, partial [Hydra vulgaris] | 1.49E-119 | Polyketide synthase, enoylreductase | Redox activity; oxidoreductase activity, transferase |
| 44 | DnaJ-like | 496 | 1158020-1166145  (-) | gi\|449664494\|ref\|XP_002163923.2\| PREDICTED: dnaJ homolog subfamily C member 3-like [Hydra vulgaris] | 0.00 | SP; Tetratricopeptide-like; DnaJ | Protein-protein binding |
