## Supplementary material for "Multiple *alr* genes exhibit allorecognition-associated variation in the colonial cnidarian *Hydractinia*": Table S13

**Table S13. Variable proteins between individuals BC-3 and BC-15 outside ARC intervals**

| FUNCTIONAL GROUP | | # | PROTEIN | LENGTH | | HOMOLOGY ANNOTATION | | FUNCTIONAL ANNOTATION | | P-DISTANCE |
| --- | --- | --- | --- | --- | --- | --- | --- | --- | --- | --- |
|  |  |  |  | **BC-3 (aa)** | **BC-15 (aa)** | **DESCRIPTION** | **E-VALUE** | **DOMAIN ARCHITECTURE** | **FUNCTIONAL DESCRIPTION** |  |
| IMMUNITY, RECOGNITION AND CELLULAR ADHESION | 1 | | CAP-containing | 351 | 284 | gi\|302833369\|ref\|XP_002948248.1\| PR-1 like protein [Volvox carteri f. nagariensis] | 6.21E-18 | CAP domain | Cellular adhesion, extracellular matrix regulation, branching morphogenesis; proteases, protease inhibition; secreted | 0.59 |
|  | 2 | | TNF-like | 268 | 294 | gi\|449689722\|ref\|XP_002170861.2\| PREDICTED: uncharacterized protein LOC100215981, partial [Hydra vulgaris] | 3.68E-12 | TM; TNF-like | Immune response; tumor necrosis factor receptor binding; protein-protein interaction; membrane protein | 0.53 |
|  | 3 | | von Willebrand factor D and EGF-containing | 745 | 1214 | gi\|449676184\|ref\|XP_002158680.2\| PREDICTED: von Willebrand factor D and EGF domain-containing protein-like [Hydra vulgaris] | 2.61E-101 | Nidogen, extracellular domain ; Ig-like fold; von Willebrand factor, type D domain; GPS motif; GPCR, family 2-like (TM X4) | Matrix cellular adhesion; protein-protein interaction; signaling pathway; transmembrane receptor | 0.50 |
|  | 4 | | TIR receptor homology | 385 | 298 | gi\|156383902\|ref\|XP_001633071.1\| predicted protein [Nematostella vectensis] | 3.61E-35 | Toll/interleukin-1 receptor homology (TIR) | Signaling transduction; protein-protein interaction | 0.37 |
|  | 5 | | EGF-like | 301 | 181 | gi\|449677846\|ref\|XP_002170373.2\| PREDICTED: laminin subunit alpha-like, partial [Hydra vulgaris] | 4.16E-45 | EGF-like, laminin; Laminin B type IV | Cellular adhesion, Migration, proliferation, cellular signaling and proliferation | 0.25 |
|  | 6 | | Fibrinogen-containing | 786 | 392 | gi\|530567272\|ref\|XP_005280161.1\| PREDICTED: ficolin-2-like [Chrysemys picta bellii] | 7.71E-43 | SP; Fibrinogen, alpha/beta/gamma chain, C-terminal globular domain | Immune response; adhesion | 0.19 |
|  | 7 | | MORN-containing | 233 | 175 | gi\|632970624\|ref\|XP_007901754.1\| PREDICTED: radial spoke head 1 homolog [Callorhinchus milii] | 1.01E-38 | MORN (Membrane Occupation and Recognition Nexus) motifs | Motif found in multiple copies in various proteins involved in cellular adhesion and cell binding | 0.13 |
|  | 8 | | CD20-like | 183 | 225 | No annotation | CD20-like | N/A | Integral membrane component; receptor | 0.14 |
|  | 9 | | Kringle and EGF-like | 374 | 226 | gi\|449687116\|ref\|XP_002165254.2\| PREDICTED: uncharacterized protein LOC100198726, partial [Hydra vulgaris] | 1.44E-06 | EGF-like; Kringle-like | Cellular adhesion; protein-protein interaction | 0.05 |
|  | 10 | | EGF-like | 860 | 1191 | gi\|449683277\|ref\|XP_002155677.2\| PREDICTED: neurogenic locus Notch protein-like, partial [Hydra vulgaris] | 0.00 | SP; EGF-like X14 | Cellular adhesion; protein-protein interaction; Calcium-binding | 0.06 |
|  | 11 | | EGF-like | 1474 | 1699 | sp\|P07942\|LAMB1_HUMAN Laminin subunit beta-1 OS=Homo sapiens GN=LAMB1 PE=1 SV=2 | 8.95E-115 | SP; Laminin, N-terminal; EGF-like, laminin X7 | Protein-protein interaction; Cellular adhesion | 0.03 |
|  | 12 | | Nidogen / Kazal / MAM-containing | 445 | 232 | gi\|410903396\|ref\|XP_003965179.1\| PREDICTED: MAM domain-containing protein 2-like [Takifugu rubripes] | 7.86E-17 | Nidogen; Kazal; MAM domain | Matrix cellular adhesion; protein-protein interaction; membrane; protease inhibition | 0.02 |
|  | 13 | | C-type lectin-fold and EGF-like-containing | 3282 | 3346 | sp\|P22897\|MRC1_HUMAN Macrophage mannose receptor 1 OS=Homo sapiens GN=MRC1 PE=1 SV=1 | 7.70E-35 | SP; C-type lectin fold X2; Complement Clr-like EGF domain; C-type lectin fold X3; EGF-like; TM | Protein-protein interaction; Calcium-binding; Carbohydrate-binding | 0.02 |
|  | 14 | | Frizzled domain and Kringle-like-containing | 657 | 673 | gi\|340373142\|ref\|XP_003385101.1\| PREDICTED: tyrosine-protein kinase transmembrane receptor ROR1-like [Amphimedon queenslandica] | 7.23E-20 | Frizzled domain X2; TM; Kringle-like domain; TM | Protein-protein interaction; Cellular adhesion | 0.02 |
|  | 15 | | VWA type A-containing | 1174 | 2582 | sp\|A6NMZ7\|CO6A6_HUMAN Collagen alpha-6(VI) chain OS=Homo sapiens GN=COL6A6 PE=1 SV=2 | 1.94E-23 | von Willebrand factor, type A X4 | Protein-protein interaction; Ligand-binding; Cellular adhesion, Migration | 0.01 |
|  | 16 | | Sushi / EGF-like / Ig-containing | 1855 | 887 | sp\|A1XQY1\|NR3BA_DANRE Neurexin-3b OS=Danio rerio GN=nrxn3b PE=2 SV=1 | 8.29E-06 | Sushi; EGF-like; Ig subtype | Protein-protein interaction; Cellular adhesion | 0.01 |
|  | 17 | | EGF-like | 1633 | 1549 | sp\|A0JP86\|LAMC1_XENTR Laminin subunit gamma-1 OS=Xenopus tropicalis GN=lamc1 PE=2 SV=1 | 0.00 | SP; Laminin, N-terminal; EGF-like, laminin X6 | Protein-protein interaction; Cellular adhesion | 0.01 |
|  | 18 | | Cadherin-like | 2326 | 2123 | gi\|386118337\|gb\|AFI99116.1\| seven transmembrane protocadherin flamingo [Clytia hemisphaerica] | 0.00 | Cadherin-like; EGF-like; Concanavalin A-like lectin/glucanases superfamily X 2; EGF-like; GPS motif | Cellular adhesion; protein-protein interaction; G-protein receptor-coupled activity; membrane protein | 0.001 |
|  | 19 | | MORN-containing | 1133 | 1072 | sp\|Q8WXH2\|JPH3_HUMAN Junctophilin-3 OS=Homo sapiens GN=JPH3 PE=1 SV=2 | 5.81E-58 | MORN (Membrane Occupation and Recognition Nexus) motifs | Motif found in multiple copies in various proteins involved in cellular adhesion and cell binding | 0.001 |
|  | 20 | | EGF-like | 1025 | 991 | gi\|449678542\|ref\|XP_002160645.2\| PREDICTED: laminin subunit alpha-like [Hydra vulgaris] | 1.12E-82 | SP; EGF-like, laminin X3 | Cellular adhesion; secreted protein | 0.001 |
|  | 21 | | Cadherin-like | 4451 | 4174 | gi\|156375132\|ref\|XP_001629936.1\| predicted protein [Nematostella vectensis] | 3.05E-44 | Cadherin-like X6; Ig-like fold X3; TM | Cytoskeleton anchorage; protein-protein interaction; Calcium-binding; dystrophin-associated glycoprotein complex | 0.001 |
|  | 22 | | TSP_1 repeat-containing | 348 | 603 | sp\|Q96RW7\|HMCN1_HUMAN Hemicentin-1 OS=Homo sapiens GN=HMCN1 PE=1 SV=2 | 9.59E-33 | Ig-like fold; Thrombospondin, type 1 repeats; Ig-like fold | Protein-protein interaction | 0.001 |
|  | 23 | | Ig-like | 241 | 241 | gi\|449668297\|ref\|XP_002153999.2\| PREDICTED: receptor-type tyrosine-protein phosphatase delta-like [Hydra vulgaris] | 1.21E-38 | SP; Ig-like fold X2 | Recognition; protein-protein interaction | 0.001 |
|  | 24 | | DOMON-containing | 391 | 407 | No annotation | | Dopamine beta-Monooxygenase N-terminal (DOMON) domain | Extracellular adhesion; transmembrane protein | 0.04 |
| PROTEIN-PROTEIN INTERACTION | 25 | | Tetratricopeptide-like-containing | 314 | 565 | sp\|Q8BMA6\|SRP68_MOUSE Signal recognition particle subunit SRP68 OS=Mus musculus GN=Srp68 PE=2 SV=2 | 1.01E-162 | Tetratricopeptide-like helical domain X2 | Protein-protein interaction; protein targeting to membrane; signal recognition particle binding | 0.08 |
|  | 26 | | Tetratricopeptide-like | 2227 | 1972 | gi\|449679122\|ref\|XP_002168350.2\| PREDICTED: tetratricopeptide repeat protein 28-like, partial [Hydra vulgaris] | 0.00 | Tetratricopeptide-like helical domain X4; CHAT domain | Mitotic cycle regulation; protein-protein interaction | 0.001 |
|  | 27 | | Pleckstrin homology-like-containing | 587 | 535 | gi\|555695472\|gb\|ESN98704.1\| hypothetical protein HELRODRAFT_162157 [Helobdella robusta] | 3.93E-06 | Pleckstrin homology-like domain; SH2 domain | Protein-protein interaction | 0.01 |
|  | 28 | | PKD / REJ-like-containing | 2102 | 2497 | gi\|585652993\|ref\|XP_006815028.1\| PREDICTED: polycystic kidney disease protein 1-like 2-like [Saccoglossus kowalevskii] | 3.57E-151 | PKD domain X2 (Ig-like fold); Egg jelly receptor, REJ-like; GPS motif; Lipase/lipoxygenase, PLAT/LH2 | Protein-protein interaction; protein-protein or protein-carbohydrate adhesion; signaling | 0.01 |
|  | 29 | | WD40/YVTN repeat-like-containing | 423 | 672 | gi\|585696548\|ref\|XP_006822221.1\| PREDICTED: WD repeat domain phosphoinositide-interacting protein 2-like isoform X2 [Saccoglossus kowalevskii] | 3.62E-153 | WD40/YVTN repeat-like-containing domain X3 | Protein-protein interaction | 0.05 |
|  | 30 | | Tetratricopeptide-like | 559 | 472 | gi\|156367315\|ref\|XP_001627363.1\| predicted protein [Nematostella vectensis] | 8.78E-39 | Tetratricopeptide-like helical domain X2 | Protein-protein interaction | 0.01 |
|  | 31 | | F-box / LRR-containing | 721 | 746 | gi\|291241443\|ref\|XP_002740625.1\| PREDICTED: F-box/LRR-repeat protein 7-like [Saccoglossus kowalevskii] | 5.81E-80 | F-box domain; Leucine-rich repeats, cysteine-containing subtype | Protein-protein interaction | 0.01 |
|  | 32 | | WD40/YVTN repeat-like-containing | 905 | 417 | gi\|156383902\|ref\|XP_001633071.1\| predicted protein [Nematostella vectensis] | 3.61E-35 | SP; WD40/YVTN repeat-like-containing domain; Plexin; Thrombospondin, type 1 repeats; TM | Protein-protein interaction; development; receptor activity; membrane receptor | 0.001 |
|  | 33 | | Ankyrin repeat-containing | 1326 | 1190 | SIN ANOTACIÓN | | P-loop containing nucleoside triphosphate hydrolase; Ig-like fold; Ankyrin repeat-containing domain | Protein-protein interaction | 0.45 |
|  | 34 | | Calponin homology | 1212 | 1218 | gi\|449673152\|ref\|XP_002163420.2\| PREDICTED: uncharacterized protein LOC100208415 [Hydra vulgaris] | 2.95E-112 | Calponin homology domain | Protein-protein interaction | 0.01 |
|  | 35 | | GYF domain-containing | 753 | 711 | gi\|449668186\|ref\|XP_004206732.1\| PREDICTED: PERQ amino acid-rich with GYF domain-containing protein 2-like [Hydra vulgaris] | 8.20E-96 | glycine-tyrosine-phenylalanine (GYF) domain | Protein-protein interaction; ligand-binding; recognition of proline-rich sequences | 0.001 |
| PROTEOLYSIS | 36 | | Peptidase | 443 | 251 | gi\|85057108\|emb\|CAJ57450.1\| astacin 4 [Hydractinia echinata] | 3.82E-64 | Peptidase, metallopeptidase | Proteolysis; metalloendopeptidase activity; Zinc-binding | 0.30 |
|  | 37 | | Peptidase M12-like | 234 | 894 | sp\|P28826\|MEP1B_RAT Meprin A subunit beta OS=Rattus norvegicus GN=Mep1b PE=1 SV=3 | 9.39E-26 | TM; Peptidase M12A, astacin; Concanavalin A-like lectin; Peptidase, metallopeptidase; Concanavalin A-like lectin | Proteolysis; metallopeptidase activity; Zinc-binding | 0.01 |
|  | 38 | | Peptidase M2-like | 1323 | 1323 | gi\|449662193\|ref\|XP_002165887.2\| PREDICTED: uncharacterized protein LOC100206770 [Hydra vulgaris] | 0.00 | SP; Peptidase M2, peptidyl-dipeptidase A | Proteolysis; metallopeptidase activity; peptidyl-dipeptidase activity | 0.001 |
| ENZYMES | 39 | | Thiolase-like | 357 | 419 | sp\|P32020\|NLTP_MOUSE Non-specific lipid-transfer protein OS=Mus musculus GN=Scp2 PE=1 SV=3 | 3.34E-144 | Thiolase-like X2 | Metabolic process; catalytic activity; acyl-group transferase activity | 0.52 |
|  | 40 | | MBOAT-like | 402 | 150 | gi\|449681695\|ref\|XP_004209897.1\| PREDICTED: lysophospholipid acyltransferase 5-like [Hydra vulgaris] | 1.35E-43 | Membrane bound O-acyl transferase, MBOAT | Acyltransferase enzyme | 0.46 |
|  | 41 | | Glutathione S-transferase-like | 271 | 228 | gi\|221103804\|ref\|XP_002158423.1\| PREDICTED: glutathione S-transferase-like [Hydra vulgaris] | 2.39E-21 | Thioredoxin-like fold; Glutathione S-transferase, C-terminal-like | Oxidation-reduction process; oxidoreductase activity, transferase | 0.20 |
|  | 42 | | Ferric reductase, NAD-binding-like | 244 | 483 | gi\|449679778\|ref\|XP_002157071.2\| PREDICTED: NADPH oxidase 3-like, partial [Hydra vulgaris] | 9.87E-68 | Ferric reductase transmembrane component-like domain; Ferric reductase, NAD binding (TM X3) | Oxidation-reduction process; oxidoreductase activity | 0.37 |
|  | 43 | | Thioredoxin-like-fold-containing | 377 | 316 | sp\|Q09652\|GSTK1_CAEEL Glutathione S-transferase kappa 1 OS=Caenorhabditis elegans GN=gstk-1 PE=3 SV=1 | 9.08E-06 | Thioredoxin-like fold X2 | Disulfide bond oxidoreductase activity | 0.03 |
|  | 44 | | Pyridoxal phosphate-dependent transferase | 933 | 627 | gi\|449680764\|ref\|XP_002168228.2\| PREDICTED: uncharacterized protein LOC100197171, partial [Hydra vulgaris] | 0.00 | Pyridoxal phosphate-dependent transferase X2 | Carboxylic Acid Metabolic Process; catalytic activity | 0.37 |
|  | 45 | | Hydrolase fold-containing | 221 | 223 | gi\|449683921\|ref\|XP_004210498.1\| PREDICTED: ovarian cancer-associated gene 2 protein homolog, partial [Hydra vulgaris] | 5.04E-58 | Alpha/Beta hydrolase fold | Hydrolytic enzyme | 0.10 |
|  | 46 | | GTP-binding hydrolase | 549 | 549 | sp\|Q6DCC6\|GTPB6_XENLA Putative GTP-binding protein 6 OS=Xenopus laevis GN=gtpbp6 PE=2 SV=1 | 4.23E-106 | GTPase HflX N-terminal domain; P-loop containing nucleoside triphosphate hydrolase | GTP-binding; hydrolase activity | 0.02 |
| PROTEASE INHIBITION | 47 | | Thyroglobulin type 1 and fibrinogen-containing | 2279 | 3315 | gi\|449671029\|ref\|XP_002162557.2\| PREDICTED: uncharacterized protein LOC100202739 [Hydra vulgaris] | 0.00 | Thyroglobulin type-1 X2; Concanavalin A-like lectin; Fibrinogen | Serine endopeptidase inhibitory activity | 0.01 |
|  | 48 | | Kazal-containing | 751 | 633 | SIN ANOTACIÓN | | SP; Kazal domain; TM | Protein-protein interaction; protease inhibition | 0.02 |
| TRANSPORT | 49 | | Band 7-Stomatin | 329 | 230 | gi\|449683403\|ref\|XP_002161494.2\| PREDICTED: mechanosensory protein 2-like isoform 1 [Hydra vulgaris] | 1.29E-133 | TM; Band 7 protein - Stomatin family | Regulation of cation transport in mechanosensor proteins; membrane protein | 0.13 |
|  | 50 | | Neurotransmitter-gated ion channel | 292 | 405 | sp\|P57695\|GLRA1_BOVIN Glycine receptor subunit alpha-1 OS=Bos taurus GN=GLRA1 PE=2 SV=1 | 2.42E-39 | Neurotransmitter-gated ion-channel ligand-binding; Neurotransmitter-gated ion-channel transmembrane domain (TM X4) | Ion transport; integral membrane component; extracellular ligand-gated ion channel | 0.51 |
|  | 51 | | Ion transport domain-containing | 900 | 895 | gi\|221122827\|ref\|XP_002153926.1\| PREDICTED: galactose-3-O-sulfotransferase 3-like [Hydra vulgaris] | 0.00 | Ion transport domain; P-loop containing nucleoside triphosphate hydrolase | Ion transport; transmembrane transport; ion channel activity | 0.01 |
| UNRELATED FUNCTIONS | 52 | | Concanavalin A-like lectin | 444 | 498 | sp\|P97927\|LAMA4_MOUSE Laminin subunit alpha-4 OS=Mus musculus GN=Lama4 PE=1 SV=2 | 5.99E-10 | Concanavalin A-like lectin/glucanases superfamily X2 | Carbohydrate-binding | 0.001 |
|  | 53 | | Ras guanine nucleotide exchange factor | 1021 | 1019 | sp\|Q552M5\|GEFY_DICDI Ras guanine nucleotide exchange factor Y OS=Dictyostelium discoideum GN=gefY PE=1 SV=2 | 1.38E-28 | Ras guanine nucleotide exchange factor | GTPase-mediated signal transduction; guanyl-nucleotide exchange factor activity | 0.001 |
|  | 54 | | EF-hand-containing | 189 | 162 | gi\|449674275\|ref\|XP_002169270.2\| PREDICTED: programmed cell death protein 6-like [Hydra vulgaris] | 2.87E-87 | EF-hand domain X3 | Calcium-binding | 0.001 |
|  | 55 | | Prominin | 854 | 742 | gi\|156388972\|ref\|XP_001634766.1\| predicted protein [Nematostella vectensis] | 6.29E-14 | SP; Prominin protein family (TM X 4) | Integral membrane component | 0.02 |
|  | 56 | | Zinc finger-containing | 425 | 438 | gi\|40019235\|emb\|CAE88928.1\| TNF-receptor-associated factor 1 [Hydractinia echinata] | 0.00 | Zinc finger, RING/FYVE/PHD-type; Zinc finger, TRAF-type X2; | Protein ubiquitination; activation of NF-kappaB-inducing kinase; signal transduction activity | 0.03 |
|  | 57 | | Small GTP-binding | 191 | 190 | gi\|221116683\|ref\|XP_002158815.1\| PREDICTED: ras-related C3 botulinum toxin substrate 1-like [Hydra vulgaris] | 1.26E-130 | Small GTP-binding protein domain | GTP catabolic process; GTPase-mediated signal transduction; protein transport | 0.05 |
|  | 58 | | Sterol regulatory element-binding | 740 | 863 | sp\|A0JPH4\|SCAP_XENLA Sterol regulatory element-binding protein cleavage-activating protein OS=Xenopus laevis GN=scap PE=2 SV=1 | 4.38E-31 | Sterol-sensing domain (TM X3); TM X2; WD40/YVTN repeat-like-containing | Protein-protein interaction; integral membrane component | 0.04 |
|  | 59 | | Notch-like | 770 | 1084 | gi\|156401699\|ref\|XP_001639428.1\| predicted protein [Nematostella vectensis] | 6.7E-12 | SP; Delta/Serrate/lag-2 (DSL) protein; TM; Delta/Serrate/lag-2 (DSL) protein; TM | Cellular communication; membrane protein | 0.01 |
|  | 60 | | Histone-fold-containing | 399 | 508 | gi\|449663064\|ref\|XP_004205676.1\| PREDICTED: uncharacterized protein LOC101238105 [Hydra vulgaris] | 3.96E-23 | Domain of unknown function DUF4477; Histone fold | DNA-binding | 0.15 |
|  | 61 | | Tyrosine-protein phosphatase non-receptor | 1191 | 1737 | gi\|533176166\|ref\|XP_005401411.1\| PREDICTED: tyrosine-protein phosphatase non-receptor type 13 isoform X9 [Chinchilla lanigera] | 1.47E-55 | FERM domain; PDZ domain X5; Protein-tyrosine phosphatase-like | Protein dephosphorylation; tyrosine phosphatase activity; protein-protein interaction | 0.01 |
|  | 62 | | HSP20-like chaperone | 265 | 279 | gi\|449670958\|ref\|XP_004207394.1\| PREDICTED: uncharacterized protein LOC100204018 [Hydra vulgaris] | 6.70E-76 | HSP20-like chaperone | Chaperone protein | 0.49 |
|  | 63 | | BRCT-containing | 502 | 495 | gi\|551508250\|ref\|XP_005805654.1\| PREDICTED: uncharacterized protein C15orf43 homolog [Xiphophorus maculatus] | 1.62E-07 | BRCT domain (C_terminal domain of a breast cancer susceptibility protein) | Involved in cell cycle checkup functions in response to DNA damage | 0.15 |
|  | 64 | | Endonuclease-like | 390 | 300 | gi\|449675419\|ref\|XP_002166096.2\| PREDICTED: endonuclease 8-like 1-like [Hydra vulgaris] | 8.09E-55 | DNA glycosylase/AP lyase, catalytic domain; Endonuclease VIII-like 1, DNA binding | DNA-repair; DNA-binding; DNA lyase activity; Zinc-binding | 0.15 |
|  | 65 | | Importin subunit alpha-2-like | 481 | 514 | gi\|221128609\|ref\|XP_002165850.1\| PREDICTED: importin subunit alpha-2-like [Hydra vulgaris] | 0.00 | Importin-alpha-beta-binding domain; armadillo repeats | Protein import into the nucleus; protein-protein interaction; transporter activity; mediation of protein-protein interactions | 0.52 |
|  | 66 | | Rap GTPase activating | 881 | 509 | gi\|449663035\|ref\|XP_002157783.2\| PREDICTED: rap1 GTPase-activating protein 1-like [Hydra vulgaris] | 6.31E-89 | Zinc finger, LIM-type; Rap GTPase activating protein; BTB/POZ fold | Regulation of signal transduction by GTPases; GTPase activation; Zinc-binding | 0.001 |
|  | 67 | | TSP_1 repeat-containing | 80 | 337 | sp\|Q3UHD1\|BAI1_MOUSE Brain-specific angiogenesis inhibitor 1 OS=Mus musculus GN=Bai1 PE=1 SV=1 | 1.43E-21 | Thrombospondin, type 1 repeats; TM | Cell-cell interaction; inhibition of angiogenesis and apoptosis | 0.37 |
|  | 68 | | MARVEL-containing | 173 | 205 | No annotation | | MARVEL domain (TM X4) | Membrane component; vesicular trafficking; membrane binding | 0.24 |
| UNCHARACTERIZATED FUNCTIONS | 69 | | No annotation | 2536 | 2536 | No annotation | | SP | Secreted protein | 0.001 |
|  | 70 | | No annotation | 815 | 724 | gi\|156395754\|ref\|XP_001637275.1\| predicted protein [Nematostella vectensis] | 8.97E-65 | SP | Secreted protein | 0.01 |
|  | 71 | | No annotation | 357 | 411 | gi\|221129269\|ref\|XP_002157161.1\| PREDICTED: sorting nexin-7-like [Hydra vulgaris] | 9.04E-114 | TM | Transmembrane protein | 0.14 |
|  | 72 | | No annotation | 1038 | 1752 | gi\|449689390\|ref\|XP_002159385.2\| PREDICTED: uncharacterized protein LOC100210512, partial [Hydra vulgaris] | 9.93E-85 | SP | Secreted protein | 0.43 |
|  | 73 | | No annotation | 464 | 321 | No annotation | | TM X2 | Transmembrane protein | 0.11 |
|  | 74 | | No annotation | 685 | 558 | gi\|449672170\|ref\|XP_004207650.1\| PREDICTED: uncharacterized protein LOC101240082 [Hydra vulgaris] | 4.68E-20 | SP | Secreted protein | 0.02 |
|  | 75 | | No annotation | 343 | 328 | gi\|383458753\|ref\|YP_005372742.1\| FG-GAP repeat-containing protein [Corallococcus coralloides DSM 2259] | 9.28E-36 | SP; TM | Transmembrane protein | 0.05 |
|  | 76 | | No annotation | 230 | 230 | gi\|449676298\|ref\|XP_002167064.2\| PREDICTED: uncharacterized protein LOC100215046 [Hydra vulgaris] | 2.84E-10 | TM X4 | Transmembrane protein | 0.02 |
|  | 77 | | No annotation | 353 | 279 | No annotation | | SP; TM | Transmembrane protein | 0.001 |
|  | 78 | | No annotation | 727 | 727 | gi\|449679518\|ref\|XP_002165926.2\| PREDICTED: uncharacterized protein LOC100207090 [Hydra vulgaris] | 0.00 | Protein of unknown function DUF1399 | Uncharacterized | 0.03 |
